# A generalizable interface-seeded framework for *de novo* design of functional oligomers

**DOI:** 10.64898/2026.07.02.736098

**Authors:** Ho Yeung Chim, Mubarak Olanrewaju Idris, Dominic Rieger, Phillip Schlegel, Nicolas Manuel Goldbach, Miguel Atienza Juanatey, Bhoomika Basu Mallik, Stephen Buckley, Samrat Basak, Sandrine Georgeon, Kelvin Lau, Florence Pojer, Leonard Kaysser, Philip Tinnefeld, Clara T. Schoeder, Bruno E. Correia, Alena Khmelinskaia

## Abstract

Protein oligomers are ubiquitous in biological systems and essential for function. However, the de novo design of oligomers that controllably assemble in response to exogenous stimuli remains challenging. Here, we present an AI-based generative approach that leverages an interface-seeded strategy for designing responsive homo-oligomers from isolated interaction modules. Experimentally validated designs are highly accurate and explore new-to-nature topologies. We show that designs effectively respond to their chemical triggers with conditional oligomerization or to phosphorylation-driven conformational changes with reversible oligomerization. We further functionalized our responsive assemblies to build ligand-dependent membrane binding systems and phosphorylation-controlled gene regulatory switches. Our framework enables the generalizable design of responsive protein complexes, opening novel possibilities for the engineering of biosynthetic systems with sophisticated regulatory mechanisms.

## Main Text

The field of protein design has advanced rapidly in recent years and stands as a fundamental tool for the generation of novel proteins, enabling faster and more efficient exploration of protein structural and functional landscapes than natural evolution. Although the field has extensively explored the design of unconditional protein assemblies (*1–6*), programming proteins with controlled assembly processes remains challenging, hindering their effective integration with biological systems (*7*). In contrast, natural proteins dynamically respond to environmental cues and can undergo changes in oligomeric states to transduce biochemical information, dynamically regulating cellular processes essential for life (*8–10*). The formation of assemblies composed of more than two subunits enables the amplification of weak input signals by concentrating active signaling components into organized higher-order molecular complexes. Such assemblies can generate rapid, switch-like responses, enhance signaling efficiency and robustness by sequestering active intermediates from negative regulators (*11*). Many of these systems reversibly transition between inactive monomeric states and active oligomeric states, allowing the rapid and dynamic transduction of signals in response to stimuli (*12*). Mastering the design of responsive assemblies could unlock a broad range of applications in the development of biologics, synthetic protein networks, biosensing and nanocomputing.

Tailored strategies for designing environmentally responsive assemblies have been developed, using pH (*13–15*), redox potential (*16*), light (*17*), an effector peptide (*18*), or a small molecule (*19*) as triggers. Most rely either on empirically built chemical intuition or a “docking-while-binding” approach combined with symmetric constraints. However, neither strategy is easily generalizable to different classes of stimuli. A method that allows versatile, combinatorial construction of complex architectures with predefined geometries using a validated set of responsive motifs would provide significant advantages in modularity, predictability, and reconfigurability.

We thus sought to establish a general protocol for designing programmable protein architectures from pre-validated interaction interfaces (Fig. 1). We reasoned that reusing experimentally verified protein-protein interfaces (PPIs) would offer an efficient path towards modular and programmable assembly design. We use this concept to design oligomers with diverse levels of functional complexity, from obligate oligomers, to oligomers induced by metal- or small molecule-mediated interactions or reversibly controlled by a phosphorylation-stabilized conformational change. We further show that the designed assemblies can be used to engineer oligomerization-controlled signaling mechanisms in biological contexts, opening new avenues to build artificial pathways featuring regulatory mechanisms akin to those found in nature.

**Fig. 1.**
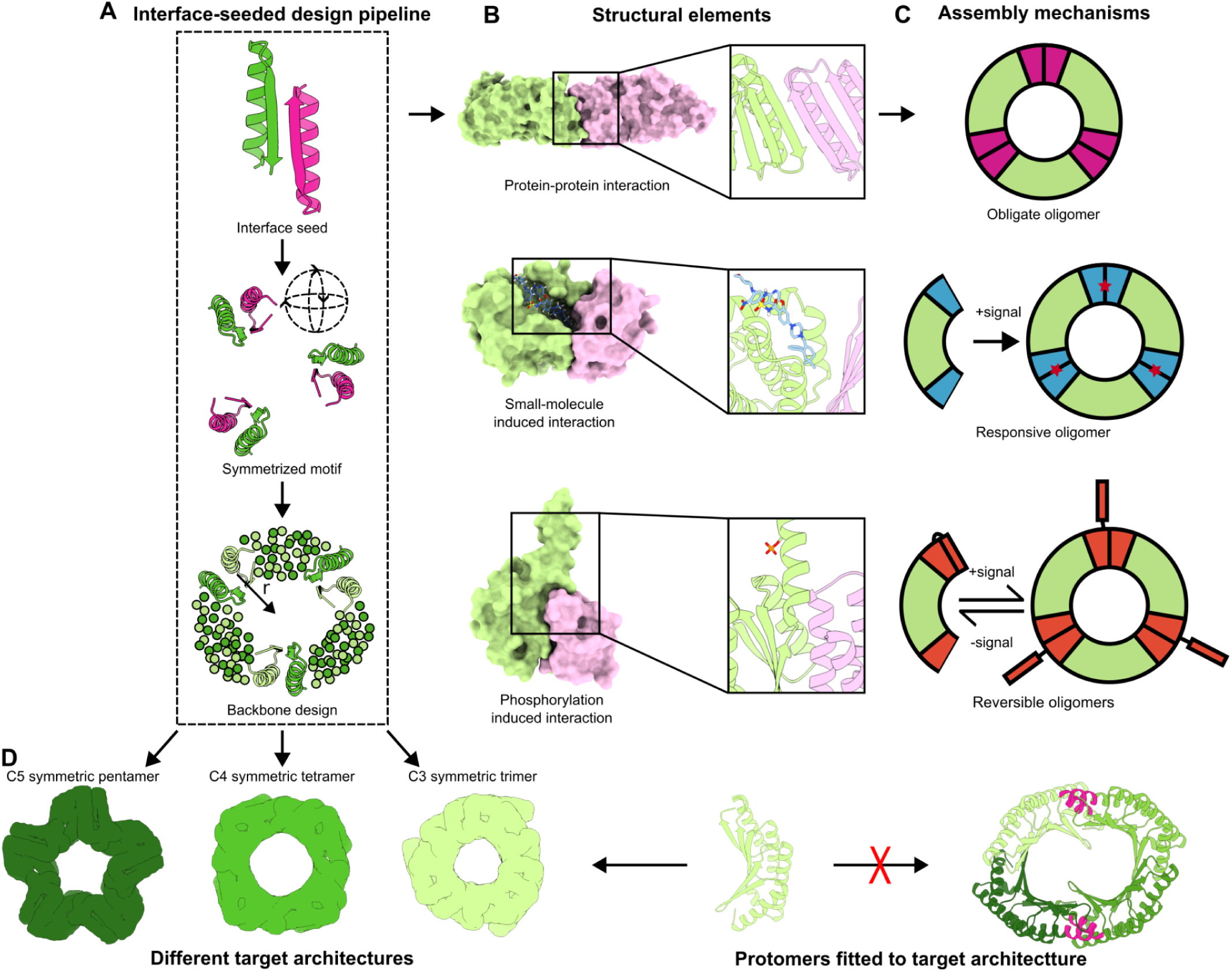
Interface-seeded design of functional symmetric oligomers. (**A**) Schematic representation of our interface-seeded approach for the design of protomer shapes tailored to the desired symmetry from interface solely. Starting from a known interaction mechanism, the interface is extracted and symmetrized according to the target geometry (here a C3 trimer). At this step, its relative position is randomized by sampling both the rotational and translational degrees of freedom. During the denoising trajectory, the radial position of the motif is adjusted, guided by the center of mass of the generated scaffolds, biasing generation towards well packed backbones. (**B**) Examples of structural elements explored as seeds for (**C**) design of functional oligomers with different assembly mechanisms. (**D**) The design pipeline can be used for the design of any target cyclic oligomer and generates protomers specifically fitted to stabilize the interface and match the shape requirements of the target architecture.

### Computational design approach

We first evaluated the state-of-the-art dock-and-design approach (*6*, *20*) (Fig. S1) for generating symmetric oligomers from a limited heterodimer set. We chose to dock the well-characterized and broadly used heterodimer LHD101 (*18*, *21–26*) into C3, C4, and C5 symmetric assemblies. LHDs provide an attractive testbed because of their stable monomeric state in isolation and polar, high-affinity PPIs, which rely on shape complementarity and precisely designed interfacial hydrogen bonds (*21*). Their successful use across protein-design applications makes them reliable benchmarks for evaluating design strategies (*18*, *21–26*). We observed that docking configurations in which higher order oligomerization can occur only if both heterodimer subunits are present, could be generated only for a single symmetry class (Fig. S1). To increase protomer diversity, we generated a dimer library by motif scaffolding the LHD101 interface prior to docking, but observed similar bias for a single type of symmetric oligomer and poor overall success rate, as judged by the residue-pair transform (rpx) score and buried solvent-accessible surface area (SASA) reflecting the quality of the created interfaces. Such observations across different heterodimer sets (Fig. S1), reinforce our prior intuition that docking-based assembly design is constrained by the compatibility between protomer geometry and target oligomer symmetry.

To address this challenge, we developed an interface-seeded design strategy in which a validated PPI can be used to generate building-blocks compatible with a target symmetry (Fig. 1). While docking is constrained by the inherent shapes and surface properties of the chosen protein building blocks, generating such chimeric building blocks would simultaneously satisfy the assembly PPI and match the internal geometry of the target architecture. We implemented a new symmetric motif-scaffolding module in RFdiffusion to generate diverse, compact symmetry-compatible backbones that preserve (functional) interface motifs (Fig. 1A). To avoid manual tuning of the relative orientation of the motifs within the target geometry and increase the diversity of generated backbones, the motif is randomly rotated prior to initialization of the diffusion trajectory within a user-specified range of radial distances. During the diffusion process, the radial position of the motif is guided by the center of mass of the denoised residues in each step. Sequences for backbones passing structural filters (see Materials and Methods) were designed using SolubleMPNN (*27*) while preserving motif residues, and filtered using AlphaFold2 (AF2) (*28*, *29*) alone or complemented with AlphaFold3 (AF3) (*30*). We experimentally validated this framework across multiple PPIs of diverse origins and with different environmental response mechanisms (Fig. 1B,C), demonstrating its generality and adaptability (Fig. 1D).

### Benchmark on *de novo* obligate protein-protein interaction seeds

We first evaluated our interface-seeded approach by designing obligate cyclic assemblies of varying symmetry order (C3, C4 and C5 symmetric trimers, tetramers and pentamers, respectively) using two LHD heterodimeric, obligate PPIs as binding modules (Fig. 2A), i.e. LHD29 and LHD101.

**Fig. 2.**
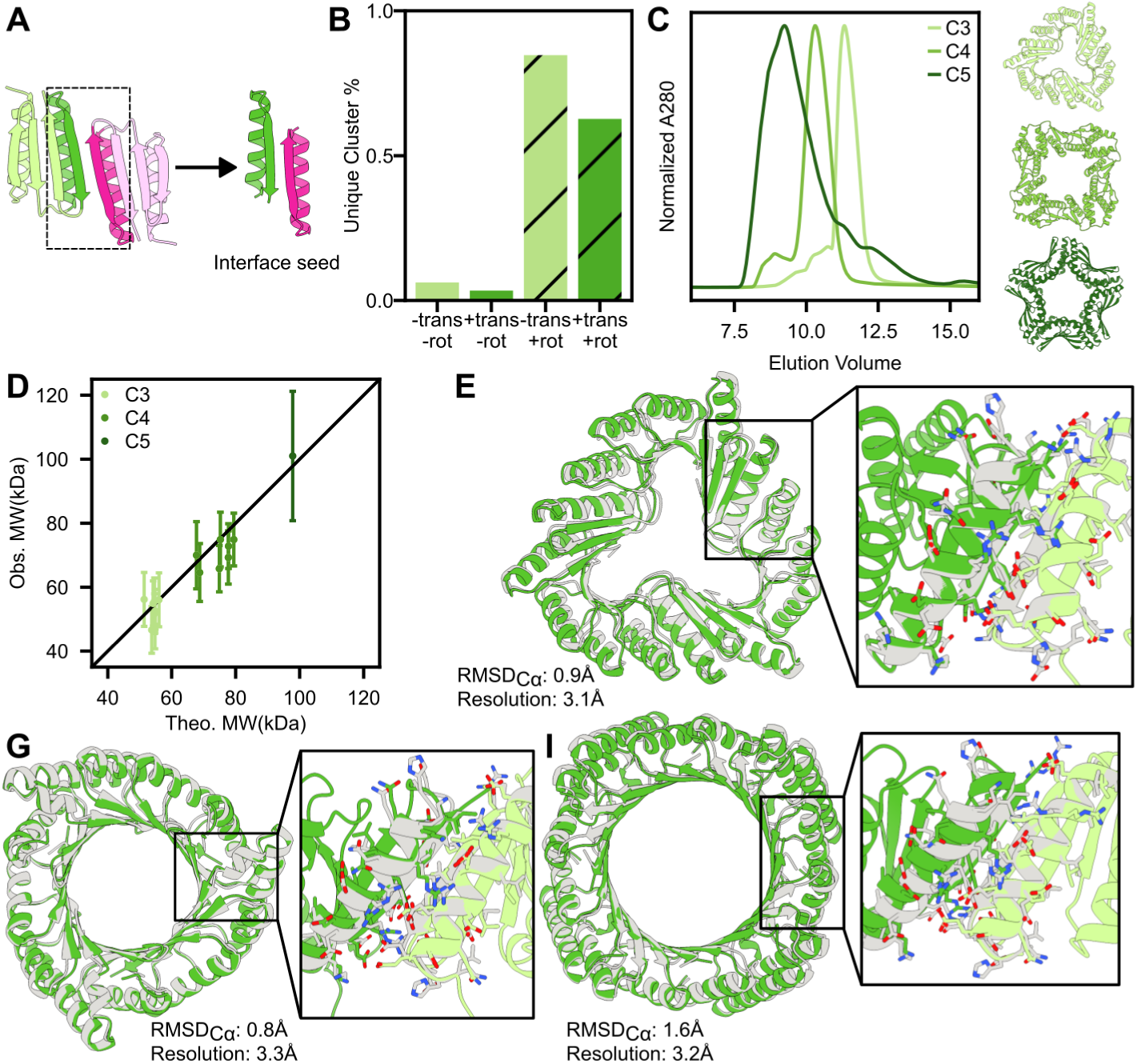
*De novo* design of protein oligomers with diverse cyclic symmetry order from a single interface seed. (**A**) Interface seed extracted from the LHD101 heterodimer (PDB ID 7MWR). (**B**) Generated design diversity increases when sampling the orientation (rotational degrees of freedom, rot) of the interface seed, while the applied dragging force (along the translational degree of freedom, trans) constrains the final oligomer radius, slightly reducing diversity. (**C**) SEC profiles for C3, C4, and C5 designs match the expected assembly states of the respective backbones shown on the right. (**D**) The observed molecular weights of designs determined by SEC-MALS are in close agreement to those computed from the design models. Error bars correspond to the standard deviation. (**E**) PI25 structural overlays between the design model (grey) and the solved structure (green) with side-chain rotamers of the solved structure to LHD101 interface seed (grey). (**F**) PI31 structural overlays between the design model (grey) and the solved structure (green) with side-chain rotamers of the solved structure to LHD101 interface seed (grey).(**G**) PI57 structural overlays between the design model (grey) and the solved structure (green) with side-chain rotamers of the solved structure to LHD101 interface seed (grey).

We compared the diversity of backbones generated from a single interface seed orientation or from random seed orientations initialized in a range of radial distances (Fig. 2B). Through structural clustering of 1000 designed backbones from LHD101, we confirmed that sampling the seed orientation increased the obtained diversity by > 13 fold. The radial repositioning during the diffusion trajectory slightly reduces the generated diversity, while avoiding loosely packed backbones by constraining the translational degree of freedom. After sequence design and *in silico* structural validation, we selected 27 C3, 21 C4, and 12 C5 designs from LHD101 and 4 C3 designs from LHD29 for experimental *in vitro* characterization. The selected designs were expressed in *Escherichia coli* with either an N or C-terminal polyhistidine-tag and 63/64 designs were successfully expressed. 33/63 had retention volumes in size exclusion chromatography (SEC) consistent with the size of the designed oligomeric state (Fig. 2C, Fig. S2), and for 18/63 the molecular weight determined by SEC–multi-angle light scattering (SEC-MALS) matched the design model (Fig. 2D, Fig. S3).

To evaluate structural accuracy, we attempted crystallization of 9 designs and determined the structure for four assemblies with resolution <3.3Å. Two C3 oligomers (PI25 and PI31) and one C4 oligomer (PI57) aligned closely with their design models (RMSD_Cα_ < 1.6 Å), including the junction adjacent to the two sides of the LHD101 interface and the PPI itself (heavy-atom sidechain RMSD 1.5Å, 1.7Å, and 1.7Å for PI25, PI31, and PI57 respectively) (Fig. 2E,G, I). The crystallographic structures of PI25 and PI31 contain multiple non-crystallographic symmetry copies of the trimer, identified as such based on the trimeric state observed by SEC-MALS, within a single asymmetric unit. These NCS copies exhibit pairwise RMSD_Cα_ < 1 Å, further substantiating the accuracy of the structural design. While the monomer of the PI56 C4 symmetric design closely matched the design model (RMSD_Cα_ ≈ 1.6 Å), it formed an alternative C5 symmetric assembly (Fig. S4A). As the SEC–MALS determined mass matched the intended C4 geometry, the observed alternative assembly geometry likely reflects a crystallization artifact arising from the inherent flexibility of the backbone. Interestingly, a posteriori AF3-based predictions suggested that the tetrameric state is only weakly favored over the alternative pentameric state (ipTM 0.82 and 0.79, respectively) (Fig. S4B). In contrast, for PI57, whose tetrameric assembly is supported by both SEC-MALS measurements and the determined crystal structure, AF3 predictions revealed a substantial confidence gap between the designed tetrameric and an alternative pentameric assembly (ipTM 0.80 and 0.55, respectively). This observation suggests that negative *in silico* selection against alternative assembly architectures may improve overall on-target success rate.

Despite originating from the same interface seed (LHD101), the solved assemblies exhibited distinct backbone geometries (pairwise TM-scores ≤ 0.71), underscoring the design diversity generated by our interface-seeded design strategy. These designs are also distinct from structures in the PDB based on FoldSeek search (*31, 32*) with TM-scores ≤ 0.56 (Fig. S5). These results demonstrate that validated PPIs can be reliably repurposed to generate cyclic oligomers with novel topologies, varied radii and symmetries, providing a foundation for constructing protein assemblies of higher structural and functional complexity.

### Design of metal and small molecule-responsive protein oligomers

We next applied our interface-seeded framework to design metal or small molecule-dependent assemblies. In contrast to previously developed approaches, our interface-seeded strategy only requires a validated PPI, simplifying the design problem and allowing for generalisation. We thus sought to design trimeric oligomers that respond to: Cu²⁺, repurposing a Cu²⁺-dependent homodimeric PPI engineered into cytochrome (*33*); cholic acid (CHD), repurposing a CHD-dependent heterodimeric *de novo* PPI (*34*); venetoclax (LBM), repurposing a BCL2-*de novo* binder interface responsive to LBM (*35*); and progesterone (P4), repurposing an antibody-*de novo* binder interface responsive to P4 (*35*) (Fig. 3A). These PPIs pose a new challenge compared to the previously tested, non-responsive PPIs (LHD heterodimers) as they are composed of more than two discontinuous segments (4 for Cu^2+^ responsive oligomers, 5 for CHD responsive oligomers, 5 for LBM responsive oligomers, >8 for P4 responsive oligomers) distributed among two distinct spatial protein regions, raising the challenge of finding compatible backbone topologies that can present both binding sites reliably.

**Fig. 3.**
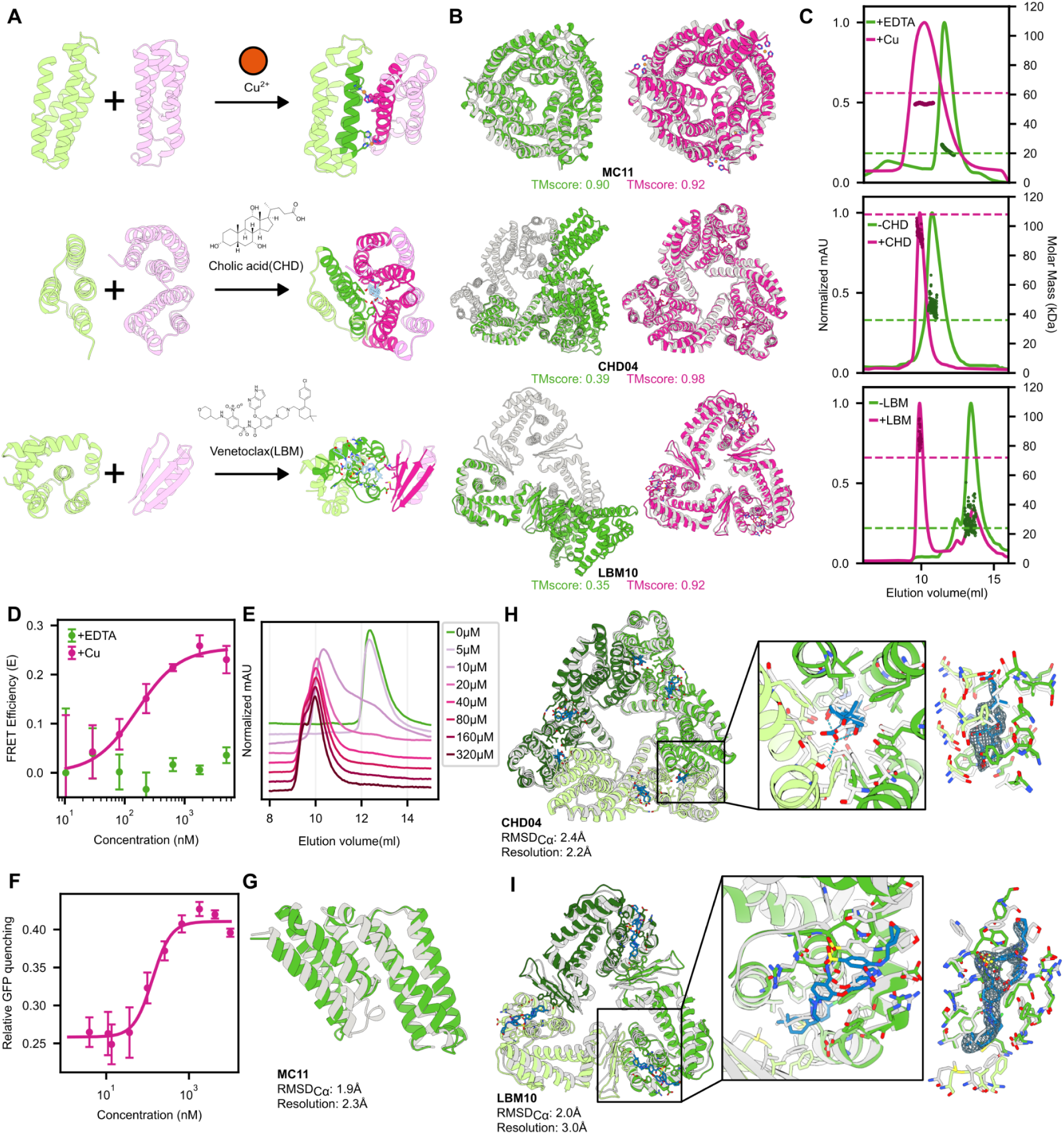
*De novo* design of metal and small molecule-responsive trimers. (**A**) Functional interface seeds are extracted from chemically induced dimers that are responsive to different stimuli. From top to bottom: Cu^2+^, CHD, and LBM. (**B**) Designed backbone (grey) and AF3 predictions of the corresponding design with (pink) and without (green) stimulus, suggesting (improved) assembly fidelity in the presence of the stimulus. (**C**) SEC-MALS profiles for the responsive oligomer designs shown in (**B**). Dashed lines representing the theoretical molecular weight with (pink) and without (green) stimuli. (**D**) Assembly affinity (K_d_) measurement for MC11 using a FRET-based assay. Fluorescent protein fusions at a 1:2 donor:acceptor ratio were used to monitor trimer assembly, while Cu²⁺ was maintained at a 50-fold molar excess relative to protein concentration (**E**) SEC profiles for Cu^2+^ serial titration to 10 μM MC11. Concentration of Cu^2+^ is coloured with a gradient of green-to-pink. (**F**) Assembly affinity measurement for LBM10 using GFP quenching by LBM. Relative GFP quenching of titrated LBM10 in the presence of 10 µM LBM, calculated after subtraction of the matched GFP control and fitted to estimate an effective K_d_. (**G**) MC11 structural overlays between the design model (grey) and the solved protomer structure (green). The solved off-target oligomeric structure can be found in Fig. SX. (**H**) CHD04 and (**I**) LBM10 structural overlays between the design model (grey) and the solved crystal structure (green). The binding pocket containing the small-molecule ligand (blue) is shown in the middle and right panels in two different orientations. Side-chain rotamers and the Polder omit map, contoured at 4.5σ and 4.2σ respectively, demonstrate ligand occupancy within the designed binding interface.

Using this Cu^2+^-responsive interface seed, we generated a series of trimeric assemblies and selected 16 designs for experimental evaluation. Expression and purification followed the same experimental pipeline described above. In the absence of metal and in the presence of EDTA, the SEC retention volumes of 12/16 designs were consistent with monomeric species (Fig. S6). To assess metal responsiveness, we added Cu^2+^ at a 2:1 metal:protein stoichiometry and determined the respective oligomeric state by SEC. We observed metal-dependent assembly for 4/16 designs (Fig. S6), one of which (MC11) was further analyzed by SEC-MALS and formed the intended C3 symmetric trimer (Fig. 3C). Cu^2+^ titration confirmed correct oligomer formation for MC11 only at the expected stoichiometric ratio and above, with the formation of smaller intermediate assemblies at substoichiometric ratios (Fig. 3F). Using fluorescent protein (FP)-fused variants of MC11, we further measured FRET at a range of protein concentrations (10 nM to 5 μM) in the presence of stoichiometric Cu^2+^, and determined the effective affinity of the Cu²⁺-dependent oligomer to be K_d, eff_ = 137 nM (Fig. 3D).

Building on the success of our metal-responsive designs, we generated C3 assemblies for three small molecule-responsive interfaces and selected 16 CHD-, 14 LBM-, and 7 P4-responsive designs for experimental characterization. Following expression and purification as described above, 5/16 CHD-responsive designs had SEC retention volumes consistent with monomeric species in the absence of CHD (Fig. 3C, Fig. S7). After incubation with CHD, 3/5 designs underwent ligand-dependent assembly and formed the intended C3 oligomers. Among the 14 LBM-responsive designs, 9/14 designs were soluble and well expressed. As assessed by SEC analysis, 4/9 designs possessed well defined profiles, where 1/4 possessed only monomeric species, 1/4 only oligomeric species, and 2/4 possessed mixed species. Among the 7 P4-responsive designs, 5 were soluble and displayed moderate expression levels. 4/5 of these showed monodisperse peaks by SEC, of which 1/4 designs possessed only monomeric species, 1/4 only oligomeric species and 2/4 with mixed species. Neither LBM or P4-responsive designs were responsive to small molecule addition (Fig. S8). To improve the designs, we partially diffused the previously selected backbones, which after sequence redesign resulted in higher confidence AF2 monomer predictions (*36*). Based on improved AF3 trimer prediction confidence in the presence of the respective small molecule, we selected 7 LBM-responsive designs for a second round of experimental validation. In the absence of LBM, 6/7 LBM-responsive designs had SEC retention volumes consistent with monomeric species, one of which (LBM10) formed the intended C3 oligomer upon incubation with LBM (Fig. 3C, Fig. S9). We observed sfGFP-LBM10 fluorescence quenching upon addition of venetoclax, indicating assembly (see Methods). We titrated sfGFP-LBM10 protein (5 nM to 10 μM) in the presence of 10 μM venetoclax and determined the effective affinity of the LBM-dependent oligomer to be K_d, eff_ = 130 nM (Fig. 3F), in good agreement with the K_d_ of 96 nM reported for the original hetero-dimer (*35*). Designs in each responsiveness category are structurally novel, with Foldseek TM-scores to their best match of 0.43, 0.23 and 0.37, respectively for MC11, CHD04, and LBM10 (Fig. S10).

We subsequently attempted crystallization of one design of each type of ligand-responsive oligomer and determined the structures for all 3 assemblies. For the Cu^2+^ responsive oligomer, we observe good agreement (RMSD_Cα_ < 2 Å) between the determined monomer structure and the design model with 2.3 Å resolution (Fig. 3G). However, the crystallized protomer excludes Cu^2+^ ions and appears to pack as a dimer of dimers (Fig. S11), despite SEC-MALS indicating that the protein is exclusively monomeric in solution in the absence of Cu^2+^ (Fig. 3C). This result may be explained by the presence of substoichiometric Cu²⁺ and, consequently, unbound MC11 in the crystallization condition. Experiments conducted at a 10:1 Cu²⁺:protomer ratio ratio yielded non-diffracting crystals from a distinct precipitant solution. For the CHD and LBM-responsive designs, the determined crystal structures closely match their design models (RMSD_Cα_ 2.4 and 2.0 Å, respectively) with <3.0 Å resolution, (Fig. 3H, Fig. 3I), including all designed features, e.g. longer loop regions and helical kinks, and the interface seed bound to the small molecule (heavy-atom sidechain RMSD 1.0 and 1.8 Å respectively) (Fig. S12). The crystal induced dihedral packing of the LBM10 design, mediated by an ordered HIS-tag conformation, and appeared to cause a single rotameric deviation in LBM binding when compared to the BCL2-LBM complex (Fig. S12B, Fig. 3I). For CHD04, we additionally resorted to Cryo-EM to obtain a structure in a solution state. The design model fits the obtained electron density well, with density present in the binding pocket of each PPI pointing to the presence of bound CHD (Fig. S13). We observed additional density at a surface “pocket”, also present in the crystal, which in view of the high concentration of CHD in solution suggests low affinity binding sites. Curiously, the secondary CHD binding site could be predicted with AF3 when 9 copies of CHD were provided in the input (Fig. S14).

These results demonstrate that interface seeding can be extended to non-continuous ligand–responsive PPI motifs, enabling modular design of stimulus-controlled protein assemblies. More broadly, this framework provides a generalizable strategy for reusing ligand-responsive PPIs to build diverse, controllable protein architectures.

### Design of reversible phosphorylation-dependent protein assemblies

We then sought to challenge the generalisability of our design strategy by applying it to an interface seed with intrinsic conformational dynamics, in order to design reversible phosphorylation (PO)-responsive cyclic protein assemblies. Phosphorylation is one of the most abundant post-translational modifications in eukaryotes, and it can rapidly and reversibly regulate protein activity, allowing cells to control key processes such as signaling, metabolism, and gene expression (*37*, *38*). The ability to design PO-responsive oligomers could enable synthetic protein assemblies whose formation and function are directly controlled by cellular signaling pathways.

Here, we designed phosphorylation-responsive oligomers using a *de novo* phosphorylation switch and its conformation-specific binders. The switch was designed to populate two conformational states - an “open” binding-competent and a “closed” binding-hindering state (Fig. 4A). Phosphorylation of a serine-residue in the terminal helix shifts the equilibrium toward the open conformation, in which a cryptic binding site is exposed and recognized by a helical motif to form a PO-dependent PPI (*39*). We reasoned that this dynamic equilibrium could be coupled to a reversible oligomerization system by repurposing the PO-dependent PPI using the interface-seeded framework, whereby phosphorylation drives the transition from a monomer to an oligomeric state (Fig. 4A,B).

**Fig. 4.**
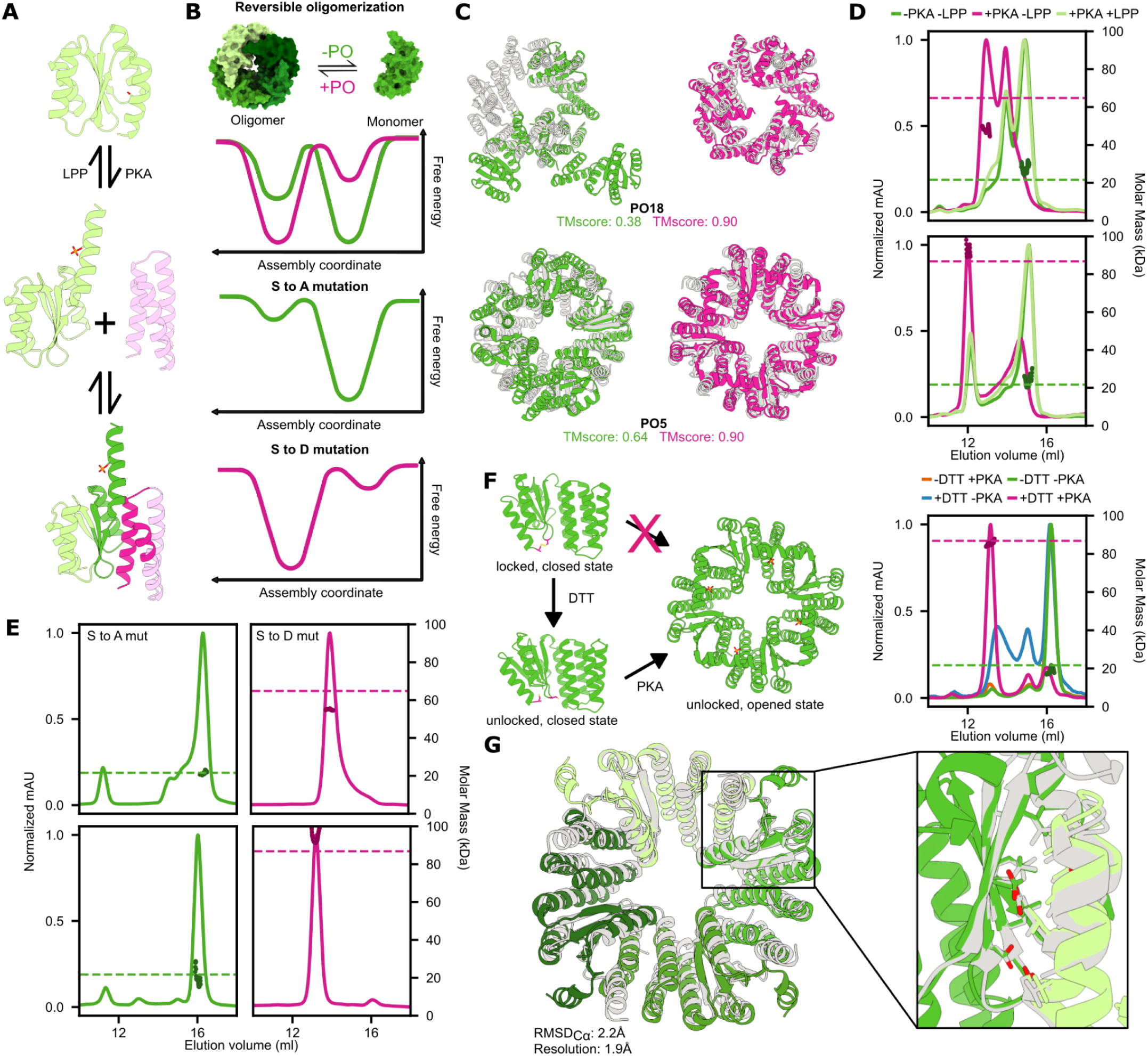
*De novo* design of dynamic phosphorylation-responsive oligomers. (**A**) Phosphorylation-responsive interface seed extracted from a phosphorylation-dependent heterodimer. A dynamic C-terminal helix reversibly undergoes a closed-to-open transition upon phosphorylation. (**B**) The assembly equilibrium is controlled by reversible phosphorylation and can be modulated by single point mutations. Phosphorylation by protein kinase A (PKA) of a serine in the terminal helix shifts the equilibrium from the monomeric to the oligomeric state, while dephosphorylation by λ protein phosphatase (LPP) reverses the assembly equilibrium to the monomeric state. Mutating the phosphorylated serine to an alanine or aspartate strongly biases the equilibrium to the closed, monomeric or open, oligomeric state respectively. (**C**) Designed backbones (grey) and AF3 predictions of the phosphorylation-responsive trimer PO18 (top) and tetramer PO5 (bottom) with (pink) or without (green) phosphorylation. (**D-E**) SEC-MALS profiles confirm the assembly state of both PO18 (top) and PO5 (bottom) in the phosphorylation (D) and mutant backgrounds (E). Data corresponds to the LEU to ALA interface mutant background (PO5s & PO18s), except for the phospho-site mutants of PO5, where the native PO5 was used. Dashed lines indicate theoretical molecular weights in the monomeric (green) and oligomeric (pink) states. (**F**) Engineering a disulfide lock for PO5 renders it responsive to two-inputs simultaneously: phosphorylation and redox. (**G**) Structural overlays of PO5 between the designed model (grey) and solved crystal structure (green). The right panel shows a close-up view of the side chains near the seeded interface.

We selected 19 C3 and 8 C4 PO-responsive designs for experimental characterisation based on the high confidence of their predicted oligomeric states in the phosphorylated relative to the non-phosphorylated state (Fig. 4C). 20 of the designs were soluble and produced well-defined SEC profiles (Fig. S15). 4/20 displayed monomeric species, 4/20 oligomeric species, and 12/20 designs mixed species (Fig. S15). For the designed PO-dependent tetramer PO5 and trimers PO15 and PO18, SEC experiments at low concentrations (0.4-1 µM) revealed a shift in the equilibrium towards the oligomerized state upon phosphorylation by protein kinase A (PKA) (Fig. S16) where phosphorylation was confirmed by an ELISA (Fig. S17). Nonetheless, the assembly PPIs have high affinities and potentially slow dissociation rates from kinetically trapped states in the absence of phosphorylation, likely due to stabilization upon scaffolding coupled to the multivalency of the oligomeric designs. This is supported by the shift of a substantial oligomeric population towards monomeric species upon incubation at 37 °C of the non phosphorylated state (Fig. S18).

To reduce assembly affinity, *in silico* site-saturation mutagenesis (SSM) was used to identify interface mutations predicted to weaken assembly affinity (Fig. S19) (*40*). A single LEU to ALA substitution introduced in the binder interface successfully reduced oligomerization propensity in the non-phosphorylated state while preserving phosphorylation-dependent assembly for PO5 and PO18 (Fig. S19). According to the obtained SEC-MALS profiles, these variants, termed PO5s and PO18s, were predominantly monomeric prior to phosphorylation and almost completely shifted toward tetrameric and trimeric assembly, respectively, following phosphorylation (Fig. 4D, Fig S18). Moreover, oligomerization was reversible, treatment of phosphorylated PO5s and PO18s with λ-phosphatase (LPP) shifted the equilibrium to mostly monomeric SEC-MALS profiles, identical to those of the untreated protein (Fig. 4D).

Mutating the PO-targeted SER to ALA or ASP strongly biased the assembly equilibrium toward monomeric or oligomeric states, respectively (Fig. 4E), demonstrating that single-point mutations can reshape the conformational energy landscape controlling oligomerization.

Additionally, the engineering of a disulfide-bond that locks the “closed” state allows the creation of a system controlled by two separate inputs (Fig. 4F). In the absence of a reducing agent, the disulfide locked PO5 variant (K134C/A183C) is assembly-incapable even after treatment with PKA and ATP, as evidenced by the monomeric SEC-MALS profile (Fig 4F). While treatment with dithiothreitol (DTT) had only a small effect on the equilibrium, recapitulating the profile of non-phosphorylated PO5s, simultaneous treatment with PKA, shifted the equilibrium entirely toward the tetrameric state, consistent with a dual redox/phosphorylation-dependent oligomerization (Fig 4F). To our knowledge, 2 input-control of the oligomerization state of an assembly has not been previously achieved in *de novo* designed systems, opening possibilities for the engineering of higher complexity regulation in synthetic pathways.

Finally, a crystal structure of the PO5 tetramer was determined at a resolution of 1.9 Å, including the terminal helix containing the SER targeted for phosphorylation. Given crystallization screens were set up at protein concentrations well above the original interface K_d_ in either phosphorylated or non phosphorylated state, we resolved the oligomer structure without phosphorylation as we reasoned that the target oligomer could be formed nevertheless.

Consistent with this reasoning, both the core tetramer and the PO-dependent interface seed closely matched the design model (RMSD_Cα_ = 2.2 Å, and heavy-atom sidechain RMSD 1.5 Å, respectively) (Fig 4G). Both PO5 and PO18 designs are structurally novel as assessed by their Foldseek-TM scores to their best-match of 0.32 and 0.35 respectively (Fig. S20).

These are the first examples of biochemically and structurally characterized *de novo* designed oligomers characterized by reversible PO-driven monomer to oligomer transitions. When combined with the potential of controlling assembly by multiple inputs, our results make a key step toward the precise manipulation of computationally designed systems.

### Determinants of design success

We have successfully applied our interface-seeded strategy to design oligomers with different response mechanisms, mediated by diverse chemical triggers. We have found that successful design often requires empiric structural intuition. Unlike conventional motif scaffolding (*41*, *42*), our interface seeds are fragmented and structurally complex motifs, making their topological arrangement a key challenge. For example, to achieve LBM-responsive oligomers, rewiring the scaffold topology to better mimic the original mini-binder architecture substantially improved backbone designability (Fig. S21A). Similarly, in CHD–responsive designs, preserving a native surface helix significantly improved designability over backbone generation conditioned on interface only (Fig. S21B). These examples highlight that even though our pipeline generalizes across diverse functional PPIs, topological sampling conditioned by motif incorporation remains a challenge and thus, successful outcomes benefit from empirical optimization.

Analysis of generated cyclic assemblies at different stages of the design pipeline revealed recurring geometric preferences. While rotational degrees of freedom are randomly sampled at generation, structurally sound backbones occupy distinct preferred regions of the conformational space that are difficult to predict from the interface seed alone (Fig. S21C-H). Despite seed-specific variability, designs chosen for experimental validation consistently favored constrained interface “twist”, with a moderate preference to align quasi-parallel to the symmetry axis rather than perpendicularly, i.e. with a flat arrangement on the plane of the ring (Fig. S21I-K). Analysis of natural C3 ring proteins from the Protein Data Bank(*43*, *44*) revealed similar trends (See Methods), suggesting that PPIs mediating assembly of cyclic oligomers occupy preferred geometric spaces that could guide future cyclic oligomer design campaigns.

### *In vitro* membrane binding controlled by stimuli

*De novo* design has found many applications in bottom-up synthetic biology, providing biochemically well-behaved and characterized building blocks for assembly of systems with complex behaviors often expanding from those observed in cells (*45*, *46*). However, dynamic control over protein localization remains largely unexplored. For example, the function of a diversity of cellular proteins is controlled by their dynamic membrane binding in response to stimuli (*47*). Here, we sought to apply our designs to control membrane-localization by mimicking one such mechanism - triggered membrane-avidity switches (Fig. 5A). Cu^2+^and CHD-responsive designs were genetically fused to (a) the membrane-targeting sequence (MTS) of MinD of *E. coli*, a well-characterized amphipathic helix widely-used as a membrane anchor that only stably interacts with negatively charged membranes in an oligomeric form (*48*), and (b) GFP for visualization using fluorescence microscopy. We incubate each purified construct with negatively charged supported lipid bilayers (DOPC:DOPG 80:20), observing low fluorescence signal at the membrane level in absence of stimulus, in good agreement with the low binding observed for the strictly monomeric construct (*49*) (Fig. 5B-C). Upon addition of the respective stimulus, the signal increases significantly due to oligomerization-driven membrane binding, at levels similar to those of a membrane-targeted obligate trimer (Fig. 5B-C).

**Fig. 5.**
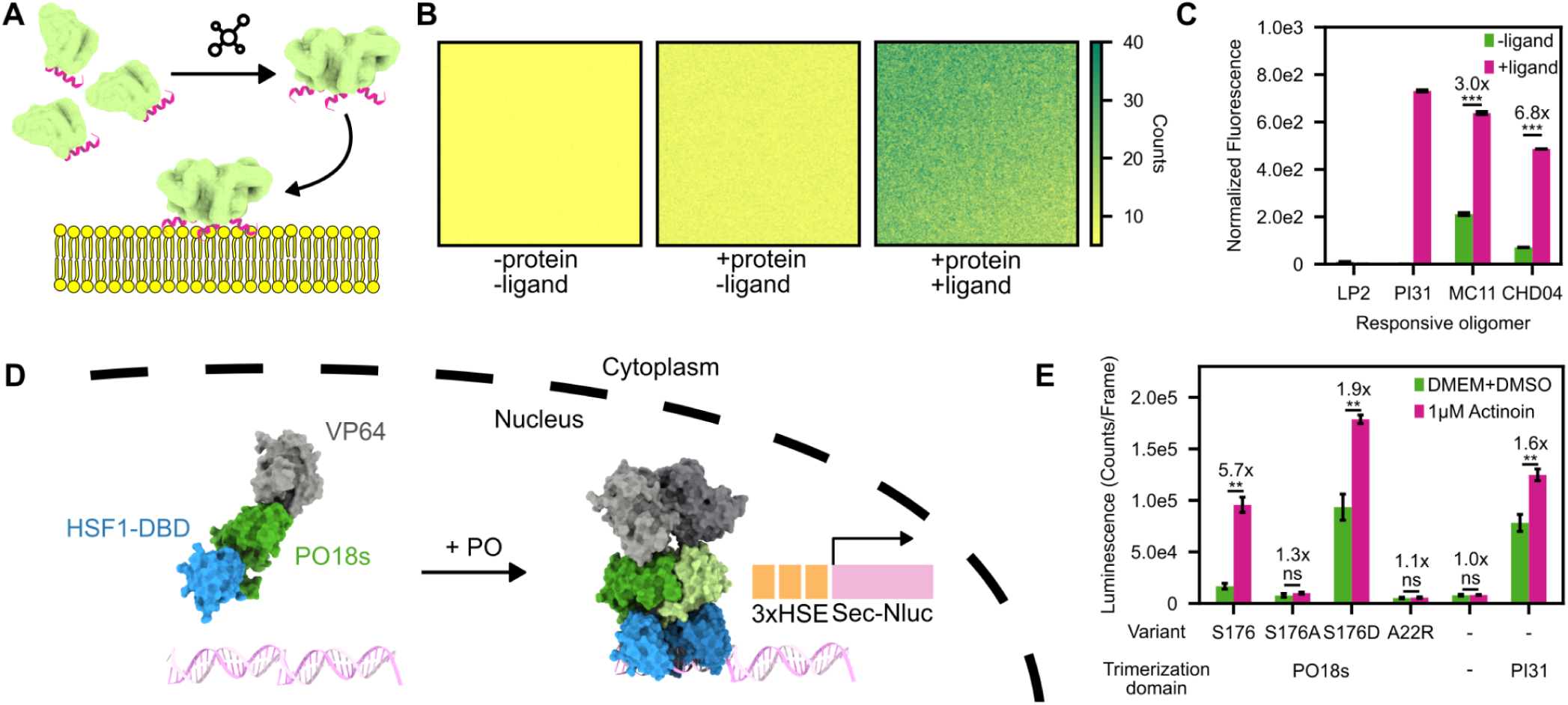
Control of protein localization and gene activation using responsive oligomers. (**A**) Illustration of controllable membrane tethering on Supported Lipid Bilayers (SLBs) with ligand induced oligomers linked with membrane targeting sequence (MTS) (**B**) Cholic acid dependent protein localization on SLB. Representative confocal images of the SLB, after addition of GFP-CHD04-MTS, and after addition of GFP-CHD04-MTS and CHD. (**C**) Quantitation of membrane binding of the designed monomer LP2-MTS, the designed PI31-MTS obligate trimer, MC11-MTS and CHD04-MTS upon addition of respective ligands. Fluorescence counts are normalized based on respective membrane only fluorescence counts. Bar graphs in (**C**) represent mean +/- s.d, performed using 10 frames. Statistical significance was determined using an unpaired two-tailed Welch’s t-test (ns = not significant, * = p < 0.05, ** = p < 0.01, *** = p < 0.001). (**D**) Illustration of phosphorylation-mediated transcriptional regulation. Transcriptional activation of a minimal promoter engineered with HSF1-specific HSE depends on homotrimerization of a synthetic transcription factor comprising a HSF1-derived DNA binding domain (HSF1-DBD), a transactivator domain (VP64), and PO18s. (**E**) Quantitation of secreted Nanoluciferase (Sec-Nluc) levels for the synthetic transcription factor comprising different variants of PO18s, the designed PI31 obligate trimer, or without a trimerisation domain. Bar graphs in **(E)** represent mean +/- s.d, performed using n = 3 biological replicates. Statistical significance was determined using an unpaired two-tailed Welch’s t-test (ns = not significant, * = p < 0.05, ** = p < 0.01, *** = p < 0.001).

These experiments demonstrate the potential of our strategy to contribute to the design of responsive switches that can control protein localization and ultimately aid the engineering of protein-based logic and orthogonal signaling pathways in natural and synthetic cells.

### Phosphorylation-inducible transcriptional control

Many natural signaling systems regulate protein function through the formation of higher-order oligomers (*8*, *10*, *50*, *51*). In many cases, these signaling pathways culminate in the regulation of gene expression, coupling molecular switches to broader cellular decisions. To demonstrate the ability of our responsive oligomers to control transcription, we implemented a cell-based assay based on the trimerization-dependent activation of heat shock factor protein 1 (HSF1). Upon activation, HSF1 forms trimers that bind heat shock response elements (HSEs) and activate transcription (*52*). Inspired by recent work where the native HSF1 trimerization domain was replaced with a chemically inducible trimer (*19*), we tested whether our designed oligomers could similarly induce transcription in response to cellular PKA activity.

To achieve this, the phosphorylation-induced trimer PO18s, was genetically fused to the HSF1 DNA-binding domain (DBD) along with the VP64 transactivation domain (Fig. 5D). We then measured its activity in human embryonic kidney (HEK) 293T cells by monitoring the luminescence of a HSE-specific secreted NanoLuciferase (NanoLuc) reporter (see Materials and Methods). To induce PKA activity, cells were co-transfected with an engineered split-PKA (*53*) fused to a designed chemically-inducible dimer (CID) that assembles upon addition of actinonin (*35*) (Fig. S22A). Treatment with 1 µM actinonin led to an almost 6-fold increase in secreted NanoLuc activity (Fig. 5E). Consistent with their effects on oligomerization, the ALA and ASP mutants at the PO-targeted SER exhibited constitutively low and high reporter activity respectively, with the ASP producing an approximately 12-fold increase in reporter activity compared to the ALA mutant in uninduced conditions. An interface-disrupting ALA-to-ARG mutant showed minimal transcriptional activation as in the absence of a trimerization domain, while our constitutively trimeric design PI31 showed high response. Moreover, with the addition of 25 µM forskolin (*54*) (which increases endogenous PKA activity by increasing intracellular cAMP) and 1 µM actinonin, we observed over 14-fold increases in transcriptional activity for constructs harbouring the PO18s switch (Fig. S22C) as a result of the simultaneous activation of endogenous and engineered split PKA. Additional controls, including the use of the higher affinity PO18 variant, along with a positive control for PKA activity were implemented to further validate the assay (Fig. S22B).

Together, these results demonstrate that *de novo* designed phosphorylation-responsive oligomers can be coupled to endogenous signaling activity to control synthetic transcriptional outputs in human cells.

## Conclusion

*De novo* design of protein assemblies, whether achieved through traditional physical docking or state-of-the-art deep learning methods, has enabled the construction of oligomeric assemblies with diverse architectures for a range of applications. However, most designed assemblies have been static, lacking triggering mechanisms to modulate assembly in response to external stimuli. Our interface-seeded approach expands the designable functional space of protein complexes by reusing responsive interfaces in diverse assembly contexts, going beyond current approaches that generate new conditional interfaces(*17*, *19*). The ability to construct functional oligomers solely from existing interfaces represents an important advance in protein design, providing a generalizable route for creating modular protein materials with dynamic and controllable properties.

Despite the successes outlined here, we have identified several challenges to further improve our interface-seeded approach to the design of responsive protein materials. Our current design protocol relies on the ability of generative models in producing large designable backbones (Fig. S23) (*42*), needing dock-and-design approaches to expand to larger architectures. By reducing the seed to a minimal set of interacting residues, we maximally exploit the generative capabilities of current methods. However, stripping the interface from its original context reduces the predictability over the dynamic properties of the interface, while the formulated design task becomes an advanced multi-motif scaffolding task with a proportionally constrained solution space, further exacerbated by symmetry. While previously developed approaches to reuse validated building-blocks in novel contexts simplify the design task to the generation of a rigid genetic fusion between pre-existing domains (*22*, *55*, *56*), they have only been applied to simpler non-responsive PPIs and are even more limited by the scalability of current design methods.

While these observations highlight the continuous need for design methods scalable to large symmetric assemblies, as well as efficient solutions for sampling the available design space and filter generated designs prior to experimental validation, our interface-seeded framework provides an innovative way for the design of smart protein nanomaterials that sense and respond to the environment.

We foresee that through iterative refinement of our pipeline and expansion of the available collection of characterized interaction modules, we will bring protein design closer to the modularity of nucleic acid nanotechnology, omitting the need for “new purpose, new design”. This “plug-and-play” capability will ultimately enable a wide range of researchers without protein design expertise to rapidly generate smart protein-based nanomaterials tailored to their specific synthetic biology, biotechnology and biomedicine applications.

## Acknowledgements

The synchrotron data was partly collected at beamline operated by EMBL Hamburg at the PETRA III storage ring (DESY, Hamburg, Germany). We also thank Dr. Renato Weiße for support regarding macromolecular crystallography, and Gert Weber, Melanie Oelker and Manfred Weiss for their help at BESSY II, Helmholtz Zentrum Berlin.We thank Emiko Uchikawa and the entire team of DCI Lausanne for access to electron microscopes and support during data acquisition. Computing time was provided by the Leibniz Supercomputing Center on its Linux-Cluster and AI Systems.We thank the Center for Nanoscience (CeNS), the Khmelinskaia’s group members and Anthony Marchand for helpful discussions.

## Funding

Federal Ministry of Education and Research and Bavarian State Ministry for Science and the Arts under the Excellence Strategy of the German Federal Government and the Länder through the ONE MUNICH Project Munich Multiscale Biofabrication (AK, PT) and LMUexcellent (AK)

Federal Ministry of Research, Technology, and Space of Germany and by Sächsische Staatsministerium für Wissenschaft, Kultur und Tourismus in the programme Center of Excellence for AI-research “Center for Scalable Data Analytics and Artificial Intelligence Dresden/Leipzig”, project identification number: ScaDS.AI (CTS)

German Research Foundation through SFB1032 – 201269156 (AK), grant KH 626/2-1–558049915 (AK), INST 86/2224-1 FUGG 519922049 (PT) and under Germany’s Excellence Strategy through the excellence cluster BioSysteM EXC3092/1-533751719 (PT, AK)

Swiss National Science Foundation through the grants 10.005.296 and TMCG-3_213750 (BC)

## Author contributions

Conceptualization: HYC, AK

Methodology: HYC, MI, DR, PS, NMG, BBM, MAJ, BBM, SB, SB, SJ, KL, AK

Investigation: all authors Visualization: HYC, MI, AK

Funding acquisition: LK, CS, BC, AK Project administration: CS, BC, AK Supervision: CS, BC, AK

Writing – original draft: HYC, MI, DR, PS, NMG, BC, AK Writing – review & editing: all authors

## Data, code, and materials availability

All data is available in the main text or as supplementary materials. Scripts, computational methods and design models are available on GitHub (https://github.com/Khmelinskaia-Lab/RFdiffusion_interfaceseed.git and https://github.com/Khmelinskaia-Lab/interface_seeded_oligomers.git), Crystallographic datasets have been deposited in the PDB (accession codes: 31PT, 31PU, 31PV, 31PW, 31PX, 31SW, 31VQ, 31VR, 31VP). EM maps have been deposited in the EMDB (accession codes: EMD-58650).

