## Supplementary Information for "A generalizable interface-seeded framework for *de novo* design of functional oligomers"

**The PDF file includes:**

Materials and Methods
Figs. S1 to S23
Tables S1 to S5
References

### Materials and Methods

Benchmarking the dock-and-design approach

For LHD101-derived dimer docking into different cyclic oligomers, the interface of LHD101 was extracted from the Protein Data Bank (PDB) (*43*, *44*) deposit PDB ID 7MWR by selecting the residues of each chain within 8 Å of the other in pymol. These extracted motifs were then used to generate a library of 100 dimers using the RFdiffusion motif-scaffolding module. Through combinatorial mixing, a library of 10000 dimers was then docked into C3, C4, and C5 symmetric assemblies using RPXDock (*20*) using the SASA-priority scoring function. Docks were filtered based on the obtained rpx score (> 0) and SASA-estimate values (750 $Å^2$ < SASA < 1500 $Å^2$). The same procedure was used on the LBM interface extracted from the design model from Marchand et al. (*35*).

Implementing a new symmetric motif scaffolding module in RFdiffusion

The original symmetric motif scaffolding in RFdiffusion only allows scaffolding from pre-symmetrized motifs. We implemented an RFdiffusion module which can automatically symmetrize the motif and sample their relative positioning during the diffusion process. The motif is randomly oriented within a defined rotational space (where the default is sampling the whole rotational space) at the start of each design trajectory. Then, the motif is symmetrized and initialized within a range of user defined radii. The symmetrized motif is used as input to the RFdiffusion model and after each diffusion step, the center of mass (com) of the diffused noise is calculated to be used as a guide to readjust the radial placement of the motif by applying a pulling force towards the center of the assembly. The updated 3D coordinates ($X_{t-1}$) of the motif are calculated following the equation:

$$X_{t-1} = f(X_t) + \frac{1}{T} \cdot com\_diff_{scaffold} \cdot \frac{M\_int_T}{M\_int_t} \cdot M\% \cdot U(0.5, 1.5)$$

where $f(X_t)$ are the 3D coordinates at time step *t*, T is the total number of diffusion time steps, $com\_diff_{scaffold}$ is the difference in radial distance of the com of the diffused scaffold and the placed motifs, which is the main parameter guiding the pulling of the motif, $M\_int_T/M\_int_t$ is the ratio between the number of interactions between the scaffold and the motif at the start of the diffusion trajectory and that of the current step, which reduces the weight of pulling when the generated scaffold starts forming well-packed regions in contact with the interface motifs, M% is the percentage of protein length that is occupied by the motif, and U(0.5,1.5) adds fluctuations within the updating process by uniformly sampling from 0.5 to 1.5.

Computational design and *in silico* filtering

5000-10000 backbones were generated for each design objective using manually curated contigs. For most design campaigns, the olig_contacts potential has been used as demonstrated in the cyclic oligomer example provided in the RFdiffusion (https://github.com/RosettaCommons/RFdiffusion/blob/main/examples/design_cyclic_oligos.sh ), except for CHD responsive oligomer in which the potentials.olig_inter_all is set to False. Generated backbones are then first filtered using structural filters, i.e. loop percentage and radius of gyration (lowest 50 percentile) to filter out backbones that are likely to be un-designable (unpacked or unstructured), leaving a total of approximately 10% of backbones available for sequence design.

32 amino acid sequences are then designed for each filtered backbone, while the sequence of the interface seed is kept fixed to retain its properties and all chains are tied together to generate

homo-oligomers. We used the SolubleMPNN model with the sampling temperature set to 0.2, using a fixed seed (37), and omitting CYS during the design process. Note that CYS were allowed in the first rounds of design, including the design of obligate oligomers as well as the first set of LMB- and P4-responsive oligomers, although in the latter CYS residues in selected designs were redesigned using SolubleMPNN prior to gene order.

For *in silico* validation, we first predicted from the designed sequences the individual protomers in their monomeric state using model_1_ptm of the AF2 Colabfold (*28*, *29*, *57*) implementation with no MSA information, no templates and 3 recycles. Predicted structures are aligned to the designed backbone using TMalign, designs with monomer plddt < 85, RMSD < 3 Å and TMscore < 0.8 are filtered out. As AF3 and other models alike were not available at the time of $Cu^{2+}$-metal ion responsive oligomer design, designs were selected solely based on monomer prediction metrics. For the remaining design campaigns, the oligomeric states of designs passing these filters are predicted. More specifically, for the obligate oligomer, we used model_1_multimer of the AF2 Colabfold implementation with no MSA information, no templates and 3 recycles. Predicted structures were aligned to the designed oligomeric backbone using MMalign and designs with multimer plddt < 85 and TMscore < 0.8 are filtered out. Oligomers are then scored using Rosetta to confirm that the interface ddG, buried SASA, and interface shape complementarity did not get strongly penalised during design. . For the small molecule-responsive designs and PO-responsive designs, AF3 was used to predict the respective oligomeric state with and without stimulus. Designs with poor prediction confidence without stimulus (plddt < 75) were selected based on the high prediction accuracy for the oligomer with stimulus, i.e, multimer TMscore > 0.8, plddt > 85, and RMSD < 3 Å. A subset of designs were selected for *in vitro* experimental validation after visual inspection.

*In silico* SSM for PO designs
In silico SSM was carried out as described previously (*58*). Briefly, adjacent pairs of protomers were extracted from the AF3 predictions of phosphorylated PO5, PO15 and PO18. All residues C-terminal to the TEV cleavage site were removed from the models prior to analysis. Residues from the conformation-specific binder located within 4 Å of the exposed hydrophobic cryptic binding site were individually mutated to every other canonical amino acid using a RosettaScripts protocol. For each mutation Rosetta 'Ddg' and 'EnergyPerResidue' (EPR) scores were calculated to evaluate effects on binding affinity and local stability. Candidate mutations were picked based on their predicted ability to weaken but not fully abolish the oligomerization interface. All selected mutations were subsequently inspected visually to corroborate the SSM result. For PO5, three alanine substitutions (L18A, L25A, L26A) were selected for experimental validation, while in the case of the trimers, a single substitution was selected for experimental characterisation: L20A for PO15 and L18A for PO18. Note that L18A for PO5, L20A for PO15 and L18A for PO18 correspond to the same position within the interface seed shared amongst all 3 designs.

Protein expression and purification
Cloning, expression and purification were adapted from Qian et al. (*59*) with modifications for larger expression volumes. Designed amino acid sequences were reverse-translated and codon-optimized for expression in *Escherichia coli* using the GenScript online codon optimization tool (https://www.genscript.com/tools/gensmart-codon-optimization). Codon-optimized sequences were flanked with cloning adapters and ordered as synthetic DNA fragments from IDT or GenScript.

Gene fragments were assembled into either LM627 (https://www.addgene.org/232210/) or LM1371 (https://www.addgene.org/232214/) expression vectors using Golden Gate assembly, generating constructs with either a C-terminal or N-terminal hexahistidine (6×His) tag under the control of a T7 promoter. Cloning reactions were prepared in 10 µL volumes containing BsaI-HFv2, T4 DNA ligase, and DNA fragments in T4 ligase buffer. Assembly reactions were incubated at 37°C for 20 min, followed by 60°C for 5 min.

Assembled plasmids were transformed into laboratory-prepared chemically competent DH5α *E. coli* cells. For each transformation, 1 µL of the Golden Gate reaction mixture was added to 10 µL of ice-cold competent cells and incubated on ice for 30 min. Cells were heat-shocked at 42°C for 30 s and returned to ice for 2 min. Subsequently, 100 µL of SOC medium was added, and cells were recovered at 37°C for 1 h with shaking at 700 rpm before plating on LB agar supplemented with kanamycin and incubated overnight. Individual colonies were inoculated into 2 mL LB medium containing kanamycin and grown overnight for plasmid propagation. Cells were harvested by centrifugation, plasmids were purified using commercial miniprep kits (QIAGEN QIAprep Spin Miniprep Kit), and purified plasmids were transformed into laboratory-prepared chemically competent BL21(DE3) *E. coli* cells using the same protocol described above.

For protein expression, individual BL21(DE3) colonies were inoculated into 50 mL auto-induction medium consisting of TB II supplemented with glycerol (5 g/L), glucose (0.5 g/L), lactose (2 g/L), $MgSO_4$ (2 mM), and kanamycin sulfate (50 µg/mL) in 250 mL baffled flasks. Cultures were grown at 37°C for 8 h and subsequently incubated at 18°C for at least 20 h and a maximum of 28 h with shaking at 750 rpm.

Cell cultures were harvested by centrifugation (30 mins at 4000 rpm) in 50 mL tubes, and cell pellets were resuspended in 10 mL lysis buffer containing 50 mM Tris-HCl (pH 8.0), 250 mM NaCl, 20 mM imidazole, 5% (v/v) glycerol, a pinch of bovine pancreatic DNase I (Sigma-Aldrich) and lysozyme (Merck). Cells were lysed on ice through sonication over a period of 2min, using 2 s on-off cycles (Branson Ultrasonics). Lysates were clarified by centrifugation (30 mins at 4000 rpm), and the resulting supernatants were affinity purified on pre-equilibrated Ni-NTA gravity columns (QIAGEN). Protein-bound resin was washed with 5–10 column volumes (CV) of wash buffer (lysis buffer without DNase I and protease inhibitors), and bound proteins were eluted with 3 CV of the same buffer supplemented with 500 mM imidazole.

Genes encoding the 27 PO-responsive designs and their corresponding mutant variants C-terminally fused to a 2x TRP tag (GSWSW) were ordered from Twist Bioscience and cloned using the Golden Gate Assembly protocol described above. For DNA plasmid extraction, 5 µl of the Golden Gate reaction mixture was transformed into 20-25 µl HB101 E.coli. Plasmids were isolated and purified as described above, and all constructs were verified by Sanger sequencing. Sequence-verified plasmids were transformed into E. coli T7 Express competent cells.

For protein expression, overnight starter cultures (5 mL) were used to inoculate a 500 ml auto-induction medium supplemented with 100 µg/ml ampicillin. Cultures were grown to an OD600 of approximately 0.6 and cultured at 18 °C overnight with shaking at 200 rpm..

Cells were harvested by centrifugation at 4000 g for 30 minutes, and pellets resuspended in 35 ml lysis buffer (50 mM Tris-HCl pH 7.5, 500 mM NaCl, 5% glycerol, 1 mg/ml lysozyme, 1 mg/ml phenylmethylsulfonyl fluoride and 1 µg/ml DNase) then lysed by sonication. Lysates were clarified by centrifugation at 20,000 g's for 20 minutes and filtered with a 0.2 µm syringe filter. Filtered lysates were loaded unto 0.5-1 ml Ni-NTA Superflow (Qiagen) charged HisTrap gravity columns, washed with 5-10 column volumes of 50 mM Tris-HCl pH 7.5, 500 mM NaCl and 10 mM imidazole and eluted with Tris-HCl pH 7.5, 500 mM NaCl, 500 mM imidazole. For PKA, the murine catalytic subunit α was expressed and purified as described previously (*39*).

Size exclusion chromatography (SEC)

Filter-sterilized immobilised metal affinity chromatography (IMAC) elution fractions were further purified by size-exclusion chromatography using either a Superdex 75 10/300 GL column or a Superose 6 Increase 10/300 GL column (Cytiva) connected to an ÄKTA Pure chromatography system (Cytiva) coupled to an ALIAS autosampler (Cytiva) and a fraction collector (Cytiva). Up to 1 mL of IMAC elution fraction was injected and analysed in 25 mM Tris-HCl (pH 8.0), 150 mM NaCl, and 5% (v/v) glycerol buffer, with a flow rate of 0.5 mL/min. Instrument control, data acquisition, and chromatogram analysis were performed using the UNICORN software (Cytiva). Fractions corresponding to the desired elution peaks were collected manually for subsequent analyses.

For the PO-responsive designs, eluted IMAC fractions were injected into a Superdex 200 increase 10/300 GL (Cytiva) or Superdex 200 16/600 GL column (Cytiva), and separated in PBS using an ÄKTA Pure chromatography system (Cytiva). Desired species were collected, concentrated using 3 kDa Amicon filters (Sigma Aldrich), used immediately for subsequent analyses or flash frozen with liquid nitrogen and stored at -80°C.

Size exclusion chromatography coupled with multi-angle static light scattering (SEC-MALS)

Purified protein samples were adjusted to a final concentration of 10 µM in a total volume of 500 µL. For ligand-responsive designs, samples were split into two aliquots representing apo and ligand-treated conditions. Monomeric samples in absence of ligand were incubated at 45°C for 1 h, prior to analysis; for metal-responsive designs, EDTA was included during this treatment at 1 mM. Ligand-induced oligomerization was assessed by incubating samples with the appropriate ligand (20 µM $Cu^{2+}$, 1 µM CHD, or 10 µM LBM) for at least 1 h at 37°C. Following incubation, samples were analyzed by SEC-MALS using a miniDAWN detector (Wyatt Technology) coupled to the ÄKTA Pure system described above. Samples were loaded on a Superdex 75 Increase 10/300 GL column equilibrated in SEC buffer supplemented with the corresponding ligand where applicable and operated at a flow rate of 0.5 ml/min. UV280, differential refractive index (dRI) and light scattering signals were recorded and analysed to determine the molecular weight (MW) of protein species in solution using the ASTRA software (version 8.2.2.119, Wyatt Technology).

To screen for phosphorylation-induced oligomerization, proteins were diluted to concentrations ranging from 0.4-1 µM (0.01-0.02 mg/ml)and were split into unphosphorylated and phosphorylated samples. ATP was added to both reactions at 10x molar excess to the PO responsive protein. For phosphorylation reactions, PKA was added at a 1:1000 molar ratio relative to the PO protein. Both reactions were carried out in a buffer consisting of a 10x PKA reaction buffer (500 mM Tris-HCl, 100 mM $MgCl_2$,1 mM EDTA, pH 7.5) diluted in PBS, and supplemented with 1 mM dithiothreitol (DTT). Samples were incubated at 30°C shaking for 1 hour and subsequently kept on ice until analysis. Oligomeric state was assessed using a SEC-MALS instrument (miniDAWN TREOS, Wyatt Technology) by injecting 110 µl of each sample unto a Superdex 200 10/300 GL column (Cytiva), equilibrated in PBS and operated at a flow rate of 0.5 ml/min. Data was recorded and analysed as above (ASTRA software version 6.1, Wyatt Technology). For screening purposes at these concentrations of analytes, elution volume was used to discern monomeric vs oligomeric species.

For analysis of the reversible oligomerization of PO5s and PO18s, proteins were diluted to 23 µM and split into three conditions: untreated (- PKA/– LPP), phosphorylated (+ PKA/- LPP) and phosphorylated followed by dephosphorylation (+ PKA/+ LPP). Samples treated with PKA were phosphorylated as described above. Unphosphorylated proteins were treated identically, but lacked PKA in the reaction mixture. Following incubation at 30°C for 1 hr, residual ATP was removed by repeated buffer exchange cycles into LPP buffer (50 mM HEPES, 100 mM NaCl, pH 7.5 (>5000x dilution) with 4 ml, 3 kDa Amicon filters (Sigma Aldrich), before protein concentrations were redetermined with a NanoDrop. With minimal residual ATP left in the reaction mixture, both PO responsive proteins were diluted to 7-10 µM (0.2 mg/ml) in LPP buffer supplemented with 1 mM DTT, diluted 10x $MnCl_2$ (P0753S, NEB) with or without LPP (P0753S, NEB) at a concentration of 30 U/nmol of phosphorylated protein substrate. Dephosphorylation was allowed to take place at 30°C over 1 hour. Oligomerization state was determined using SEC-MALS as mentioned above, with the SEC-MALS column equilibrated in LPP buffer instead of PBS.

For analysis of the ALA and ASP mutants of the PO-targeted SER of PO5 and PO18s, proteins were diluted to 7-10 µM (0.2 mg/ml) and the MW distribution was analysed by SEC-MALS in PBS as described above.

To lock the dynamic C-terminal helix in a closed conformation, pairs of cysteines were manually introduced into PO5 at the C-terminus (A183C) and target residues located on the opposing β-turn. 3 candidate disulfide forming sequences (Table S4) were selected based on their ability to form disulfides when predicted with AF3 in single-sequence mode or with templates. Cloning, protein expression and purification were performed as described for the PO-responsive designs, except that cell pellets were resuspended in 120 ml lysis buffer prior to sonication. To probe disulfide formation in the locked variants, proteins were concentrated to 46 µM and analysed by SEC-MALS in PBS. At these concentrations, no oligomeric species were observed, suggesting the cysteine pairs lead to disulfide-locked variants.

For dual redox/PO-dependent oligomerization, a sample of one of the disulfide-locked PO5 variant (K134C, A183C) were divided into four treatment conditions: untreated (-DTT/-PKA), phosphorylated (-DTT/+PKA), unphosphorylated and reduced (+DTT/-PKA) and phosphorylated and reduced (+DTT/+PKA). Proteins were diluted to 9.2 µM (0.2 mg/ml) and phosphorylated and unphosphorylated reaction mixtures were prepared as described for the PO-responsive designs above and where needed (reducing conditions) supplemented with 1 mM DTT. The MW distribution for each mixture was analyzed by SEC-MALS in PBS (pH 7.5) with or without 1 mM DTT, as appropriate. Data acquisition and analysis were performed as described above.

<u>Förster resonance energy transfer (FRET)</u>

Assembly of MC11 was monitored using a Förster resonance energy transfer (FRET)-based assay with a FP-genetically fused variant of the protein. Genes encoding the protein of interest were cloned into vectors containing either sfGFP (*60*) or mScarlet-I (*61*), to generate donor- and acceptor-labeled fusion proteins, respectively. The fusion proteins were expressed and purified separately following the protocol described above. For assembly measurements, donor- and acceptor-labeled proteins were mixed in Corning® low-volume 384-well black flat-bottom polystyrene plates at a 1:2 donor-to-acceptor molar ratio. Prior to sample addition, wells were passivated with BSA to reduce nonspecific adsorption to the plate surface, through incubation with a solution of 0.1 mg/ml  for 10min and subsequent buffer wash. Protein samples were prepared over a final concentration range of 10 nM to 5 µM. To induce oligomerization, $Cu^{2+}$ was added at a 50-fold molar excess relative to the total protein concentration.

Fluorescence in three channels was recorded on an iD5 plate reader (Molecular Devices®) equipped with a monochromator: donor excitation/donor emission ($I^{DD}$), donor excitation/acceptor emission ($I^{DA}$), and acceptor excitation/acceptor emission ($I^{AA}$). Raw intensities were corrected for spectral bleed-through (*62*) using calibration factors determined from donor-only and acceptor-only control wells. The α factor accounts for direct excitation of the acceptor upon donor excitation and was determined from acceptor-only wells as $\alpha = I^{DA}{}_{,a} / I^{AA}{}_{,a}$, where $I^{DA}{}_{,a}$ is the acceptor emission measured after donor excitation and $I^{AA}{}_{,a}$ is the acceptor emission measured after acceptor excitation. The β factor accounts for donor bleed-through into the acceptor emission channel and was determined from donor-only wells as $\beta = I^{DA}{}_{,D} / I^{DD}{}_{,D}$, where $I^{DA}{}_{,D}$ is the acceptor-channel signal measured after donor excitation and $I^{DD}{}_{,D}$ is the donor emission measured after donor excitation.

For samples containing both donor- and acceptor-labeled proteins, the corrected sensitized emission (CSE) was calculated as $CSE = I^{DA} - \alpha I^{AA} - \beta I^{DD}$, where $I^{DA}$, $I^{AA}$, and $I^{DD}$ are the raw intensities measured for the mixed donor–acceptor sample. An effective γ factor was determined experimentally to account for differences in donor and acceptor quantum yields and detection efficiencies. FRET efficiency was calculated as $E = CSE / (CSE + \gamma I^{DD})$, where E is the apparent FRET efficiency. Effective assembly affinity $K_{d,\,eff}$ was obtained from the concentration-dependent change in apparent FRET efficiency by fitting a Hill-type saturation model (*63*).

Venetoclax-dependent GFP quenching assay

Upon addition of venetoclax to LBM10-sfGFP, we observed a decrease in sfGFP fluorescence consistent with venetoclax-dependent quenching. A comparable decrease was not observed for LBM10-mScarlet-I or for MC11-sfGFP, a control protein that is not responsive to venetoclax, indicating that the effect depends both on the fluorophore and on its proximity to venetoclax.

To quantify this effect, LBM10-sfGFP and MC11-sfGFP samples were titrated from 5 nM to 10 µM protein in the presence of 10 µM venetoclax and measured using a plate reader. Purified mScarlet-I was added to each well as an internal fluorescence reference. Fluorescence intensities were recorded in the GFP emission channel, $I^{GfP}$, and in the mScarlet emission channel, $I^{mscarlet}$. Since free mScarlet-I fluorescence was not substantially affected by venetoclax under these conditions, $I^{mscarlet}$ was used to normalize the GFP signal for differences in protein concentration, pipetting, and well-to-well fluorescence intensity.

For each protein, baseline wells without venetoclax were used to determine the unquenched GFP-to-mScarlet fluorescence ratio, $I^{GfP}/I^{mscarlet}$. GFP quenching was calculated as $1 - [(I^{GfP}/I^{mscarlet})_{venetoclax} / (I^{GfP}/I^{mscarlet})_{baseline}]$. To isolate the LBM10-specific component of the signal, relative GFP quenching was calculated by subtracting the quenching measured for MC11-sfGFP from the quenching measured for LBM10-sfGFP at matched protein concentrations. The resulting relative GFP-quenching titration was fitted with a Hill-type saturation model to estimate an effective assembly affinity, $K_{d,eff}$ (62).

ELISA

PO5, PO18 and PIPS2 were diluted to 1 mg/ml (47-79 µM), split into two reactions (unphosphorylated, phosphorylated) and were prepared for phosphorylation as described in above sections. Following phosphorylation, 20 µl of the proteins diluted in coating buffer (PBS, pH 7.4) were added to F96 Maxisorp Nunc immunoplate (Thermo Scientific) and incubated overnight at 4 °C. Plates were then washed and blocked with 200 µl blocking buffer (PBS + 5% BSA) for 2 hours, shaking at room temperature (RT). After blocking, the plates were washed then labelled with 20 µl of rabbit anti-Phospho-PKA Substrate (RRXS*/T*) primary antibody (9624S, Cell Signaling, 1:500 dilution in blocking buffer) for 30 minutes at RT, shaking, then washed and labelled with 20 µl of HRP-conjugated goat anti-rabbit IgG (H+L) secondary antibody (SA00001-2-20UL, Proteintech, 1:1500 dilution in blocking buffer) at RT, shaking for 30 minutes. Following a final wash, plates were developed by adding 50 µl of a 1:1 mixture of TMB substrate and peroxide solution (Thermo Scientific) until desired colouration was achieved. Reactions were quenched with 100 µl of 0.5 M HCl. Absorbance at 450 nm was measured with a Tecan Spark plate reader. All washing steps were carried out with 100 µl of PBS + 0.05% Tween 20.

Crystallization, data collection, and structure determination

Crystallization screening was performed for PPI-designs (PI25, PI31, PI56, PI57, PI43, PI61, PI75, PI50, PI52) and the metal-sensing oligomer MC11, ultimately yielding diffracting crystals for PI25, PI31, PI56, PI57, and MC11 under sub-stoichiometric $Cu^{2+}$ concentration, whereas screening MC11 with super-stoichiometric copper concentration resulted in non-diffracting crystals. Samples of MC11 (3.2 mg/ml (175 µM monomeric MC11) supplemented with 150 µM CuCl) and PI56 (7.5 mg/mL) were crystallized at 20°C with the sitting-drop vapor diffusion method using 200 nl protein in 20 mM HEPES (pH 8.0), 150 mM NaCl buffer plus 200 nl reservoir solution. Crystals of MC11 grew within one day in 0.1 M TRIS (pH 8.5), 1.5 M LiSO4. PI56 crystallized in 0.1 M HEPES (pH 8.5), 20% PEG300 within one day. Protein crystals of PI25 (7.0 mg/mL), PI31 (8.0 mg/mL), PI57 (5.0 mg/mL) were obtained by hanging drop vapor diffusion at room temperature using 1 µl protein in 20 mM HEPES (pH 8.0), 150 mM NaCl buffer and 1 µl reservoir solution. Crystal growth of PI25 in 16% PEG3350, 0.2 M NaForm and of PI31 in 8% PEG8K, 0.1 M NaAcet, 0.05 M MgAcet occurred within three days. PI57 crystallized in 0.1 M HEPES (pH 8.5), 20% PEG300 within 6 months. Crystals were cryoprotected using 25 % (v/v) ethylene glycol before flash-freezing in liquid nitrogen.

Crystallographic data collection was conducted at BESSY II Beamline MX14.1 (PI31, PI57) and EMBL/DESY Beamline P14 (PI25, PI56, MC11), respectively(*64*). Diffraction data were processed using the autoPROC or XDSAPP package and phases were determined by molecular replacement using the predicted structures as a search model(*65–68*). Datasets of MC11, PI25, PI31, PI56 were anisotropy corrected using STARANISO (*69*). Manual model refinement was performed in Coot (0.9.8), and automated refinement was carried out with PHENIX.refine (v2.1)(*70*, *71*). In the case of the PI31 dataset, severe pseudo-symmetry was detected, as indicated by a prominent off-origin Patterson peak ($p = 2.20 \times 10^{-4}$) confirming the presence of translational non-crystallographic symmetry (tNCS) within the *a*/*c* plane. To accommodate this structural pathology, tNCS corrections were utilized during both molecular replacement and subsequent structural refinements. Data processing software packages were used from the

SBGrid software environment, which provides curated, version-controlled structural biology applications and reproducible execution environments across platforms(*72*).

CHD04 and LBM10 were diluted to 16 µM (0.6 mg/ml) and 14 µM (0.4 mg/ml) respectively in a buffer containing 25mM Tris-HCl, 150 mM NaCl, pH 8.0, then treated with 200 µM and 50 µM of CHD and LBM respectively. The proteins were then concentrated in 3 kDa Amicon filters (Sigma) to a final concentration of approximately 285 µM and 602 µM for CHD04 and LBM10 respectively (10 and 15 mg/ml) before setting crystal screens. For PO5, following IMAC purification as described above, the protein was further purified by SEC on a Superdex 200 16/600 GL column (Cytiva) equilibrated in a buffer containing 30 mM Tris-HCl, 150 mM KCl, pH 7.5. Fractions corresponding to the oligomeric species of PO5 were collected and concentrated in 3 kDa Amicon filters (Sigma) to a final concentration of 465 µM (10.1 mg/ml) prior to setting up crystal screens.

CHD04, LBM10 and PO5 were crystallized by sitting-drop vapour diffusion at 18 °C using a SPT LabTech Mosquito Robot. CHD04 crystals were obtained in 0.8 M sodium formate, 0.1 M Tris pH 7.5, 25% w/v PEG 2000 MME (*73*). LBM10 crystals were grown in 0.15 M lithium sulfate, 0.05 M magnesium chloride hexahydrate, 0.1 M Bis-Tris, pH 6.8 and 25% (v/v) PEG Smear Low (*73*). PO5 was crystallized in a solution consisting of 0.2 M sodium fluoride and 20% (w/v) PEG 3350 (*73*). Crystals were cryoprotected in 25% (v/v) glycerol and frozen in liquid nitrogen. Diffraction datasets were collected on the MASSIF-1 (ID30A1) beamline of the European Synchrotron Radiation Facility (ESRF, Grenoble, France) at a temperature of 100 K (*74*, *75*). Data were processed with the Global Phasing autoPROC and staraniso packages. Initial phases were obtained by molecular replacement in Phaser using the corresponding design models as search templates. Iterative model building and manual correction were performed in Coot (*71*), followed by refinement using Phenix.refine (*76*). Final model quality was evaluated with MolProbity (*77*).

Data collection and refinement statistics for all samples are provided in Table S1.

<u>Cryogenic electron microscopy (CryoEM) and structure determination</u>

To prepare samples for cryo-EM, CHD04 was subjected to two distinct treatment conditions. In the first condition, CHD04 was diluted to 5.5 µM (0.2 mg/ml) in 5 ml of buffer containing 25mM Tris-HCl, 150 mM NaCl, 5% glycerol, pH 8 supplemented with 500 µM CHD. The sample was incubated at 50 °C for 20 minutes then concentrated at 4 °C to a final volume of approximately 500 µl, before injection into a Superdex 200 increase 10/300 GL column equilibrated in 10 mM HEPES, 100 mM NaCl and 50 µM CHD. Eluted trimeric fractions were collected then concentrated to a final protein concentration of 1-2 mg/ml (27-54 µM) prior to plunge freezing on cryo-EM grids. In the second condition, CHD04 was diluted to 5.5 µM (0.2 mg/ml) in a buffer containing 25mM Tris-HCl, 150 mM NaCl at pH 8 supplemented with 200 µM CHD and then concentrated at 4° C to 2-6 mg/ml (55-164 µM) prior to plunge freezing on cryo-EM grids. All protein concentration steps were performed using 3 kDa Amicon filters (Sigma).

For grid preparation, ultrAuFoil 1.2/1.3 300 mesh grids (Quantifoil) were glow discharged for 90s at 15 mA. Grids were loaded on a Vitrobot Mark IV (FEI), 3 µl sample (1.2 - 5.9 mg/ml) was applied to the grid, blotted (blot force 5, 4s), and plunge frozen in liquid ethane. Grid screening and data collection was done on a 200 kV Glacios Cryo-TEM with a Falcon 4i (Thermo Fisher

Scientific). In total, 2 grids from both sample preparation conditions (1.9 and 4.6 mg/ml) were collected at a magnification of 190000x, physical pixel size of 0.72Å, and a total electron dose of 50 $e^-/Å^2$, resulting in a total of 16270 movies. Collected data was processed in cryoSPARC(*78*). Movies were motion corrected and CTF values were estimated using patch motion correction and patch CTF estimation, respectively. Micrographs were filtered for low relative ice thickness and a CTF fit resolution of 7Å or better. Particles were picked using blob-picker, extracted with a box size of 270 pixels and downsampled to 180 pixels. Junk particles were discarded in the first two rounds of 2D classification. 2D classes were then separated in top and bottom views, and other viewing directions. For top and bottom views, 2D classes were refined in 2 additional rounds of classification, while side views required 4 rounds of 2D classification to obtain high quality classes. Classes were merged, classified another round in 2D and re-extracted unbinned from micrographs with a box size of 324 pixels. A single volume was reconstructed using ab-initio 3D reconstruction using C1 symmetry and refined using non-uniform refinement with enforced C3 symmetry on a subset of particles. Final non-uniform refinement (C3) with all particles resulted in a 3.63Å (FSC at 0.143) reconstruction. Local resolution was estimated and the map was sharpened with a B-value of -50. The initial model was generated using Namdinator(*79*) and cholic acid was docked into the corresponding density in Coot (0.9.8) (*71*). The model was then iteratively refined in Phenix (1.21) (*76*) using automated real space refinement(*80*) and manual refinement in Coot (0.9.8). Overall model quality and fit to map was assessed in Phenix(*77*, *81*).

<u>Diversity and novelty analysis using FoldSeek</u>
Benchmarking was performed using obligate protein oligomers, and structural diversity was quantified across different design parameters. Designed backbones were clustered using the Foldseek easy-cluster algorithm, and diversity was defined as the fraction of unique clusters, calculated as the number of clusters divided by the total number of designed backbones.

Structural novelty of successful designs was assessed using the Foldseek web server by querying each designed backbone against the PDB and the CATH database. Structural matches were first identified in the CATH database. For designs with no detectable matches in CATH (*82*), matches from the PDB50 database were used for visualization and downstream analysis. Structural similarity was further quantified by calculating TM-scores using MM-align (*83*), aligning each designed backbone to its corresponding closest structural match.

<u>Comparison of interface orientations within designed ring-like oligomer and those in the PDB</u>
Rotation angles sampled during the diffusion process were recorded for all designed backbones and projected into a three-dimensional rotation space represented on a unit sphere, where the initial orientation corresponds to the north pole ([0, 0, 1] in Euclidean space).

Ring-like protein trimers, akin to those designed by us, were collected from the PDB using the RCSB API under the following criteria: (i) global symmetry annotation corresponding to C3 symmetry, (ii) a single polymer entity per biological assembly, (iii) three polymer instances per biological assembly, and (iv) polymer entity type classified as protein. Retrieved sequences were clustered using MMseqs2 easy-cluster (*84*, *85*) to remove redundancy and homologous sequences. Representative sequences from each cluster were used to retrieve the corresponding structures in CIF format, which were subsequently clustered again using Foldseek easy-cluster (*31*, *32*) to remove structural redundancy. Representative structures from each cluster were retained.

Protein-protein interfaces were identified in PyMOL using an 8 Å distance cutoff between residues from different chains. Structures in which all three chains participate in a fully connected cyclic interaction network were retained as canonical ring-like protein trimers. All structures were aligned to a common coordinate frame such that the C3 symmetry axis was oriented along the z-axis, and chains were consistently oriented to ensure uniformity across the dataset.

Two geometric descriptors, termed Flatness and Twist, were computed for both the PDB-derived protein ring trimers and all designed backbones described in this study using a custom Python (Python Software Foundation, https://www.python.org/) script. For Flatness, interface residues were defined as those within 6 Å between chains A and B. A representative interface vector was defined using the pair of interface residues (one from each chain) exhibiting the maximum inter-residue distance, identified via a convex hull-based search. Flatness was defined as the angle between this vector and the xy-plane. For Twist, a vector was defined between the com of chains A and B. Twist was defined as the angle between this vector and the xz-plane.

*In vitro* membrane binding

SLB preparations were adapted from Ramm et al. (*86*) with minor modifications. Coverslips were cleaned by mechanical etching between rinses with ethanol (EtOH). After air drying, each coverslip was mounted with a bottom-trimmed inverted 0.5ml tube  to form a sample chamber. Mounted chambers were further cleaned using a plasma cleaner (Diener electronic GmbH & Co. KG) for 10 min at 30% power and 0.3 mbar, using oxygen as the process gas. All lipids were purchased from Avanti Polar Lipids (Alabaster, AL, USA). Small unilamellar vesicles (SUVs) were prepared at a lipid concentration of 4 mg/mL in SLB buffer (25 mM Tris-HCl pH 7.5, 150 mM KCl) using and 80:20 molar mixture of 1,2-dioleoyl-sn-glycero-3-phosphocholine (DOPC) : 1,2-di-(9Z-octadecenoyl)-sn-glycero-3-phospho-(1'-rac-glycerol) (sodium salt) (DOPG) containing 0.05 mol% DiI (Invitrogen). Lipids dissolved in chloroform were dried under a nitrogen stream and further desiccated for at least 30 min to remove residual solvent. Lipid films were subsequently rehydrated in the SLB buffer. SUVs were generated by bath sonication until the solution became clear. For SLB formation, SUVs were added to the reaction chamber at a final concentration of 0.5 mg/mL in SLB buffer supplemented with 5 mM $MgCl_2$. After incubation for 10 mins at 37°C on a heating block, the chamber was washed 10 times with a total of 2 mL SLB buffer to remove excess vesicles.

Confocal imaging of protein recruitment to SLBs was performed on a Luminosa confocal microscope (PicoQuant GmbH). Samples were excited using a 485 nm pulsed laser operated at 0.5 µW with 20 MHz repetition rate and focused through an oil-immersion objective (100×, NA 1.45, UPLXAPO, Olympus). Emitted fluorescence was collected through the same objective, spatially filtered using a 50 µm pinhole, passed through a 582/64 nm bandpass emission filter, detected by a single-photon avalanche diode (SPAD), and recorded using a MultiHarp time-correlated single-photon counting (TCSPC) module (PicoQuant GmbH). Images were acquired over a 20 × 20 $\mu m^2$ field of view with a pixel resolution of 100 nm and a pixel dwell time of 20 µs. For each imaging condition, 10 consecutive frames were recorded and summed to improve signal-to-noise ratio.

The supported lipid bilayer was first imaged prior to protein addition. To verify that lipids formed continuous fluid bilayers, fluorescence recovery after photobleaching (FRAP) experiments were performed by bleaching a centrally positioned 10 × 10 μm² region within a 25 × 25 μm² field of view using a 485 nm laser for 20 s, followed by monitoring fluorescence recovery with 532 nm laser by time-lapse confocal imaging. Subsequently, the protein solution was added and incubated for 2 min before imaging. Following addition of the ligand, the sample was incubated for 5 min and measured. Representative images acquired during the different stages of the experiment are shown in Fig. 5B. Fluorescence intensities were quantified using a custom-built python analysis routine from the summed 10-frame images to assess protein recruitment to the membrane via ligand-induced oligomerization.

HEK293T culture, transfection and induction

Cell culture, transfection and induction were performed as previously described (*39*). Briefly, Human Embryonic Kidney 293T (HEK293T cells) (Invitrogen, R70007) were maintained in Dulbecco's Modified Eagle Medium (DMEM; Gibco, 41966-029) supplemented with 10% (v/v) fetal bovine serum (FBS; Gibco, A5256701) and 1% (v/v) penicillin–streptomycin (Gibco, 15140-122). Cells were cultured at 37 °C under 5% $CO_2$ and routinely passaged every 2–3 days at around 80% confluency. Cell identity was authenticated by the supplier using short tandem repeat (STR) profiling, and cultures were confirmed to be free of mycoplasma contamination by qPCR. For reporter assays, cells were seeded into the inner 60 wells of clear-bottom 96-well plates (Greiner Bio-One, 655-180) at a density of 18,000 cells per well and allowed to adhere overnight. Approximately 24 hours after seeding, plasmids were transfected using polyethylenimine (PEI; Polysciences, 24765-1). Transfection mixtures were prepared in DMEM using a total of 850 ng DNA and 4.125 µg PEI for every six-well column, with volumes scaled to include a 10% excess. A total of 50 µl of transfection mixture was layered on top of the media in each well. Following transfection, cells were incubated for at least 12 h before the culture medium was replaced with fresh medium containing the indicated inducing compounds.

HSF1 secreted NanoLuciferase assay

For each 6-well column of cells in a 96-well plate, transfection mixtures consisted of 50 ng of HSF1-DBD and VP64 fusion constructs along with 280 ng of the HSF1 specific 3xHSE secreted NanoLuc reporter plasmid (pMI282) and 150 ng each of two plasmids each encoding a split PKA fused to one member of a designed actinonin CID (pSB490 and pSB491) (*35*). To maintain a constant total DNA amount of 850 ng, 220 ng of filler plasmid (pCTcon2) was added to the mixture. As a positive control for PKA activity, 250 ng of a secreted NanoLuc CREB reporter (expression regulated by a CRE promoter) was co-transfected with 150 ng of the split PKA plasmids each, along with 300 ng of filler plasmid. At least 12 hours following transfection cells were induced with 1 µM actinonin (AdipoGen, CAS No: 13434-13-4) resuspended in DMSO, while control cells were treated with an equivalent concentration of DMSO in DMEM. 24 hours post-induction, 10 µl of culture supernatant was transferred into a black 384-well plate (Corning) and mixed with 10 µl of diluted Nano-Glo substrate solution (Promega) prepared according to the manufacturer's instructions. After 15 seconds of orbital shaking, luminescence was measured using a Tecan Spark plate reader with an integration time of 1000 ms and an attenuation filter of OD1. Fold changes between uninduced and induced conditions were calculated and statistical significance was determined using an unpaired two-tailed Welch's t-test.

Data plotting and model visualization

Figures created using Inkscape (Inkscape Project. Inkscape Team.). Data processing and plotting were performed with Python (Python Software Foundation, https://www.python.org/). Protein structure rendering was performed in PyMOL (http://www.pymol.org/pymol) or ChimeraX (*87*).

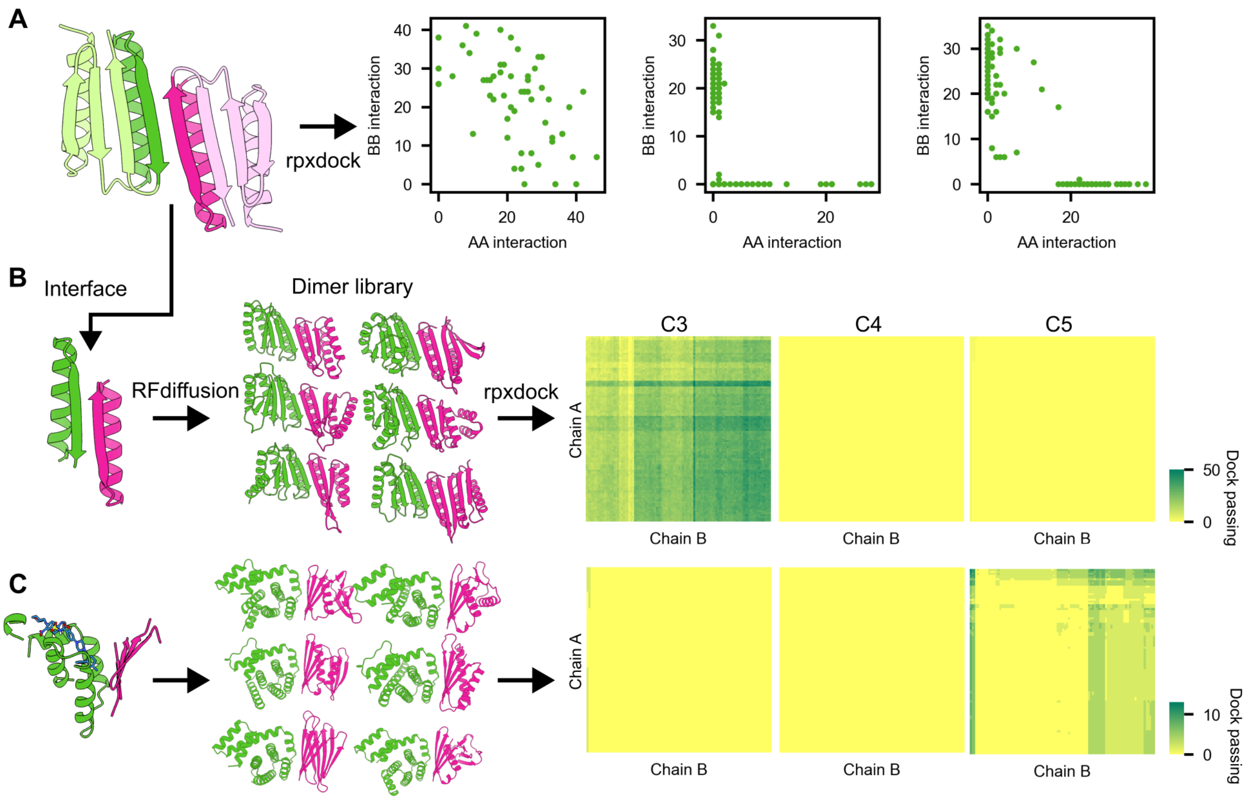

**Fig. S1. Benchmarking the design of cyclic assemblies from hetero-dimeric building blocks using a docking approach.** (A) The LHD101 dimer was docked into cyclic assemblies with different symmetries using RPXDock. The number of homomeric AA and BB contacts are shown as scatter plots to illustrate 2 component-dependent and -independent assembly formation. (B) Interfaces extracted from the LHD101 (C) and LBM-responsive dimers. A dimer library was generated using RFdiffusion. The dimer library was then docked into two-component cyclic assemblies using RPXDock and analyzed based on RPX score and solvent-accessible surface area (SASA).

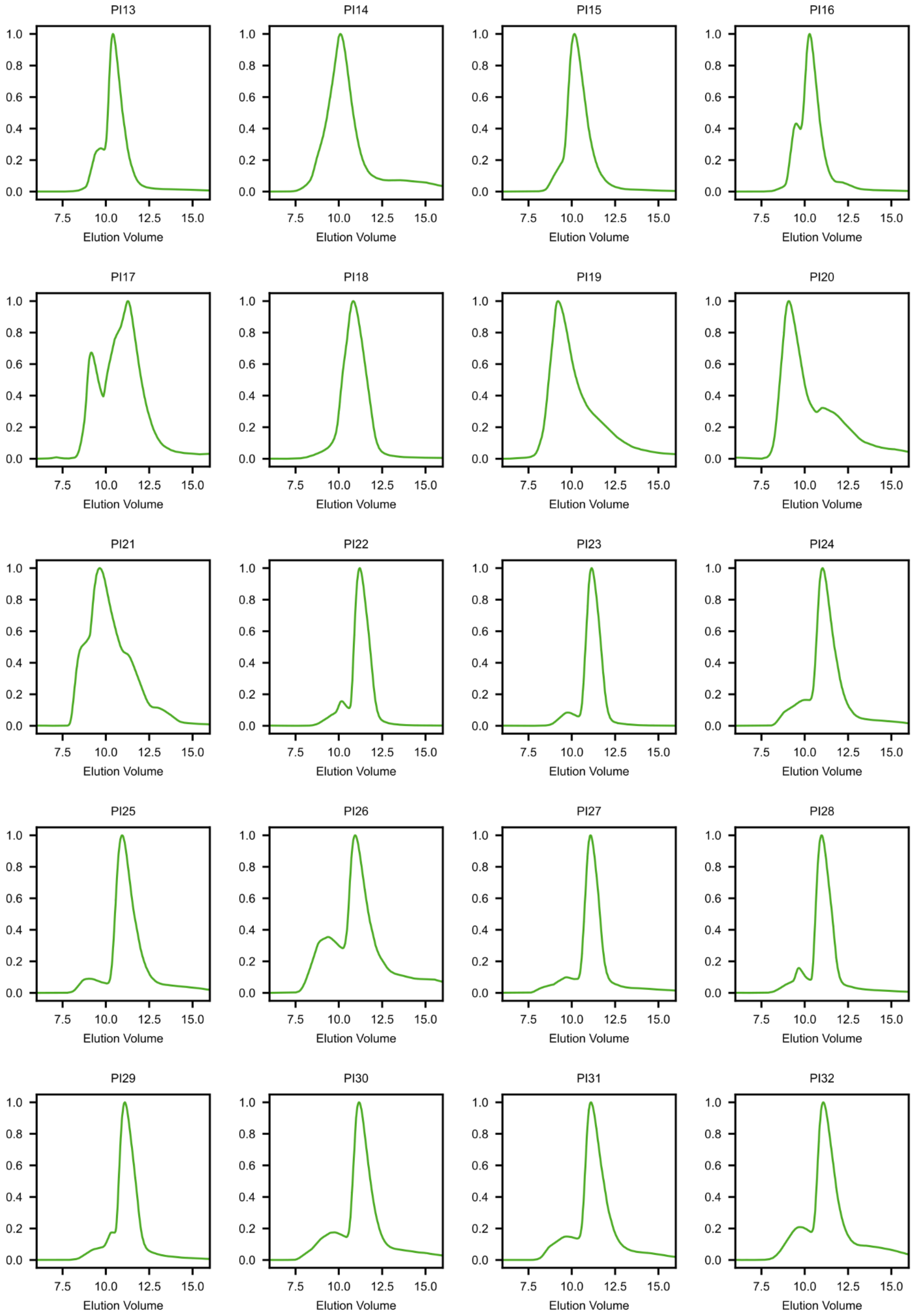
PI13
PI14
PI15
PI16
PI17
PI18
PI19
PI20
PI21
PI22
PI23
PI24
PI25
PI26
PI27
PI28
PI29
PI30
PI31
PI32
1.0
0.8
0.6
0.4
0.2
0.0
7.5
10.0
12.5
15.0
Elution Volume

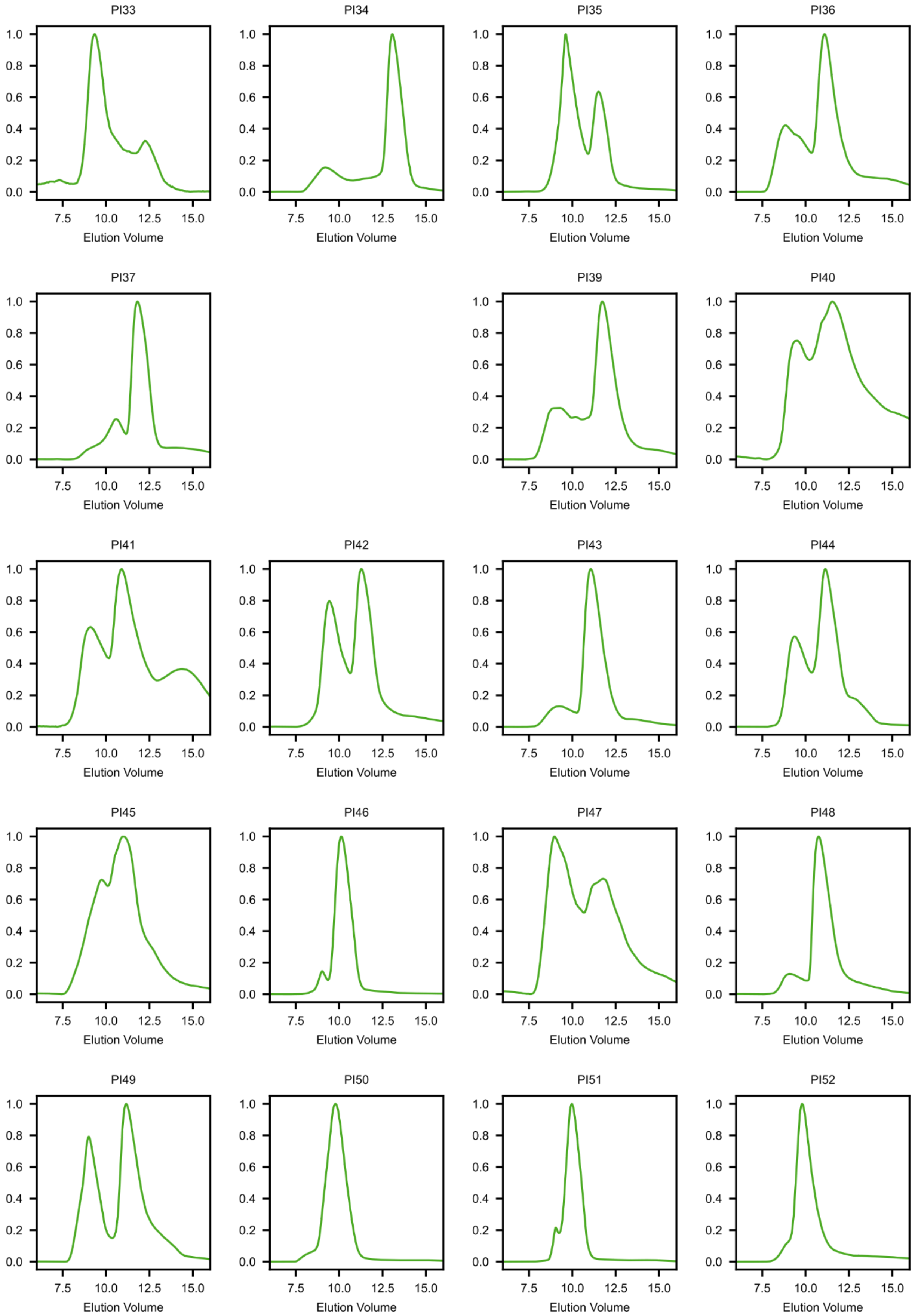
PI33
PI34
PI35
PI36
PI37
PI39
PI40
PI41
PI42
PI43
PI44
PI45
PI46
PI47
PI48
PI49
PI50
PI51
PI52
0.0
0.2
0.4
0.6
0.8
1.0
7.5
10.0
12.5
15.0
Elution Volume

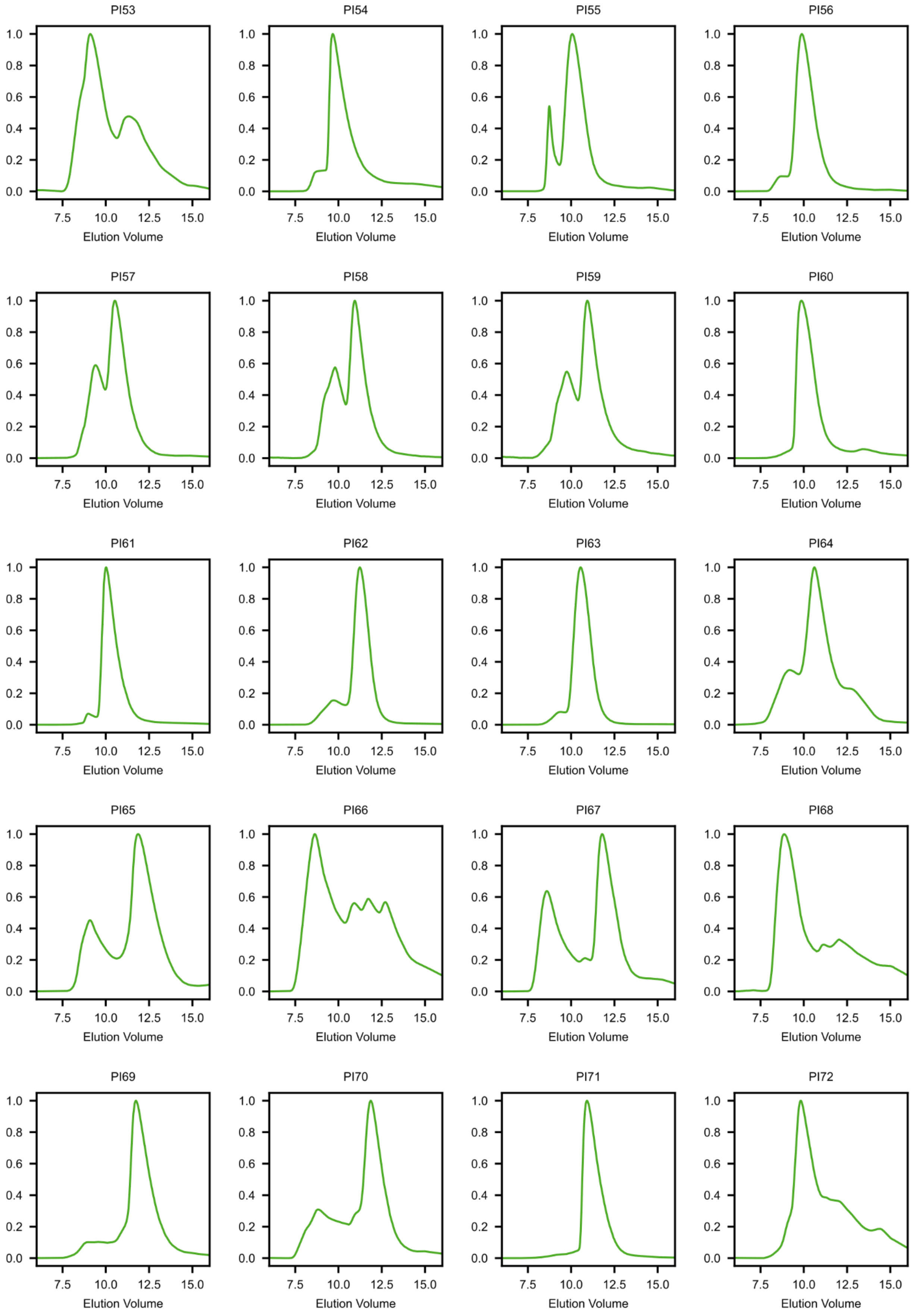
PI53
PI54
PI55
PI56
PI57
PI58
PI59
PI60
PI61
PI62
PI63
PI64
PI65
PI66
PI67
PI68
PI69
PI70
PI71
PI72
0.0 0.2 0.4 0.6 0.8 1.0
7.5 10.0 12.5 15.0
Elution Volume

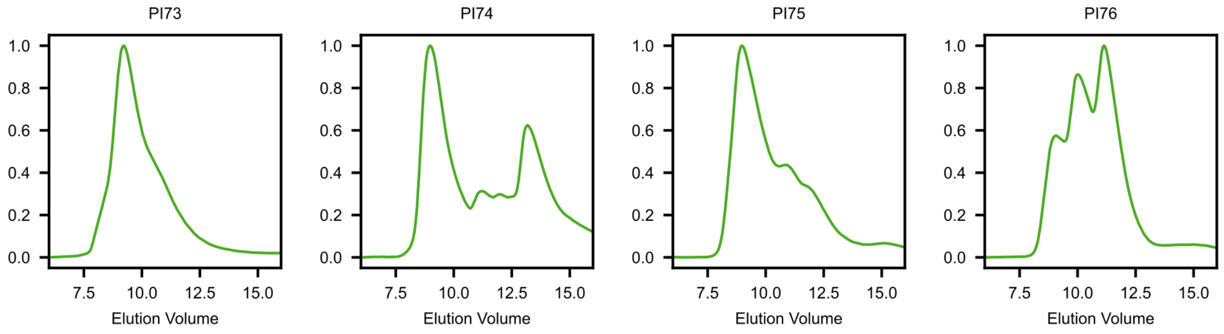

**Fig. S2. Characterization of obligate oligomers based on LHD-derived interface seeds by SEC.** Chromatogram obtained for each designed obligate oligomers using a Superdex 75 10/300 GL column. PI13–PI17 were designed using the LHD29 PPI as the starting seed, whereas all remaining oligomers were designed using the LHD101 PPI. PI13–PI43, PI44–PI64, and PI65–PI76 correspond to designed trimers, tetramers, and pentamers, respectively.

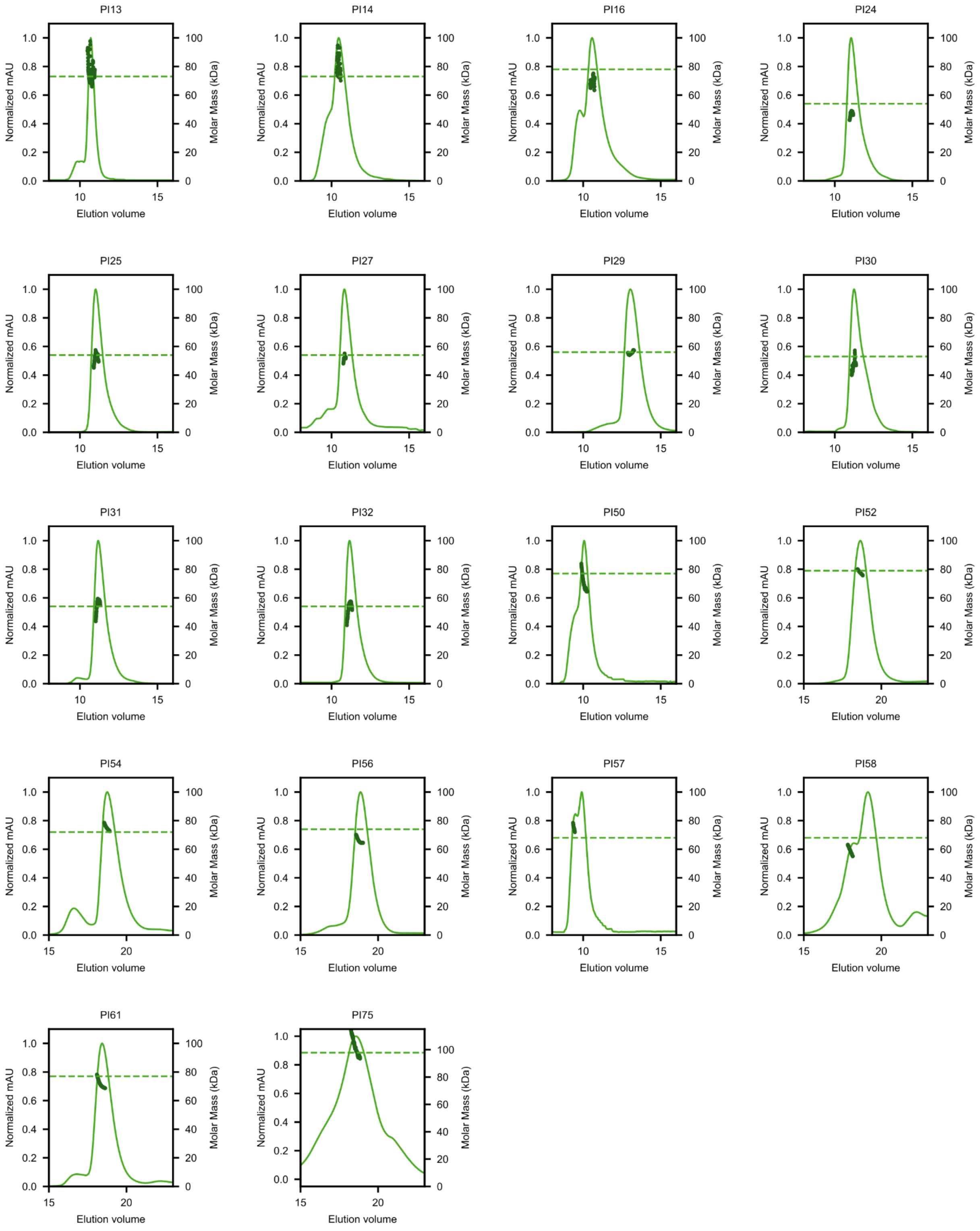

**Fig. S3. SEC-MALS analysis of selected obligate oligomers based on LHD-derived interface seeds.** SEC-MALS traces for designed obligate oligomers whose SEC chromatograms suggest the correct oligomeric state (Fig. S2). SEC elution profiles are displayed in green, experimentally measured MW shown in dark green, and theoretical MW shown in green dashed line.

SEC-MALS measurements for PI52, PI54, PI56, PI58, PI61, and PI75 were performed using a Superose 6 Increase 10/300 GL column, whereas all other designs were analyzed using a Superdex 75 10/300 GL column.

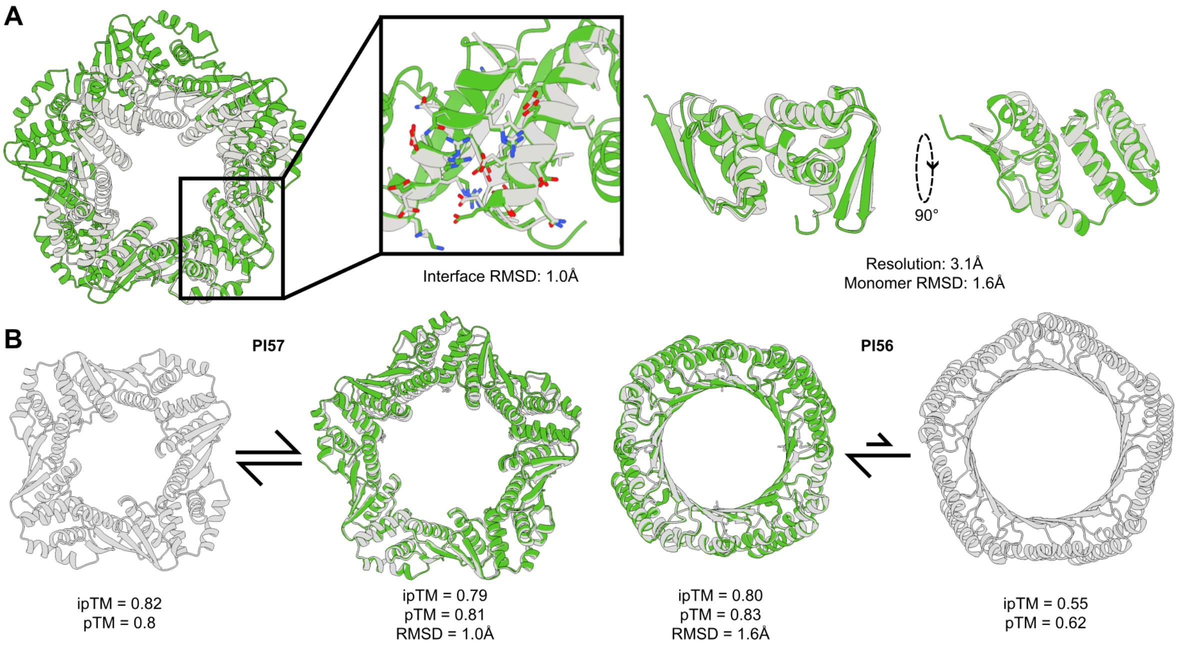

**Fig. S4. Off-target pentameric assembly of PI56.** (**A**) PI56 was experimentally determined to be a pentamer by X-ray crystallography, although only very minor deviations to the monomers extracted from the tetrameric design model are observed. (**B**) Comparison of the AF3 confidence metrics for the on-target tetrameric and off-target pentameric assembly states of PI57 and PI56. Arrows indicate the interpreted potential interconversion between tetrameric and pentameric assembly states based on AF3 confidence scores (ipTM and pTM). AF3 predictions are shown in grey, overlaid with the available crystal structures (green).

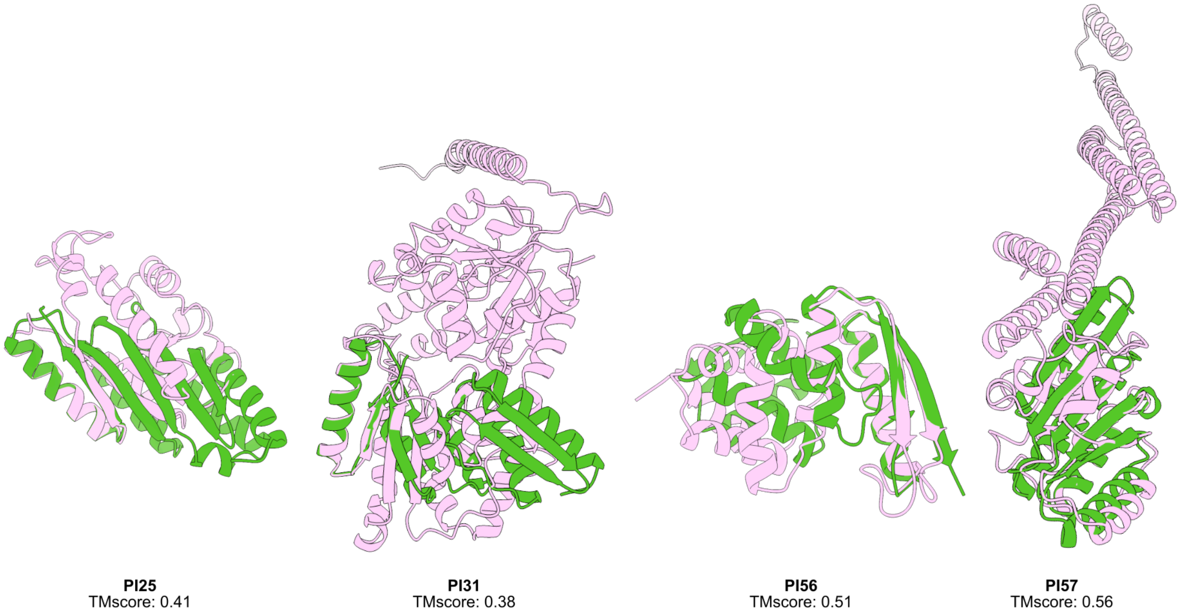

**Fig. S5. Structural novelty of designed obligate oligomers.** Structural novelty was assessed by identifying the closest structural matches (light pink) to the monomeric subunit of each structurally characterized obligate oligomer (green) using the Foldseek webserver searches against the CATH database.

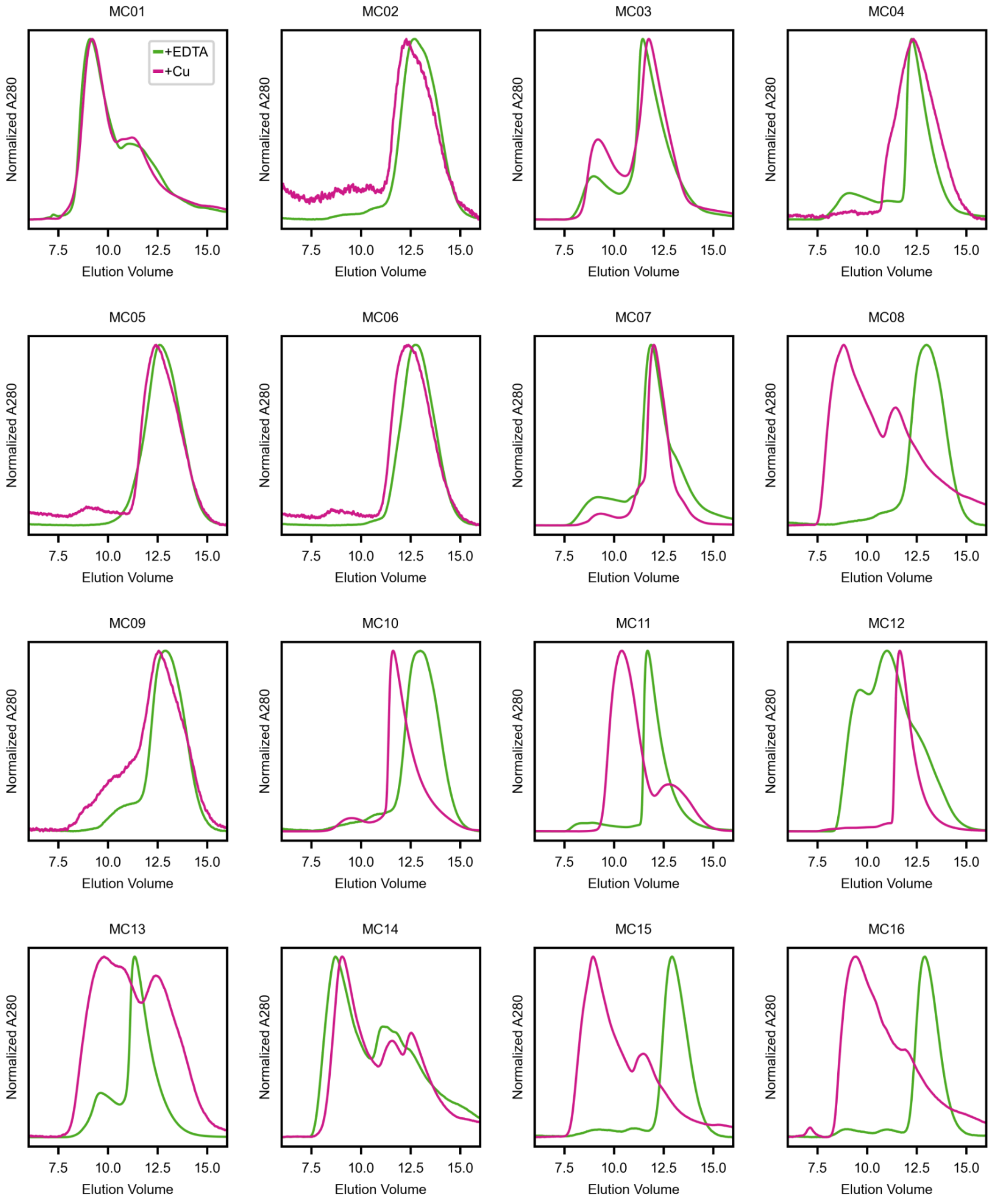

**Fig. S6. Screening of $Cu^{2+}$-responsive designs.** SEC traces of $Cu^{2+}$-responsive designs treated to stabilize the apo (1mM EDTA) or the ligand-bound (100 µM $Cu^{2+}$) state. All measurements were performed using a Superdex 75 10/300 GL column.

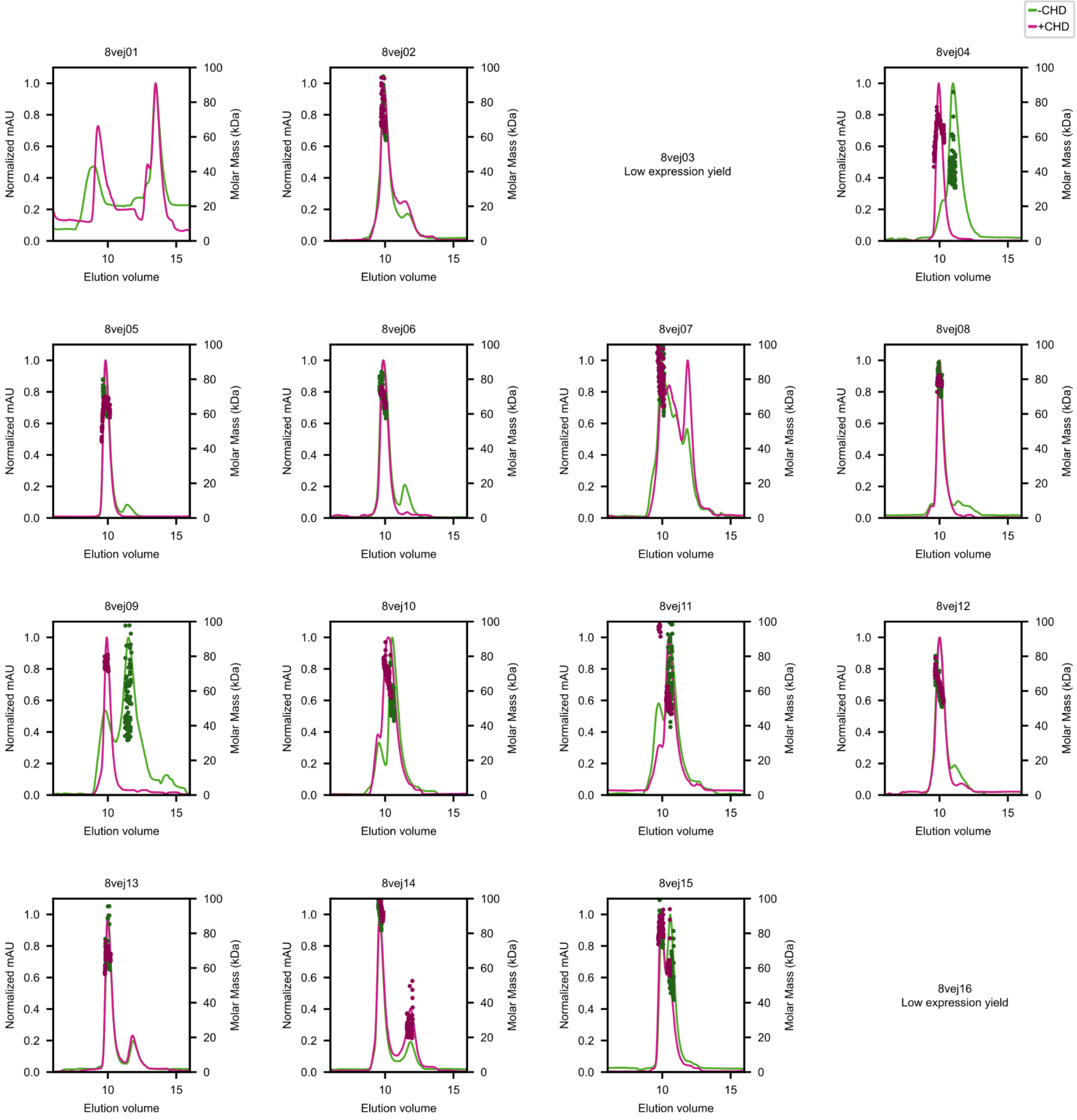

**Fig. S7. Screening of CHD-responsive designs.** SEC-MALS traces of CHD-responsive designs under apo and CHD-treated (10 μM) conditions. SEC elution profiles before and after treatment are displayed respectively in green and pink, experimentally measured MW shown in darker shade, and theoretical MW shown in dotted lines. All measurements were performed using a Superdex 75 10/300 GL column.

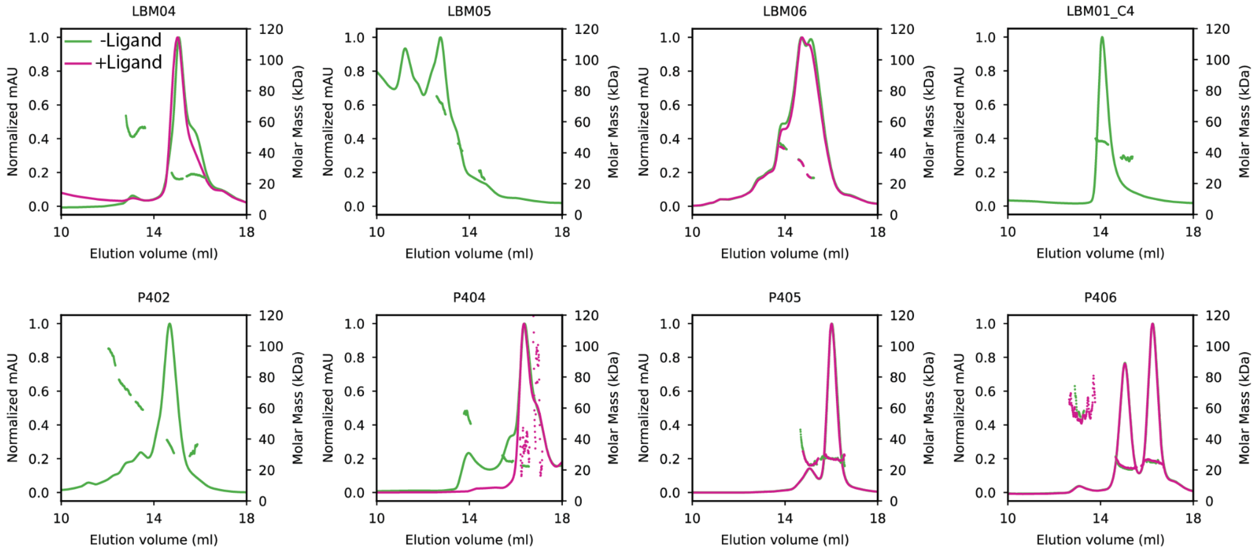

**Fig. S8. SEC-MALS traces of the first round of LBM- and P4-responsive designs.** Proteins at concentrations ranging between 10-50 µM (0.2-1 mg/ml) were injected onto a Superdex 200 10/300 column equilibrated in PBS in the absence (green) or presence (pink) of ligand added at stoichiometric or sub-stoichiometric amounts relative to the protein.

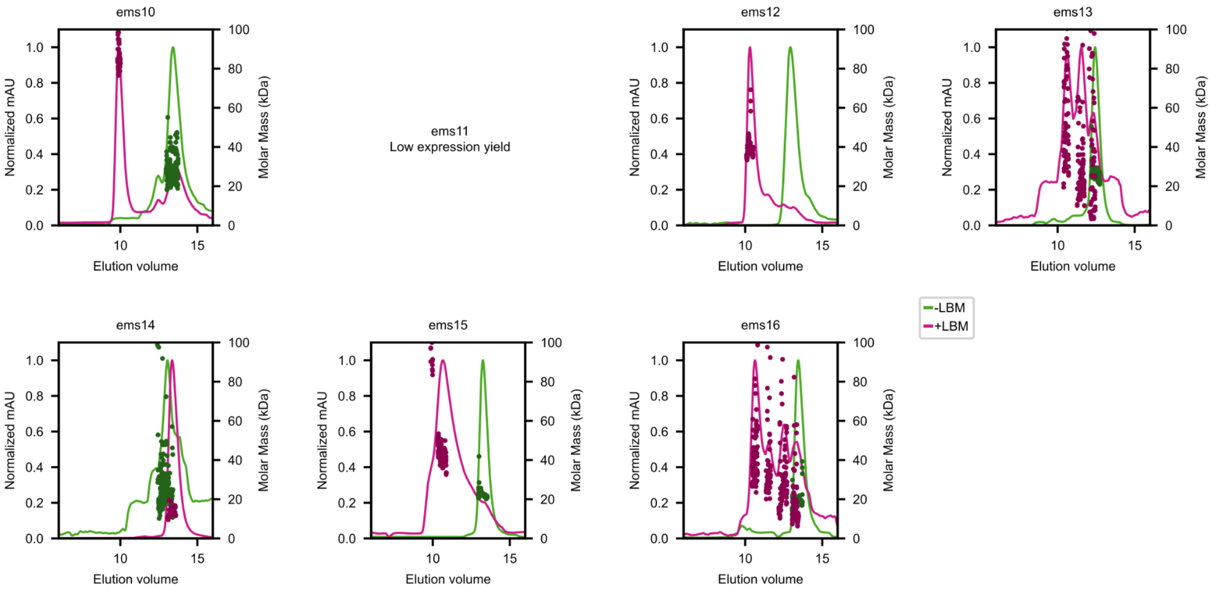

**Fig. S9. Screening of LBM-responsive designs after partial diffusion.** SEC-MALS traces of LBM-responsive designs under apo and LBM-treated (5 µM) conditions. SEC elution profiles before and after treatment are displayed respectively in green and pink, experimentally measured MW shown in darker shade, and theoretical MW shown in dotted lines. All measurements were performed using a Superdex 75 10/300 GL column.

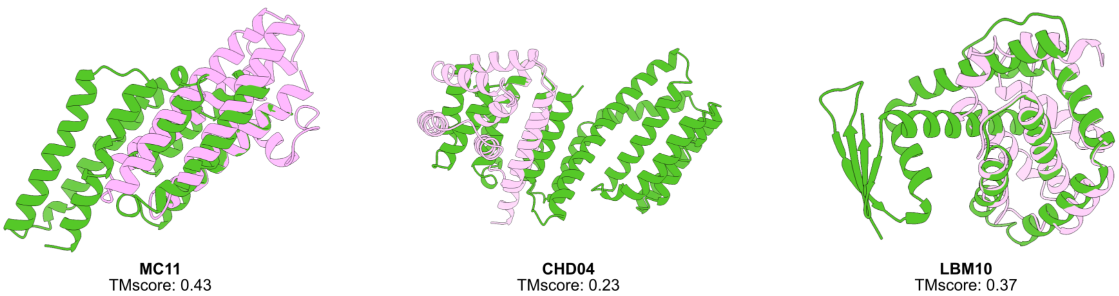

**Fig. S10. Structural novelty of designed metal or small molecule-responsive oligomers.** Closest structural matches (light pink) to the monomeric subunits of each structurally characterized responsive oligomer (green), identified using Foldseek web server. For LBM10, the closest match is Bcl-2, which is one of the binding partners of the LBM-responsive heterodimer from where the design seed has been extracted.

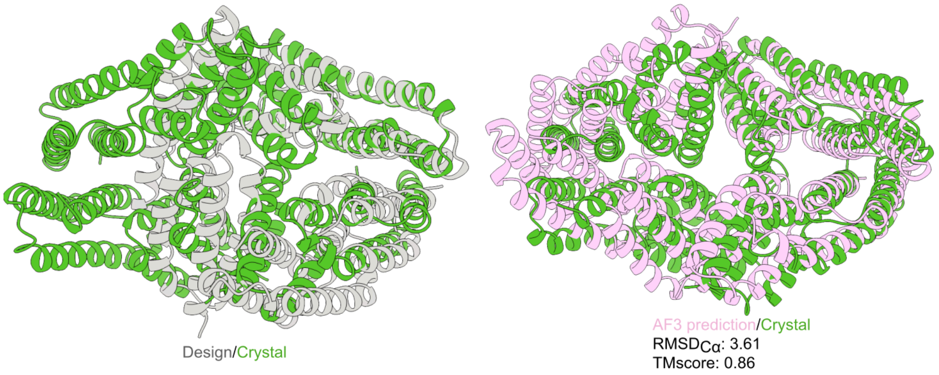

**Fig. S11. Determined crystallographic structure of MC11.** The MC11 crystal structure adopting D2 symmetry (green) is overlayed with the designed C3 symmetric backbone (grey) and the AF3-predicted tetrameric assembly without $Cu^{2+}$ (pink).

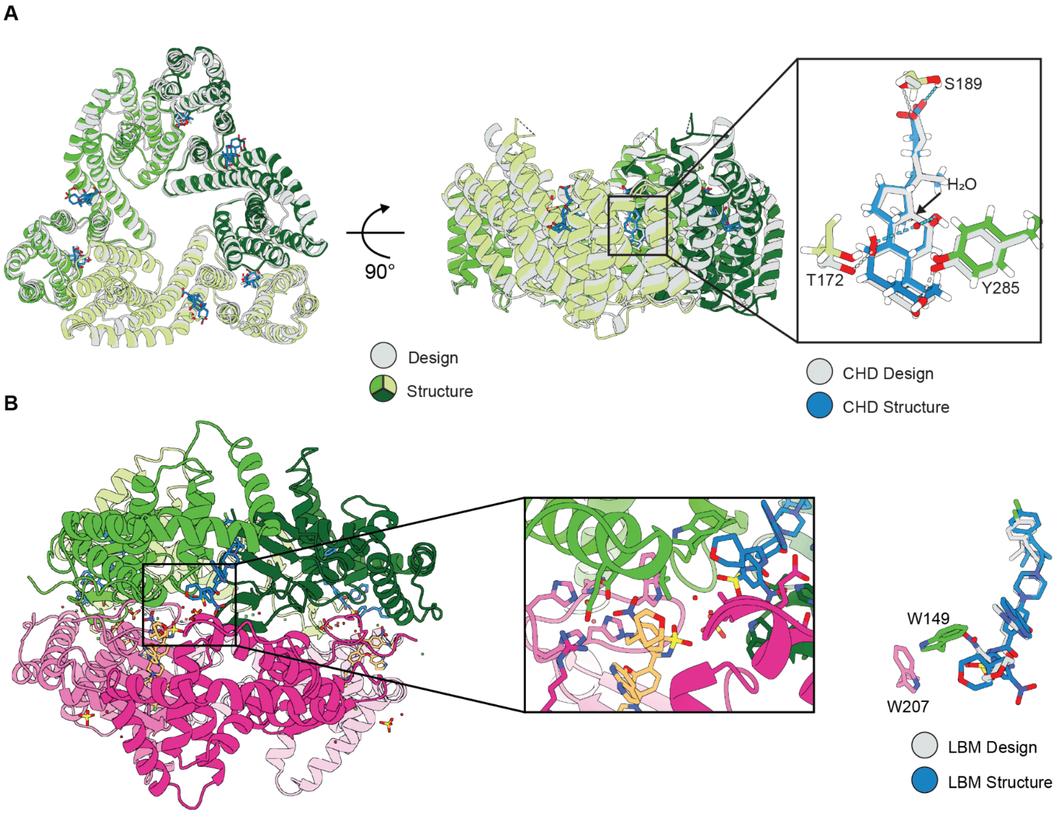

**Fig. S12. Crystal structures of CHD04 and LBM10. (A)** Detailed views of the crystal structure of CHD04 overlaid with the corresponding AF3 prediction. A water molecule mediates hydrogen bonding between two hydroxyl groups of cholic acid, replacing a hydrogen bond predicted between one of the hydroxyl groups and T172. **(B)** Detailed views of the crystal structure of LBM10 dihedral crystal packing. LBM10 crystallized with two non-identical monomers present in the asymmetric unit leading to the dihedral arrangement of two non-equivalent trimers. Crystal packing, along with purification tags present on each trimer contribute to this arrangement and lead to an alternate conformer of the nitrobenzene group present in LBM, relative to the design model.

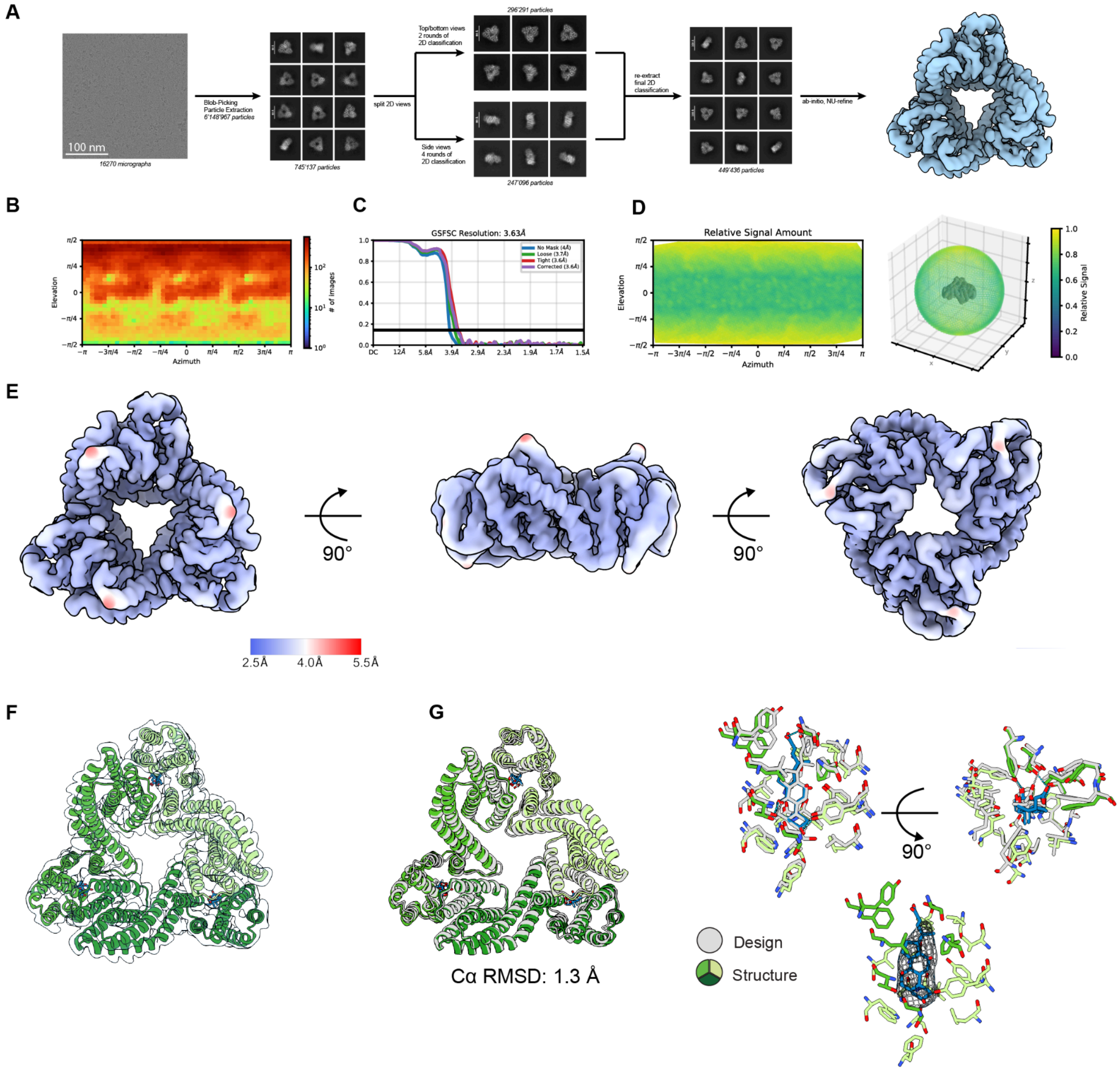

**Fig. S13. Details of Cryo-EM data processing for CHD04 in complex with CHD. (A)** Cryo-EM data processing workflow (see also Methods). **(B)** Viewing direction distribution of final particle set. **(C)** Fourier shell correlation (FSC) plot with threshold of FSC=0.143 shown as solid horizontal line. **(D)** Particle orientation distribution used for the final reconstruction. **(E)** Local resolution estimation of the final density map. **(F)** Refined atomic model of CHD04 fitted into the reconstructed density map. **(G)** Superposition of the cryo-EM structure of CHD04 (colour) and the AF3 prediction (grey) showing close agreement between the experimental structure and design, including side-chain rotamers at the binding interface. Cryo-EM density within 2 Å of the CHD ligand is shown, contoured at 0.0527σ.

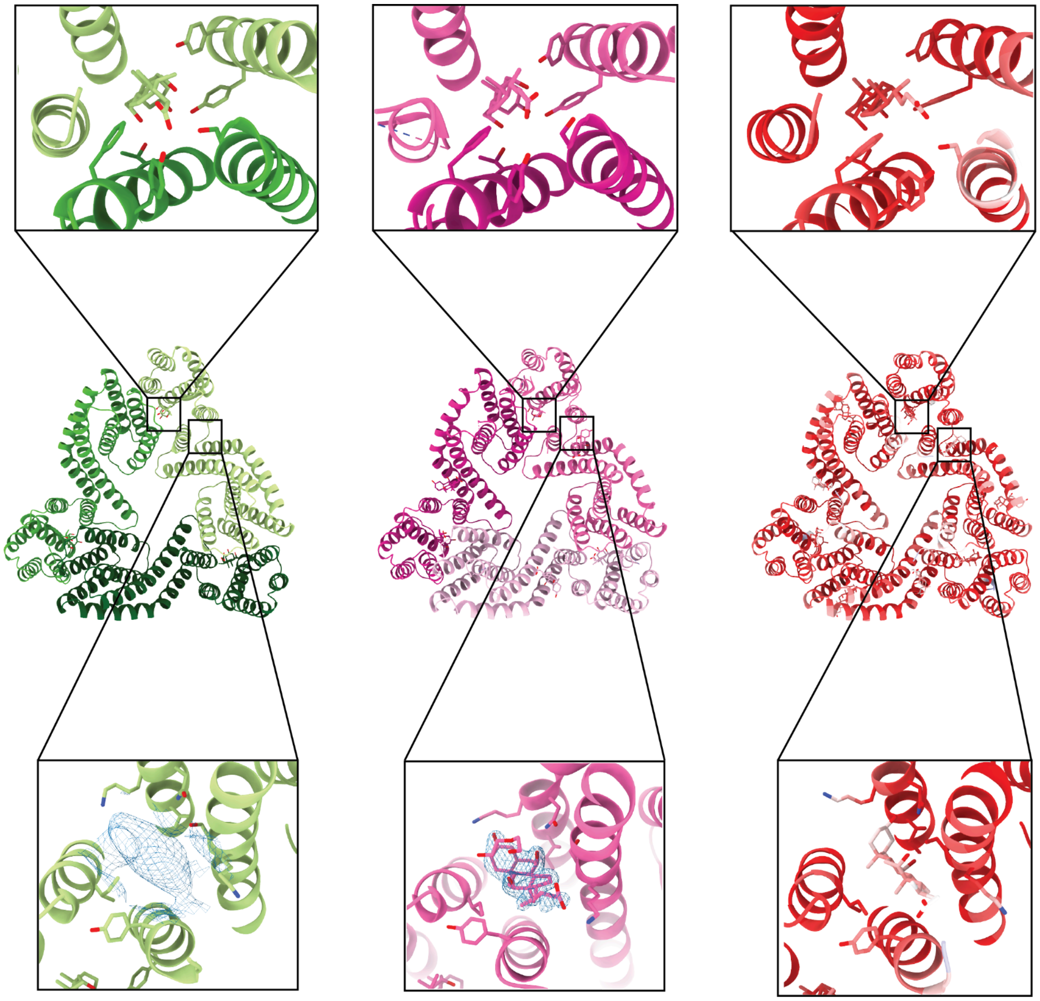

**Fig. S14. CHD binding sites in cryo-EM, crystal and AF3 structures of CHD04.** Cryo-EM (green), crystal (pink) and AF3 (coloured by plDDT, red being high confidence, blue being low confidence) structures of the designed (top) and alternate (bottom) CHD binding sites in CHD04. Both the crystal (Polder omit map, contoured at 3.9σ) and cryo-EM density maps contain additional density at the alternative binding site. However, the local resolution of the cryo-EM map was insufficient to confidently model a CHD molecule. In the AF3 prediction, the alternative CHD binding site was observed only when nine copies of CHD were included in the input.

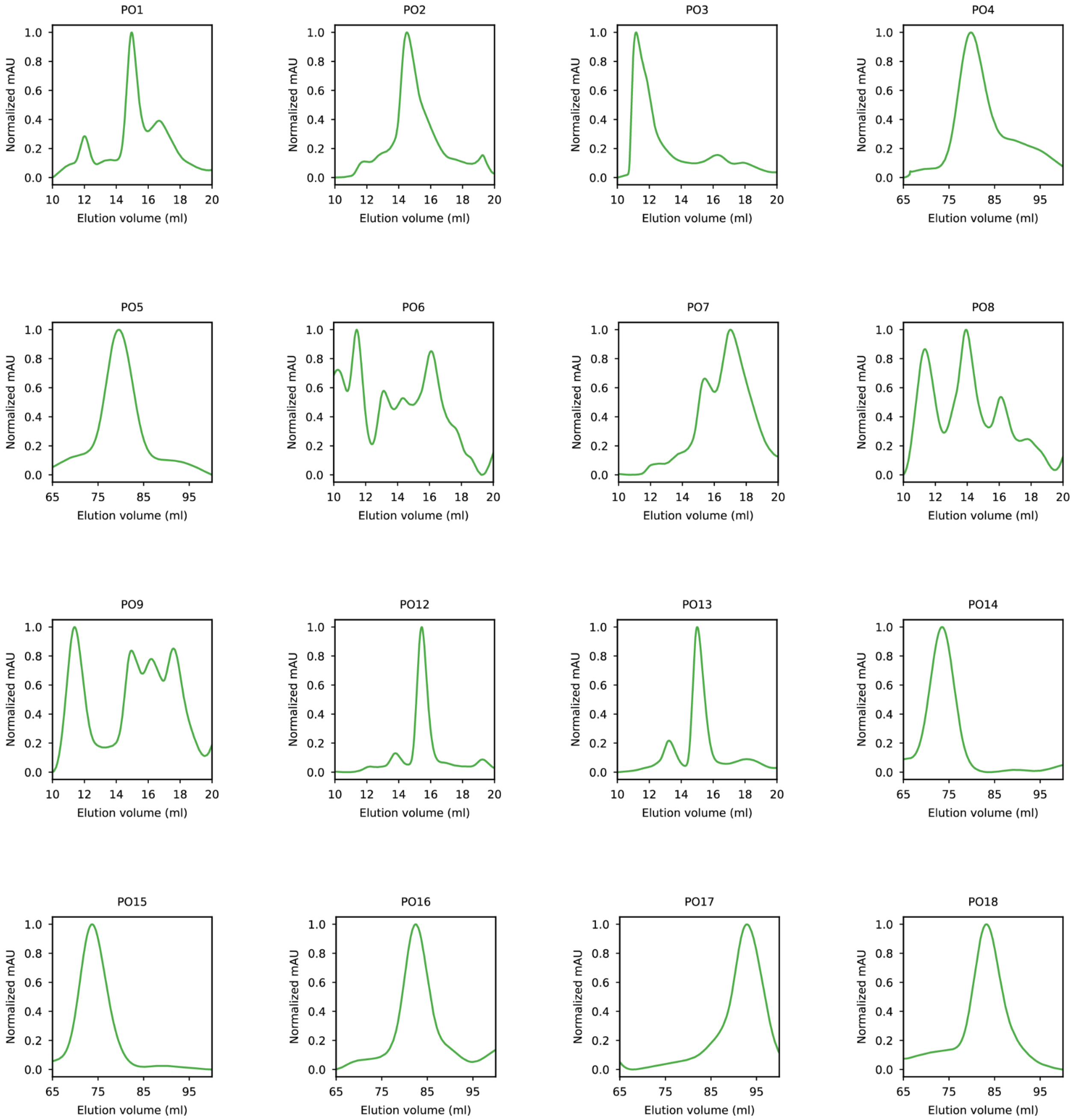
PO1
PO2
PO3
PO4
PO5
PO6
PO7
PO8
PO9
PO12
PO13
PO14
PO15
PO16
PO17
PO18
Normalized mAU
Elution volume (ml)

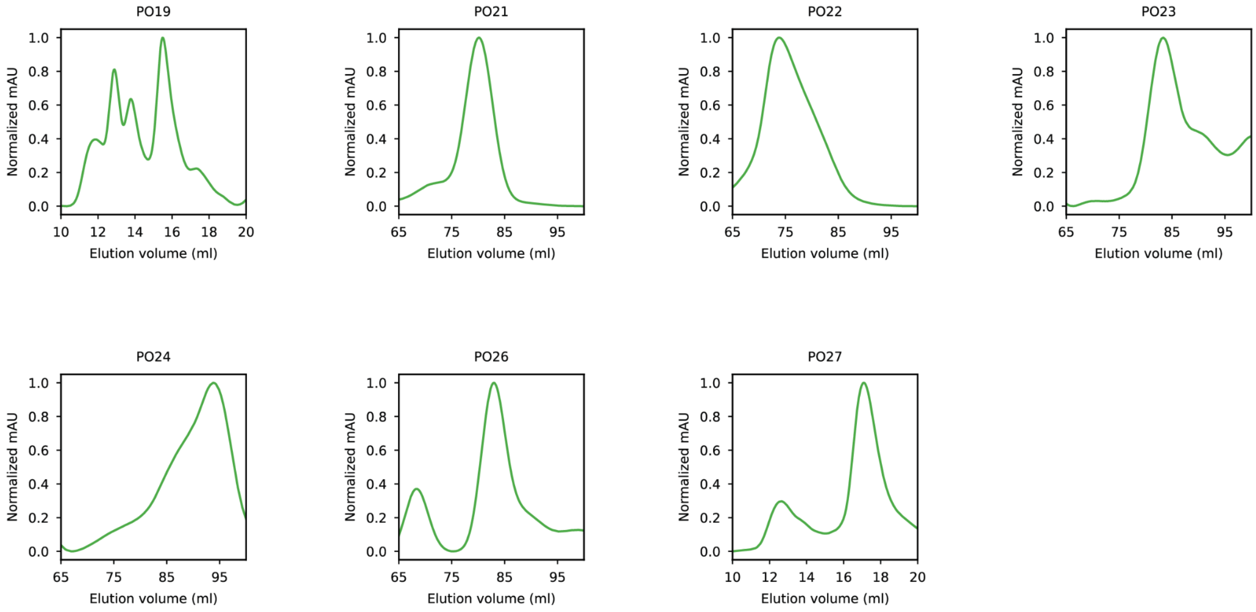

**Fig. S15. Characterization of phosphorylation-responsive oligomers by SEC.** Normalised SEC chromatograms obtained for the phosphorylation-responsive oligomers using a Superdex 200 increase 10/300 GL or Superdex 200 16/600 GL column.

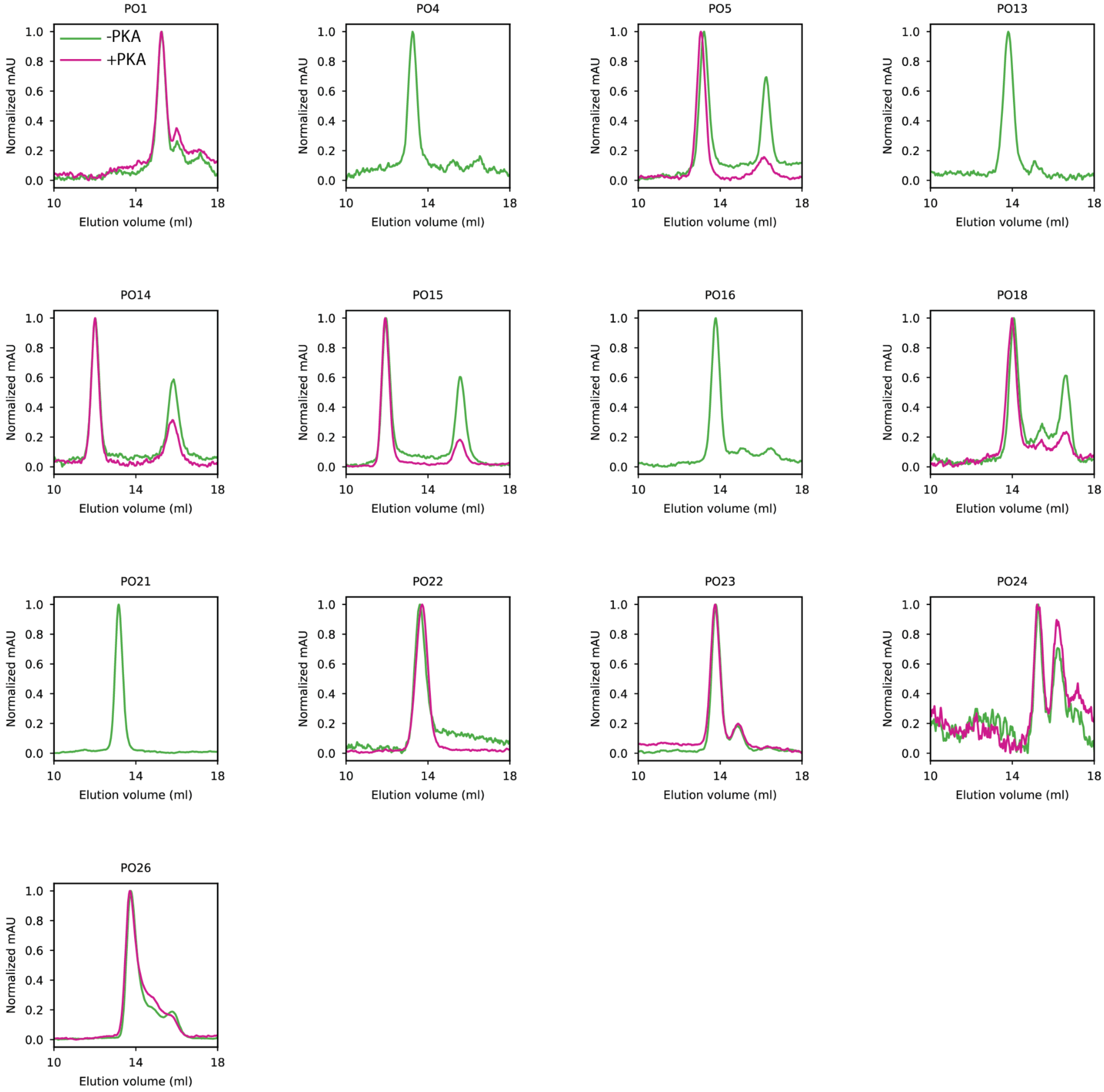

**Fig. S16. SEC profiles of phosphorylation-responsive oligomers treated with or without PKA.** 115 µl of protein at concentrations ranging from 0.4-1 µM (0.01-0.02 mg/ml) were incubated in the presence (pink) or absence (green) of PKA (1:1000 dilution, protein:PKA) prior to injection onto a Superdex 200 10/300 column.

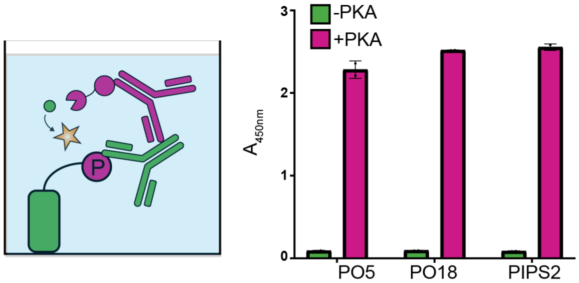

**Fig. S17. Detection of phosphorylation of PO5 and PO18 by ELISA.** (Left) Schematic of the ELISA used to detect phosphorylation of the designed oligomers. Proteins were treated with or without PKA, immobilised on a F96 Maxisorp immunoplate then probed with an anti-RRxpS/T antibody. (Right) ELISA absorbance at 450 nm for PO5, PO18 and the previously reported PIPS2 following incubation in the absence (green) or presence (pink) of PKA. Strong signal was observed only after PKA treatment confirming efficient phosphorylation of the designed proteins. Bars represent mean +/- s.d of 2 independent replicates.

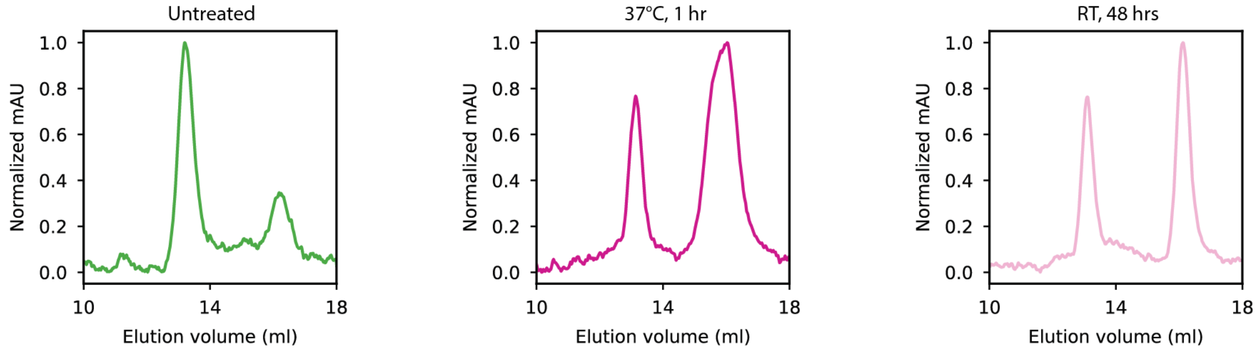

**Fig. S18. SEC analysis of PO5 reveals slow dissociation of oligomeric species into monomeric species.** 115 µl of PO5 at 0.9 µM (0.02 mg/ml) untreated (left), following incubation at 37°C for 1 hour (middle) and following incubation at room temperature (RT) for 48 hours (right) was injected onto a Superdex 200 10/300 column.

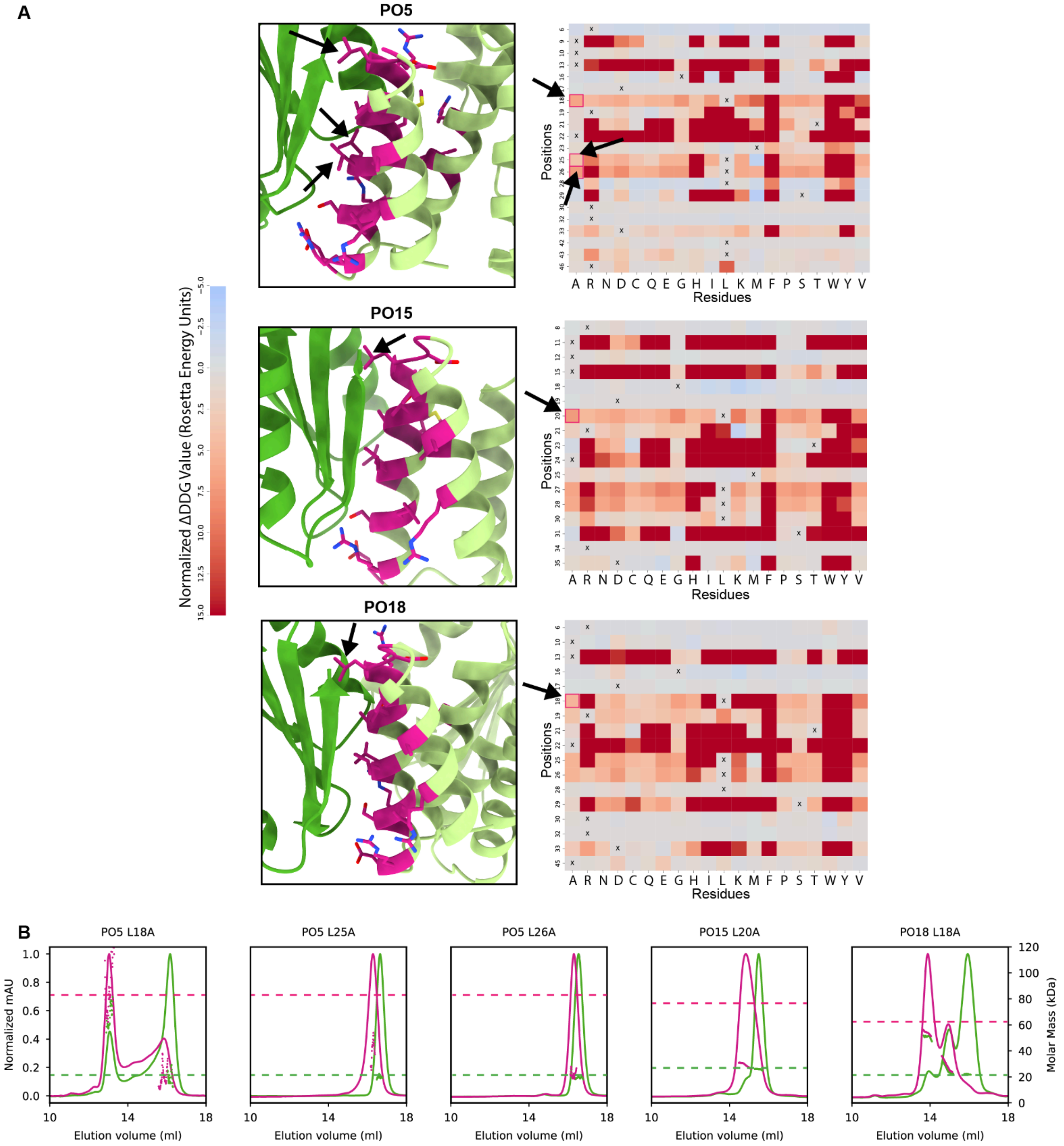

**Fig. S19. *In silico* site-saturation mutagenesis (SSM) of phosphorylation-responsive oligomers. (A)** PO5, PO15 and PO18 were each subjected to an *in silico* SSM to identify interface mutants that are predicted to weaken assembly affinity. Heatmaps representing the Rosetta ΔDDG values for mutations predicted to improve (blue) or worsen (red) binding affinity. Leucine-to-alanine mutations selected for experimental characterization are indicated by black arrows in the model and the corresponding heatmap. **(B)** SEC-MALS profile of the selected interface mutant variants of PO5, PO15 and PO18 following treatment with (pink) or without (green) PKA. Theoretical molecular weights in the oligomeric (pink) and monomeric (green) species are represented by horizontal dashed lines.

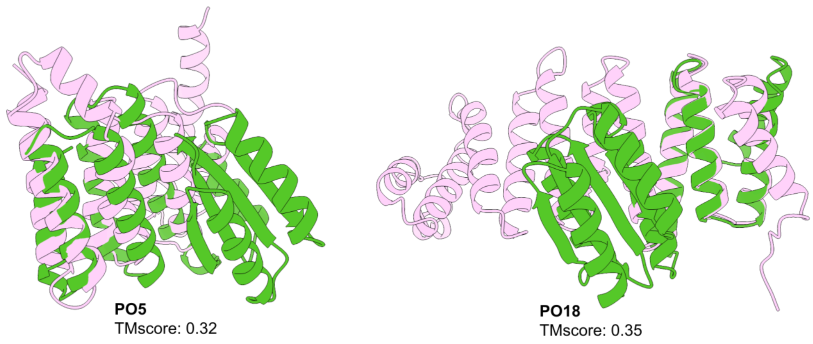

**Fig. S20. Structural novelty of designed PO-responsive oligomers.** Closest structural matches (light pink) to the monomeric subunits of each structurally characterized responsive oligomer (green), identified using Foldseek.

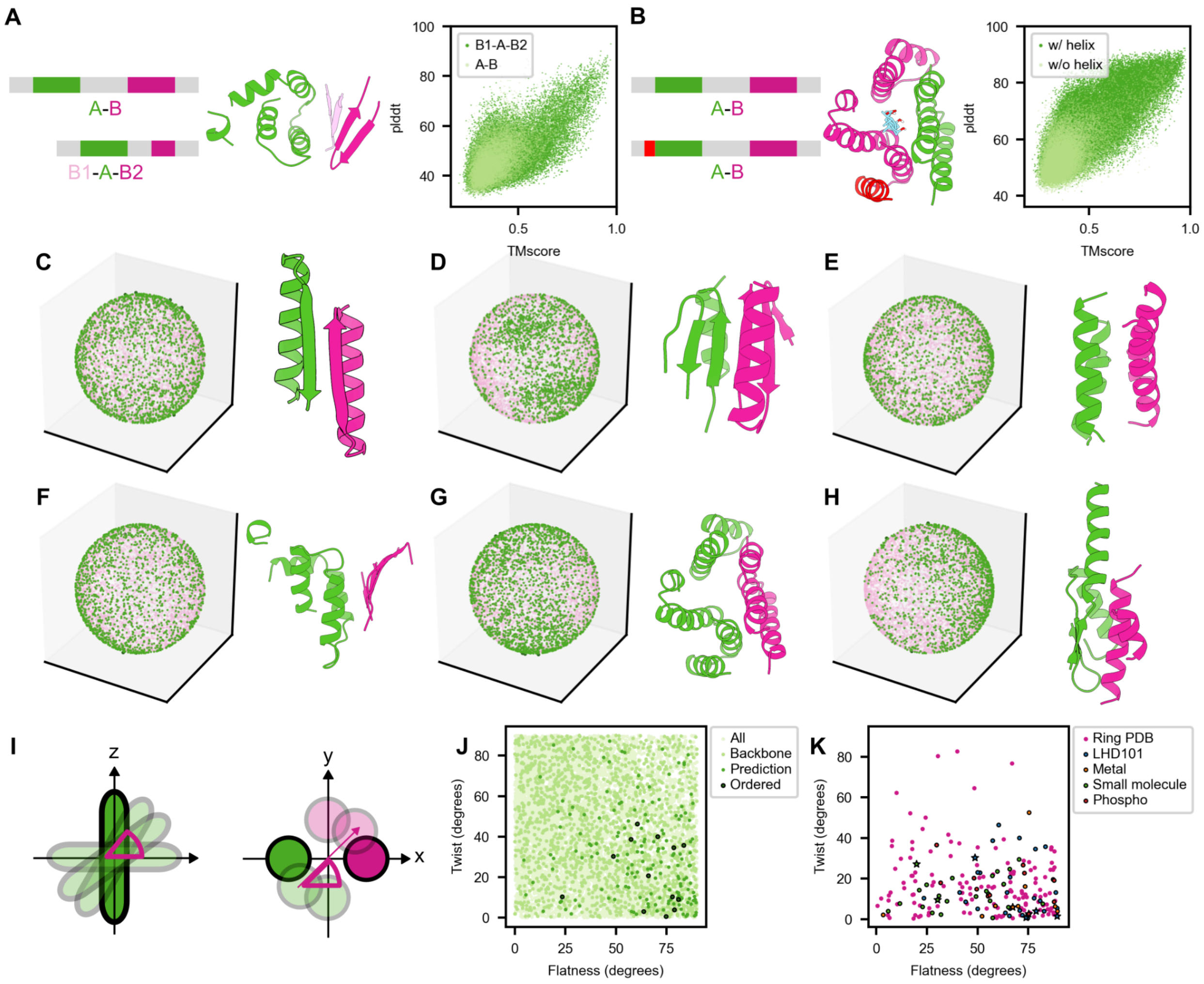

**Fig. S21. Determinants of design success.** Motif re-routing improves designability of backbones for (**A**) LBM-responsive oligomers and (**B**) CHD-responsive oligomers, as evidenced by the presence of sequences with higher AF2 monomer prediction confidence scores (pTM and pldtt) . (**C-H**) Rotational space sampled upon initialization of diffusion trajectories from interface seeds derived from (**C**) LHD101, (**D**) LHD29, (**E**) $Cu^{2+}$-, (**F**) LBM-, (**G**) CHD-, and (**H**) PO-responsive heterodimers. Filtered-out backbones are shown in pink, and backbones retained for sequence design are shown in green. (**I**) Schematic representation of the Flatness and Twist metrics. (**J**) Flatness–Twist space of the LHD101-based obligate trimer designs across different stages of the design pipeline. (**K**) Comparison of Flatness–Twist distributions between cyclic homotrimers from the Protein Data Bank and experimentally validated designs in this study. Structurally validated designs are highlighted with stars.

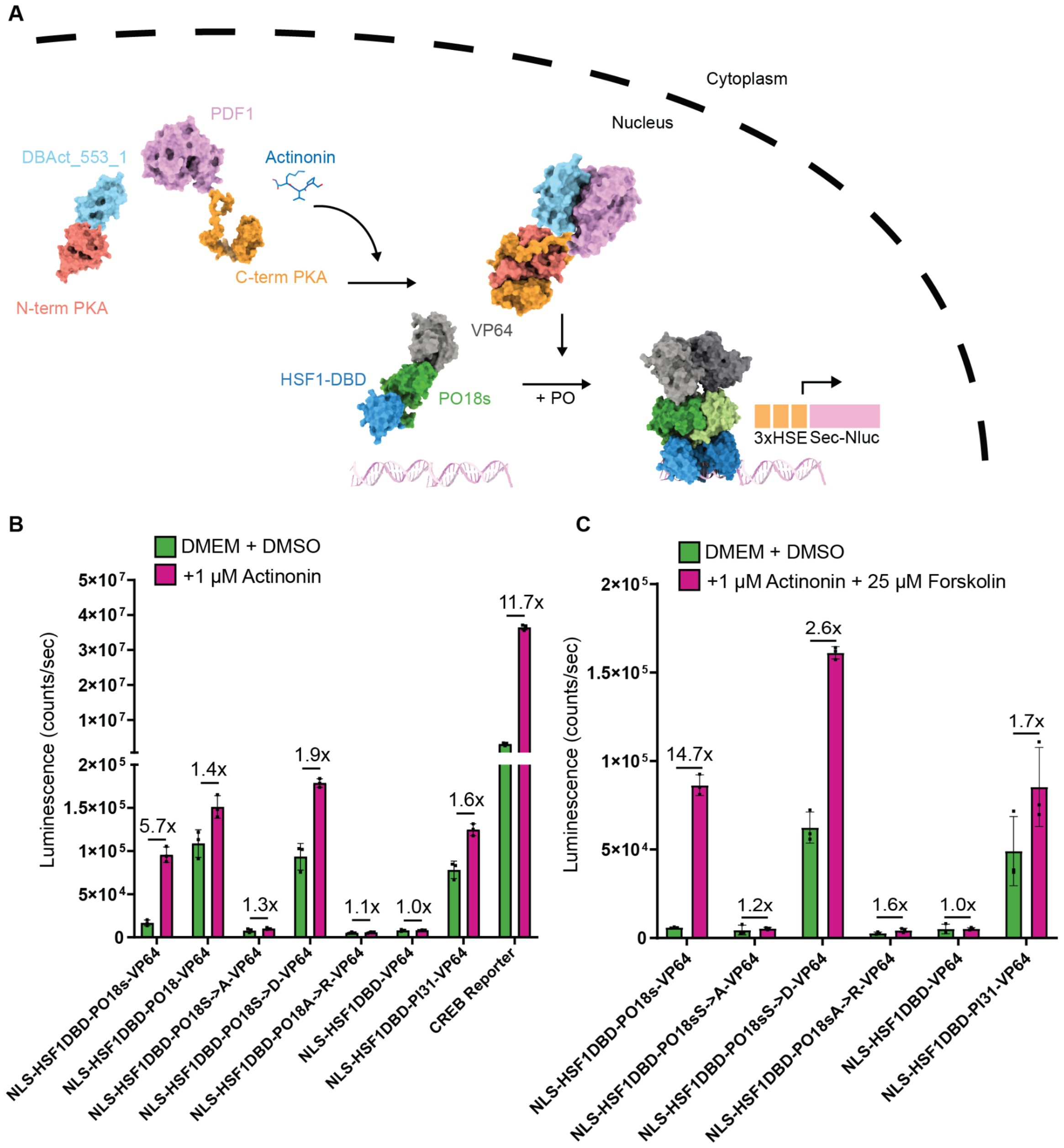

**Fig. S22. Phosphorylation-mediated activation of gene expression through an HSF1 homotrimer. (A)** Schematic of the HSF1 transcriptional reporter assay used in Fig. 5E. Addition of actinonin induces dimerization of an engineered split PKA, leading to the reconstitution of an active kinase in cells. PKA activation drives the phosphorylation-mediated trimerization of the PO18s switch fused to the HSF1 DBD and VP64 transactivation domain, driving the expression of a 3x HSE-driven secreted NanoLuciferase reporter. **(B)** Measured reporter luminescence outputs for the assay shown in Fig. 5E, including the higher-affinity PO18 variant and a CREB reporter as a positive control for PKA activity. **(C)** Measured reporter luminescence outputs as shown in Fig. 5E, however with simultaneous activation of endogenous and engineered split PKA with 25 μM forskolin and 1 μM actinonin respectively. Dual activation led to near 15-fold increases in reporter activity for the PO18s switch. Bars in (B) and (C) represent the mean +/- s.d. of n=3 biological replicates.

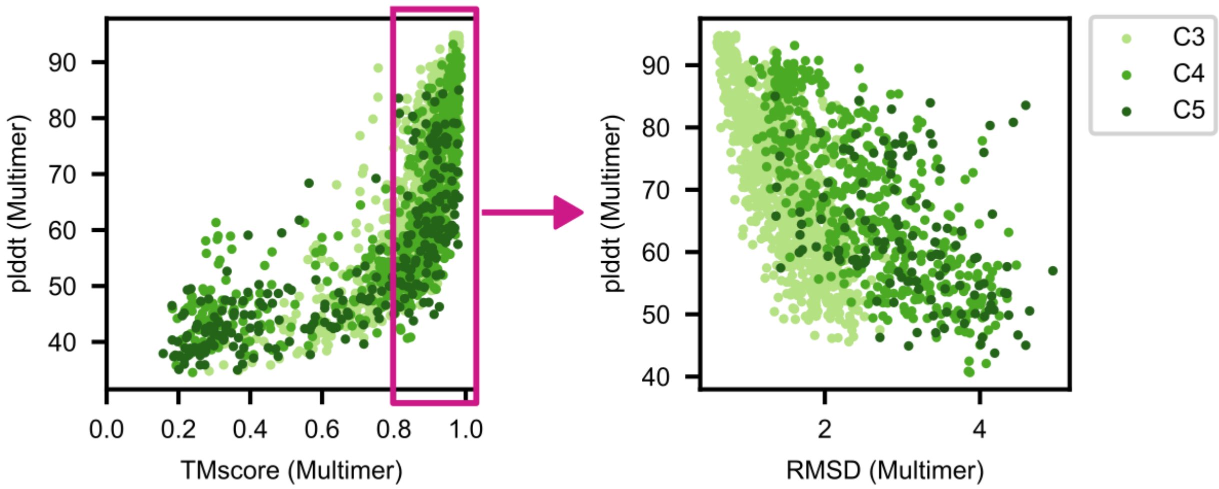

**Fig. S23. Designability analysis of obligate oligomers of different cyclic symmetries designed from a single interface seed.** TM-score, pLDDT and RMSD values for C3-C5 symmetric oligomers designs based on the LHD101-derived interface seed.

### Table S1. Data collection and refinement statistics for crystal structures.

| | **PI25** | **PI31**ǂǂ | **PI56** |
|---|---|---|---|
| **PDB ID** | **31PT** | **31PU** | **31PV** |
| **Data collection** | | | |
| Synchrotron, beamline | EMBL DESY, PETRAIII (P14) | HZB, BESSYII (MX 14.1) | EMBL DESY, PETRAIII (P14) |
| Wavelength (Å) | 0.97626 | 0.9184 | 0.97626 |
| Space group | P 1 | P 1 21 1 | P 21 21 21 |
| *a, b, c* (Å) | 59.789 71.144 71.363 | 169.419 31.699 258.226 | 62.804 163.754 371.976 |
| α, β, γ (°) | 119.851 108.151 91.110 | 90.00 102.08 90.00 | 90.0 90.0 90.0 |
| Resolution (Å)* | 60.677 – 2.881 (3.126 - 2.881) | 47.794 – 3.293 (3.383 – 3.293) | 185.988 – 3.094 (3.346 – 3.094) |
| Directional diffraction limits (Å)[+] | 3.836 2.799 3.015 | 1.812 3.485 3.237 | 3.648 3.284 3.080 |
| Total reflections* | 52072 (2649) | 255946 (12468) | 736240 (36001) |
| Total unique* | 14165 (708) | 39956 (1903) | 54199 (2710) |
| $R_{merge}$* | 0.123 (0.767) | 0.222 (2.110) | 0.206 (1.980) |
| $R_{meas}$* | 0.144 (0.897) | 0.242 (2.293) | 0.214 (2.059) |
| Mean I/σI* | 4.8 (1.2) | 6.2 (0.9) | 10.4 (1.5) |
| Completeness (ellipsoidal, %)[+] | 89.1 (60.3) | 94.6 (67.7) | 93.8 (71.7) |
| Completeness (spherical, %)[+] | 66.1 (15.2) | 93.3 (58.7) | 75.8 (18.4) |
| Multiplicity* | 3.7 (3.7) | 6.4 (6.6) | 13.6 (13.3) |
| CC1/2* | 0.994 (0.589) | 0.997 (0.367) | 0.999 (0.606) |
| **Refinement** | | | |
| Resolution (Å) [#] | 37.12 – 3.10<br>(3.34 – 3.10)ǂ | 47.79 – 3.29<br>(3.38 – 3.29) | 59.5 – 3.09<br>(3.20 – 3.09) |
| Reflections used in refinement[#] | 13561 (1546) | 39816 (1621) | 54147 (502) |
| $R_{work}$[#] | 0.2134 (0.2849) | 0.3141 (0.4088) | 0.1953 (0.4200) |
| $R_{free}$[#] | 0.2491 (0.3293) | 0.3656 (0.4029) | 0.2387 (0.4685) |
| Number of non-hydrogen atoms[#] | 6362 | 18936 | 20843 |
| macromolecules | 6362 | 18936 | 20843 |
| ligands | 0 | 0 | 0 |
| solvent | 0 | 0 | 0 |
| Protein residues | 887 | 2572 | 2980 |
| RMS deviations (bonds)[#] | 0.002 | 0.003 | 0.006 |
| RMS deviations (angles)[#] | 0.66 | 0.86 | 0.95 |
| Ramachandran favored (%)[#] | 97.23 | 96.76 | 98.37 |
| Ramachandran allowed (%)[#] | 2.77 | 3.08 | 1.63 |
| Ramachandran outliers (%)[#] | 0 | 0.16 | 0 |
| Rotamer outliers (%)[#] | 0.51 | 0.64 | 0 |
| Clashscore[#] | 1.05 | 4.02 | 1.85 |
| Average B-factor[#] | 52.26 | 84.79 | 86.90 |
| macromolecules | 52.26 | 84.79 | 86.90 |
| ligands | - | - | - |
| solvent | - | - | - |
| Molprobity Score[#] | **0.95** | **1.39** | **0.95** |
| Criteria used in determination of diffraction limits[+] | local(I/sigI) >= 1.20 | local weighted CC1/2 >= 0.30 | local(I/sigI) >= 1.20 |

[#]as reported by phenix.table_one and phenix.model_vs_data
*as reported by autoPROC
[+]as reported by STARANISO; directional limits (Å) along principal axes of ellipsoid fitted to diffraction cut-off surface; axis vectors: a* [1.00, 0.00, 1.00], b* [0.00, 1.00, 0.00], c* [0.00, 0.00, 1.00]
ǂdata were cut off at 3.10 Å during refinement to maintain an acceptable completeness threshold in the outermost shell
ǂǂdataset exhibits severe tNCS characterized by a Patterson off-origin peak at vector (1/3, 0, 2/3) with a p-value of $2.200×10^{-4}$. tNCS corrections were applied throughout molecular replacement and refinement.

### Table S1. Data collection and refinement statistics for crystal structures. (Continued)

| | PI57 | MC11 |
|---|---|---|
| **PDB ID** | **31PW** | **31PX** |
| **Data collection** | | |
| Synchrotron, beamline | HZB, BESSYII (MX 14.1) | EMBL DESY, PETRAIII (P14) |
| Wavelength (Å) | 0.9184 | 0.97626 |
| Space group | C 2 2 21 | P 31 21 |
| *a, b, c* (Å) | 92.19 113.24 122.44 | 80.027 80.027 256.582 |
| α, β, γ (°) | 90.0 90.0 90.0 | 90.0 90.0 120.0 |
| Resolution (Å)* | 46.5 – 3.24 (3.44 – 3.24) | 85.527 – 2.26 (2.387 – 2.26) |
| Directional diffraction limits (Å)$^+$ | - | 2.296 2.296 2.346 |
| Total reflections* | 135200 (21434) | 805884 (42273) |
| Total unique* | 19658 (3171) | 39832 (1992) |
| $R_{merge}$* | 0.174 (2.516) | 0.127 (2.654) |
| $R_{meas}$* | 0.188 (2.727) | 0.131 (2.719) |
| Mean I/σI* | 8.74 (0.72) | 13.5 (1.4) |
| Completeness (ellipsoidal, %)$^+$ | - | 92.9 (48.7) |
| Completeness (spherical, %)$^+$ | 99.9 (99.1)* | 87.0 (29.2)$^+$ |
| Multiplicity* | 12.9 (15.7) | 20.2 (21.2) |
| CC1/2* | 0.999 (0.308) | 1.000 (0.589) |
| **Refinement** | | |
| Resolution (Å) $^\#$ | 41.57 – 3.24 (3.41 – 3.24) | 42.76 – 2.26 (2.34 – 2.26) |
| Reflections used in refinement$^\#$ | 19617 (2815) | 39821 (573) |
| $R_{work}$$^\#$ | 0.2263 (0.3965) | 0.2081 (0.3048) |
| $R_{free}$$^\#$ | 0.2743 (0.4329) | 0.2384 (0.4513) |
| Number of non-hydrogen atoms$^\#$ | 3884 | 5022 |
| macromolecules | 3884 | 4926 |
| ligands | 0 | 35 |
| solvent | 0 | 61 |
| Protein residues | 560 | 682 |
| RMS deviations (bonds)$^\#$ | 0.003 | 0.009 |
| RMS deviations (angles)$^\#$ | 0.66 | 1.12 |
| Ramachandran favored (%)$^\#$ | 96.38 | 98.81 |
| Ramachandran allowed (%)$^\#$ | 3.62 | 1.19 |
| Ramachandran outliers (%)$^\#$ | 0 | 0 |
| Rotamer outliers (%)$^\#$ | 0 | 0.70 |
| Clashscore$^\#$ | 9.39 | 1.76 |
| Average B-factor$^\#$ | 131.28 | 71.03 |
| macromolecules | 131.28 | 71.01 |
| ligands | 0 | 87.32 |
| solvent | 0 | 63.41 |
| Molprobity Score$^\#$ | **1.74** | **0.93** |
| Criteria used in determination of diffraction limits$^+$ | *CC(1/2) >= 0.25 | $^+$local(I/sigI) >= 1.20 |

$^\#$as reported by phenix.table_one and phenix.model_vs_data
*as reported by autoPROC
$^+$as reported by STARANISO; directional limits (Å) along principal axes of ellipsoid fitted to diffraction cut-off surface; axis vectors: a* [1.00, 0.00, 1.00], b* [0.00, 1.00, 0.00], c* [0.00, 0.00, 1.00]

### Table S1. Data collection and refinement statistics for crystal structures. (Continued)

| | LBM10 | CHD04 | PO5 |
|---|---|---|---|
| **PDB ID** | **31VQ** | **31VR** | **31VP** |
| **Data collection** | | | |
| Synchrotron, beamline | ESRF, MASSIF-1 (ID30A-1) | ESRF, MASSIF-1 (ID30A-1) | ESRF, MASSIF-1 (ID30A-1) |
| Wavelength (Å) | 0.96546 | 0.96546 | 0.96546 |
| Space group | H 32 | H 3 | P 4 |
| *a, b, c* (Å) | 126.688 126.688 177.438 | 149.292 149.292 46.299 | 74.488 74.48 77.942 |
| α, β, γ (°) | 90.00 90.00 120.00 | 90.0 90.0 120.0 | 90.0 90.0 90.0 |
| Resolution (Å)* | 19.89 – 2.73 (2.96 – 2.73) | 43.59 – 2.10 (2.17 – 2.10) | 18.62 – 1.81 (1.95 – 1.81) |
| Directional diffraction limits (Å)⁺ | 2.76, 2.76, 3.32 | 1.93, 1.93, 2.55 | 1.80, 1.80, 2.41 |
| Total reflections* | 160744 (7470) | 178884 (8545) | 114929 (7229) |
| Total unique* | 11054 (553) | 18021 (901) | 27234 (1362) |
| $R_{merge}$* | 0.339 (2.882) | 0.122 (1.468) | 0.077 (1.437) |
| $R_{meas}$* | 0.351 (2.995) | 0.129 (1.554) | 0.087 (1.592) |
| Mean I/σI* | 9.3 (1.4) | 9.9 (2.0) | 7.9 (1.5) |
| Completeness (ellipsoidal, %)⁺ | 93.2 (58.1) | 90.3 (79.0) | 91.2 (56.0) |
| Completeness (spherical, %)⁺ | 75.0 (17.9) | 79.8 (43.2) | 70.5 (18.5) |
| Multiplicity* | 14.5 (13.5) | 9.9 (9.5) | 4.2 (5.3) |
| CC1/2* | 0.993 (0.506) | 0.998 (0.570) | 0.997 (0.660) |
| **Refinement** | | | |
| Resolution (Å) # | 19.89 – 2.73 (3.01 - 2.73) | 43.59 – 2.10 (2.18 – 2.10) | 18.62 – 1.81 (1.93 – 1.81) |
| Reflections used in refinement# | 11051 (792) | 18002 (1050) | 27190 (977) |
| $R_{work}$# | 0.2357 (0.2981) | 0.2207 (0.2976) | 0.2185 (0.3238) |
| $R_{free}$# | 0.2747 (0.3504) | 0.2496 (0.2955) | 0.2483 (0.3388) |
| Number of non-hydrogen atoms# | 3636 | 2468 | 2776 |
| macromolecules | 3492 | 2398 | 2769 |
| ligands | 137 | 65 | 0 |
| solvent | 7 | 5 | 7 |
| Protein residues | 412 | 292 | 363 |
| RMS deviations (bonds)# | 0.007 | 0.004 | 0.016 |
| RMS deviations (angles)# | 0.77 | 0.77 | 1.48 |
| Ramachandran favored (%)# | 95.83 | 98.61 | 96.38 |
| Ramachandran allowed (%)# | 4.17 | 1.39 | 3.62 |
| Ramachandran outliers (%)# | 0.00 | 0.00 | 0.00 |
| Rotamer outliers (%)# | 0.56 | 0.00 | 2.18 |
| Clashscore# | 1.69 | 0.00 | 0.88 |
| Average B-factor# | 59.78 | 46.14 | 44.54 |
| macromolecules | 60.32 | 46.32 | 44.56 |
| ligands | 47.37 | 40.69 | - |
| solvent | 32.43 | 32.18 | 33.73 |
| Criteria used in determination of diffraction limits⁺ | local(I/sigI) >= 1.20 | local(I/sigI) >= 1.20 | local(I/sigI) >= 1.20 |

#as reported by phenix.table_one and phenix.model_vs_data
*as reported by autoPROC
⁺as reported by STARANISO; directional limits (Å) along principal axes of ellipsoid fitted to diffraction cut-off surface; axis vectors: a* [1.00, 0.00, 1.00], b* [0.00, 1.00, 0.00], c* [0.00, 0.00, 1.00]

**Table S2. Data collection and refinement statistics for Cryo EM.**
Type or paste caption here. Create a page break and paste in the table above the caption.

| | **CHD04**<br>PDB ID: 31SW<br>EMDB-58650 |
|---|---|
| **Collection & Processing** | |
| Magnification | 190,000 |
| Voltage (kV) | 200 |
| Electron exposure (e–/Å$^2$) | 50 |
| Defocus range (μm) | -1.0 - 2.4 |
| Pixel size (Å) | 0.72 |
| Symmetry imposed | C3 |
| Number of micrographs | 16,270 |
| Initial particle images (no.) | 6,148,967 |
| Final particle images (no.) | 449,436 |
| Map resolution (Å) | 3.63 |
| FSC threshold | 0.143 |
| **Real-Space Refinement** | |
| Model resolution (Å)* | 3.57 / 3.62 |
| FSC threshold | 0.143 |
| Sharpening B-factor (Å$^2$) | -50 |
| Model composition | |
| Chains | 3 |
| Atoms | 7353 |
| Residues | 882 |
| Ligands | 3 |
| B-factors (Å$^2$) | |
| Protein** | 125.35/257.92/176.44 |
| Ligand | 156.67 |
| R.M.S. deviations | |
| Bond lengths (Å) | 0.005 |
| Bond angles (°) | 0.799 |
| Validation | |
| MolProbity score | 1.25 |
| Clashscore | 4.79 |
| Poor Rotamers (%) | 0.00 |
| Ramachandran plot | |
| Favored (%) | 99.32 |
| Allowed (%) | 0.68 |
| Disallowed (%) | 0.00 |
| Model vs Data | |
| CC (mask) | 0.86 |
| CC (box) | 0.88 |
| CC (peaks) | 0.75 |
| CC (volume) | 0.86 |
| CC (ligands) | 0.84 |

** d FSC Model 0.143 Masked / Unmasked from Phenix*

*** min/max/mean*

**Table S3. Plasmids designed and used in this study.**

| Plasmid | Description | Reference |
|---|---|---|
| LM627 | M-AGGA-promoter-ccdb-TTCC-SNAC-HIS | Qian et al. |
| LM1371 | MS-his-AGGA-promoter-ccdb-TTCC | Qian et al. |
| LM1487 | his-mScarletI-AGGA-promoter-ccdb-TTCC-his | Qian et al. |
| MA_nG | his-sfGFP-AGGA-promoter-ccdb-TTCC-his | This study |
| MA_cM | MSG-AGGA-promoter-ccdb-TTCC-mScarletI-his | This study |
| MA_cG | MSG-AGGA-promoter-ccdb-TTCC-sfGFP-his | This study |
| pMI287 | pCMV-NLS-HSF1DBD-PO18s-VP64 | This study |
| pMI392 | pCMV-NLS-HSF1DBD-PO18-VP64 | This study |
| pMI389 | pCMV-NLS-HSF1DBD-PO18s_S->A-VP64 | This study |
| pMI390 | pCMV-NLS-HSF1DBD-PO18s_S->D-VP64 | This study |
| pMI391 | pCMV-NLS-HSF1DBD-PO18s_A->R-VP64 | This study |
| pMI303 | pCMV-NLS-HSF1DBD-VP64 | This study |
| pMI302 | pCMV-NLS-HSF1DBD-PI31-VP64 | This study |
| pMI282 | pCMVmin-3xHSE-SecNluc_reporter | This study |
| pLS1042 | pCMVmin-CRE-SecNluc_reporter | Buckley et al. |
| pSB490 | pCMV-PDF1-Cterm_PKA | This study |
| pSB491 | pCMV-Nterm_PKA-DBAct553_1 | This study |

**Table S4. Protein sequence of designs experimentally validated in this study.**

| Design name | Amino acid sequence |
|---|---|
| PI13 | TWQWVLINISEKTQERLIKALEELVSFAQELYKAMGLETELSLSVEKDDGVLHIRVTCPDWFAREIHKVAKLILEVAAAERKGADKEELEEKLKKGIQEVKWKLFEHLFTPSSIFLLSNVDTDEAMEKVVEIFEKVLGKKCKVYQFTEDKVFLLCEPGIAVRIEKDDGFLTIEVKNLSEERLREIAKALQLIVDV |
| PI14 | TWQWVLINISEETQERLLKAVEELLSFAQRLYKAMGDKTELSISFEKDDGVLHIRVECPDWLAREIHKVAKLILEVAAAERRGADREELEERYRRGFEEVKWNLFKHLFTPSSIFLLSNVDLDEATEKVVEIIEEVLGKKCEVYKLTEDKTVLLCEPGIAVAIEKDDGFLTIEVKNLSEERLREIAKALQLIVDV |
| PI15 | TWQWVLINISEETQKRLIEALRDLFDFYQKLLKNMGSKEKVSLSIEKDDGVLHIRVTCSDFFAREIHKVAKLILEVAAAERKGADKAELEERLEKGISKVKWNLFKSLFTPSSIFLLSNVDLDEAMEKVVEIIEEVLGKKCEVFQFEEDKVFLLCEPNVSVSIEKDDGFLTIEVKNLSEERLREIAKALQLIVDV |
| PI16 | TWQWVLINISKETEEELIKSIEELHEIWKKLAKNRGEELGSISIEKDDGVLHIRVENISDEFAREIHKVAKLILEVAAAERKGESKEKVEKYREERQWILDRLKVLAEVSEETGVKVTLEEVEVEPPSSIFLLSNVDEIEALKAALRLVVELFPDAEITLSDSNRVVVRTRDGKVTVTIEKDDGFLTIEVKNLSEERLREIAKALQLIVDV |
| PI17 | LHESQQEELLELVLRAAELAGVRVRIRFKGDEVEVEIVDYGVRVRITGENYEEALEFVLKVLAGEPAIYIEKEGDVLKVYVTGPMSPEQIKEIIDKAKELGAKIEVHLAGLHIKQQRQLYRDVREAAKKAGVEVEIEVE |
| PI18 | LHESQQEELLELVLRAAELAGVRVRIRFKGDEVEVEIVDYGVRVRLTGENFEEALEFVLAVLAGEPYLYIEEEGDELVVIVTGPMSKEEIEKIIEKAKELGKKIRVRLAGLHIKQQRQLYRDVREAAKKAGVEVEIEVE |
| PI19 | LHESQQEELLELVLRAAELAGVRVRIRFKGDRVTVETFESDLEKTEAFFDRALELARERGGVAVLASGVTLTREQMERLYARMREAEGVSFVLSLVLSPEGHPILLEMIEELNVLQVSVTLRGAHIKQQRQLYRDVREAAKKAGVEVEIEVE |
| PI20 | LHESQQEELLELVLRAAELAGVRVRIRFKGDTVTIETFGEDAEKTMRFWETAIELARERGGVAALASGIPVSEEQMRWLYERMKEAEGVSFVLSLAILPSCHPILIKMIEELNVLQVRVRLRGAHIKQQRQLYRDVREAAKKAGVEVEIEVE |
| PI21 | LHESQQEELLELVLRAAELAGVRVRIRFKGDEVTVEVTLEDAERAAEFLSTVYEGLKKLGIKCTLHVTCSADPALVAVACEQILKLIEECGCHVVLECPLRPEQLPTILDLIEKAKGPLLLRLRIGAHIKQQRQLYRDVREAAKKAGVEVEIEVE |
| PI22 | LHESQQEELLELVLRAAELAGVRVRIRFKGDEVEVEVELEDAEKARLFLSTLYEGLKERGIRCTLHVRVSEDPELVRVACEQILRLMKECGCRVVLEAPLTPEQLPDILRLIEEATGELLLRLRIGAHIKQQRQLYRDVREAAKKAGVEVEIEVE |
| PI23 | LHESQQEELLELVLRAAELAGVRVRIRFKGDEVEVEVELEDAEKAKLFLSTLYEGLKEKGIKCTLHITLSADAELVKVGCEQILKLINECGCKVHLTAPLRPEHLPHVLELLEEAKGELHLHLLIGAHIKQQRQLYRDVREAAKKAGVEVEIEVE |
| PI24 | LHESQQEELLELVLRAAELAGVRVRIRFKGDEVEIEFEGALRLLTPEARETGIRLLALCLEAGIELRCTVVVRRAEDVRALAEVGYRGRLAVEAPTAEEMLELHRLARELGFDEVVILLRGHIKQQRQLYRDVREAAKKAGVEVEIEVE |
| PI25 | LHESQQEELLELVLRAAELAGVRVRIRFKGDEMTIEFEGPLYALTPEAQETIIELVRKLLEAGLVLDETIVVETEETVRALAEVGFRGKLAVTSKTAEEMLRKYKLAKELGFDEVVILLRGHIKQQRQLYRDVREAAKKAGVEVEIEVE |
| PI26 | LHESQQEELLELVLRAAELAGVRVRIRFKGDEVEIEFEGALELLSPEAQETGIRFLALCLEAGIECGFEVVARTAETIRRLAEVGFRGALAVTAPTAEEMLENYKLAKELGFDRVVIKLGGHIKQQRQLYRDVREAAKKAGVEVEIEVE |
| PI27 | LHESQQEELLELVLRAAELAGVRVRIRFKGDEVEIEIAEGDEEEAIIECVELIGEACQALGLQVHITINTYLSADAATKICEACQKHGLRATIHVDVKPEDLDSVKKMLEAAKTVHIHIRIGAHIKQQRQLYRDVREAAKKAGVEVEIEVE |
| PI28 | LHESQQEELLELVLRAAELAGVRVRIRFKGDEVEIEFAEGESEEALIEAVKLIGEAAQQLGLRLHIRIRTLL |

| | |
|---|---|
| | SPEAAAEICKVLQEYGLEAEIRIELRPEDMEALRETLEAAKTVHLHIRIGAHIKQQRQLYRDVREAAKKAGVEVEIEVE |
| PI29 | LHESQQEELLELVLRAAELAGVRVRIRFKGDEVEIEIAEGEPEEAIRECVRLIGEACRELGLRLRLTIHTYLSCEAAREICEILEEYGLEAELHVRLTPEELEALKEMLKAAKTVRLHIRIGAHIKQQRQLYRDVREAAKKAGVEVEIEVE |
| PI30 | LHESQQEELLELVLRAAELAGVRVRIRFKGDEMEVEIYVEEDIAPEELTERMIAFFEELVRLARERGLEEAVRSLTFTVVMERNVTPEDCKAIEEIGKAAKKAGLKVKFRVRITGGHIKQQRQLYRDVREAAKKAGVEVEIEVE |
| PI31 | LHESQQEELLELVLRAAELAGVRVRIRFKGDEIEVELYVEPEIEPEEFTERMKKFFEELKKLAEERGLVEEVKKITFTIVLERNVRPEDCEAIEEVGKEAKKAGLKVKFVVRITGYHIKQQRQLYRDVREAAKKAGVEVEIEVE |
| PI32 | LHESQQEELLELVLRAAELAGVRVRIRFKGDEMEVELYVDESESPEEFTERMIRFIEELVRLAKERGLEEEVKSLKFTIVINRNVTPEDCEAIEKVGEAFKKAGLKAKFEVRIRGGHIKQQRQLYRDVREAAKKAGVEVEIEVE |
| PI33 | LHESQQEELLELVLRAAELAGVRVRIRFKGDEVTISIMTGGGAEATARLPSELTLEQLRALIERARQLGLRLALIGELESGEKVLLVAVADAAAFEELVRELVEEEGVKKVMVVIGGHHIKQQRQLYRDVREAAKKAGVEVEIEVE |
| PI34 | LHESQQEELLELVLRAAELAGVRVRIRFKGDEVEIEVEHEDIARALGCLELIAEECRRRGLEAFARQVEEQRRRLLEEERPRIEARKAEERKKGVDFYLVTDENIKEVAEEIIKSPRPVRVETPHIKQQRQLYRDVREAAKKAGVEVEIEVE |
| PI35 | LHESQQEELLELVLRAAELAGVRVRIRFKGDEVELTLGLEVEDAEEAVRRVREACERLQQILDDLGVELKRLEFELVITRPVLTVEQLTEIFEALKALGARARVRLRGLHIKQQRQLYRDVREAAKKAGVEVEIEVE |
| PI36 | LHESQQEELLELVLRAAELAGVRVRIRFKGDEVSITLGLEVEDAEEAVRALKECVERLKRILEELGVEVKELVAEIVIRVPNLTVEQLTEMFEALKELGAKARVRLEGLHIKQQRQLYRDVREAAKKAGVEVEIEVE |
| PI37 | LHESQQEELLELVLRAAELAGVRVRIRFKGDKVSLTAGLEVEDAEEAVRRVREFCRRLKEILERLGVEVEELRFELVIRRPSLTVEQLTEIFKALKELGAKARVRLEGLHIKQQRQLYRDVREAAKKAGVEVEIEVE |
| PI38 | LHESQQEELLELVLRAAELAGVRVRIRFKGDEVRIELEVRTAAECAPARALLEMILSTLKEAGQKVEVELRLGPGVDAAEVEAMVRSILEAGATLTLEAELGTPEAVEAVKAVMELAPEAKIRVRIGAHIKQQRQLYRDVREAAKKAGVEVEIEVE |
| PI39 | LHESQQEELLELVLRAAELAGVRVRIRFKGDEVELELEIETAEQCPAAAELLRMALSRLREAGQRVRVHLRLGAGVRAEEVAEIVRALLEAGAELRVEMHVGTPEAVEAVKAVMELAPEAEIRVRIGAHIKQQRQLYRDVREAAKKAGVEVEIEVE |
| PI40 | LHESQQEELLELVLRAAELAGVRVRIRFKGDEVHVELEVETAEQCPPALRLLAALLETLRAAGQRVRVELHLGPGVRAADVRRMVEALLAAGAELRVEMHLGTPEAVEALKAVMELAPDAEIRVHIGAHIKQQRQLYRDVREAAKKAGVEVEIEVE |
| PI41 | LHESQQEELLELVLRAAELAGVRVRIRFKGDKMTVEISAKGTKVVFEVTEAAYENLKALLEEILKDKSVTPVEFLTRLIRTMLKCGQKVEVGIGPELVEEILPKLAEVGKETGKKLTCRVRISGYHIKQQRQLYRDVREAAKKAGVEVEIEVE |
| PI42 | LHESQQEELLELVLRAAELAGVRVRIRFKGDEIEVSVERFSEEMQRAMVELAIEKGFSLETLIRVLGISRELVEEVARERFEELRPLLVRVEPGLLVVDSRGVSLEQLRRLCTVAAVLGQKLRVRISGKHIKQQRQLYRDVREAAKKAGVEVEIEVE |
| PI43 | LHESQQEELLELVLRAAELAGVRVRIRFKGDTMTIDLDTPDFGIRLELRGLDDATVRRIGALLEQMVLSDRATAAELGFELYRLCIEAGCPKVSIGVRLSVEQYEELLKAAKEAGIKTVHILLKGHHIKQQRQLYRDVREAAKKAGVEVEIEVE |
| PI44 | LHESQQEELLELVLRAAELAGVRVRIRFKGDEVSLEVDLEEASREQALRGLCALLQACLKAGLEELAEELKEKAAELLEDVEVPEELDCSDVRHPIEALAVGGLIYVLGELGRLKKVRVRIGGHIKQQRQLYRDVREAAKKAGVEVEIEVEGD |

| | |
|---|---|
| PI45 | LHESQQEELLELVLRAAELAGVRVRIRFKGDEMYIEIEAEADQVEGVIAMVREICDGIKELGVKNISLGLSIASAEANTAALGILADGFIESGVDISELELTLSSGAAGKHVKRLLEECKVKKVKVHLRGAHIKQQRQLYRDVREAAKKAGVEVEIEVEGD |
| PI46 | LHESQQEELLELVLRAAELAGVRVRIRFKGDRVTIEVETEEDLVRIVLVVLETDPSGMVLLARALAEQAAAEGEEAVERLRRVCEELAKDPRSAPFLLAMEAFLKILVDKSAKKVRIRIGVMHIKQQRQLYRDVREAAKKAGVEVEIEVEGD |
| PI47 | LHESQQEELLELVLRAAELAGVRVRIRFKGDEVELLDPLSLETLEALLEVAIEEGRSVGTMVEALSFCAEQSGLREELLAWCRERLEPYMEGNELTVEELEEGKRAVVCGLLVARETGEKLRIRIRGLHIKQQRQLYRDVREAAKKAGVEVEIEVEGD |
| PI48 | LHESQQEELLELVLRAAELAGVRVRIRFKGDSVEIEFGFEARLSREEFEAREREYERAREELVEFAKEHKGIKISGSGVLEGVSDEQLPVVRRQLERGFELMAELGKATGCKVKFRFRIHGHIKQQRQLYRDVREAAKKAGVEVEIEVEGD |
| PI49 | LHESQQEELLELVLRAAELAGVRVRIRFKGDEVEIEFGFEPGLSAEEFAAREAGYPAARAELLAFVAKHKGIKISGRGELRGVSEEQLPVVREQLERGGELMAEMGKATGQKVKLEFRIHGHIKQQRQLYRDVREAAKKAGVEVEIEVEGD |
| PI50 | LHESQQEELLELVLRAAELAGVRVRIRFKGDEVEVEAEEGLEALERAAREVFREEGYSVTTGIALARLLARLGSPAVQAELDELNKKEEIELVCTSKDVGEIKWRIALALAAPNVKKLRVRISGVHIKQQRQLYRDVREAAKKAGVEVEIEVEGD |
| PI51 | LHESQQEELLELVLRAAELAGVRVRIRFKGDEVEVEVEEEIEKLYEAAKEVFIEEGYSVETGMALVQLFARLGNEEVLKELEELSKKKEIELKLTSKNVGEARLKIALALAAPNVEKLKVHVSGLHIKQQRQLYRDVREAAKKAGVEVEIEVEGD |
| PI52 | LHESQQEELLELVLRAAELAGVRVRIRFKGDEVEVEAEEALPELYRAAKEVFIEEDYSVTTGIELWRLFARLGVPEVLAELEELNKHEEIELVCRSKNVGEIKWRIALALAAPNVKRLKVRISGLHIKQQRQLYRDVREAAKKAGVEVEIEVEGD |
| PI53 | LHESQQEELLELVLRAAELAGVRVRIRFKGDEVTLETELPAEEVVRRLEALRAEERARGGDYSFVRMCLAAFLELTTGEGEAVYREMAKEEIVLRARTTPESAAYMIEQIGKITGEKLRIRIGAAHIKQQRQLYRDVREAAKKAGVEVEIEVEGD |
| PI54 | LHESQQEELLELVLRAAELAGVRVRIRFKGDEVEIEVEGDPLRLVRALLELVTREGASVGLALTALLQLSKQIPRQELLDLTQPAYEDKTVRTRGEALARILGAYLKGEKKLEIKLTGAHIKQQRQLYRDVREAAKKAGVEVEIEVEGD |
| PI55 | LHESQQEELLELVLRAAELAGVRVRIRFKGDEVEIEVEGSPLALMEAIVELCTERGESVGLAFTALLQIAKDVPREELLALTEPALKDKTVRTRGEALLRILGAYLRGEKKLEVRLSGAHIKQQRQLYRDVREAAKKAGVEVEIEVEGD |
| PI56 | LHESQQEELLELVLRAAELAGVRVRIRFKGDEVEIEVEGDVLRLFEAIVRVCTEEGASVGLALEALLLLSKSMPREELIKICEPALKDKTVRTRGEALLKILGAVLLGEKKLEIRLTGAHIKQQRQLYRDVREAAKKAGVEVEIEVEGD |
| PI57 | LHESQQEELLELVLRAAELAGVRVRIRFKGDSVEIEFEADLSGLSAEEAERMREALAELGEGLRKLGVKVPVTARLNVRGASREVLELVMEGFGALTHCKDLLVRVRLSGHIKQQRQLYRDVREAAKKAGVEVEIEVEGD |
| PI58 | LHESQQEELLELVLRAAELAGVRVRIRFKGDAVELRLEADLSDVTREEAERMREALRELGEGLREMGVRLPLTAELTVRNASREVLELVFEGLGALTHCEGLVARVRLRGHIKQQRQLYRDVREAAKKAGVEVEIEVEGD |
| PI59 | LHESQQEELLELVLRAAELAGVRVRIRFKGDEVEIELEADLSDLSVEEAERMRRALEELGEGLRRMGVRVPVEAHLEIRNASSEVLELVMEGLGALKHCEGLRVHVRLRGHIKQQRQLYRDVREAAKKAGVEVEIEVEGD |
| PI60 | LHESQQEELLELVLRAAELAGVRVRIRFKGDEVTIECSVEDAVRLAEICLEEGAYGALVVLTHLYPEAMRKLPRCAELLALIERAKAGEEVTLTVRTEGDLQAALTLMSLFKGAKFRIRLRGLHIKQQRQLYRDVREAAKKAGVEVEIEVEGD |

| | |
|---|---|
| PI61 | LHESQQEELLELVLRAAELAGVRVRIRFKGDEVTVEASDEDMLRLAEILDEEGAFGGLVFLVHLRPEVLAKLPRAAELLELMERARAGEEVEVVVETRGDLQAVLTLMSLFRGSRFRVRLRGLHIKQQRQLYRDVREAAKKAGVEVEIEVEGD |
| PI62 | LHESQQEELLELVLRAAELAGVRVRIRFKGDSVEVVMTGTGEALLALTEAQLAEAVALLQRARALGGRVTIEIEDLTLAAMPVLEALMAGCPGCHFRVRLNPGHIKQQRQLYRDVREAAKKAGVEVEIEVEGD |
| PI63 | LHESQQEELLELVLRAAELAGVRVRIRFKGDSVEVEITGTGEALLALTEEQIEEFIRLVARARELGGVITFHFEDLTLAAMPKLERILAGCPGCHWRVTLSPGHIKQQRQLYRDVREAAKKAGVEVEIEVEGD |
| PI64 | LHESQQEELLELVLRAAELAGVRVRIRFKGDSVEVEITIDGEALLALTDEQLDECIALLARARALGGRVTITVRDLLPEALPVVERLLAGCPGCRFRFRLSPGHIKQQRQLYRDVREAAKKAGVEVEIEVEGD |
| PI65 | LHESQQEELLELVLRAAELAGVRVRIRFKGDEVEIEAEGDEEKVRRLEAVIKNILNALKGEKVETVSTETTFSGGKEEALDLFLRAQVEAGEGVHLKVRLKAPHIKQQRQLYRDVREAAKKAGVEVEIEVEGD |
| PI66 | LHESQQEELLELVLRAAELAGVRVRIRFKGDEVEVEAEGDEEKVRRLEAAIKNILNAIKGEKVETEETEATYSGGKEEALDLLLRAQLEAGKGVKLRLRLRAPHIKQQRQLYRDVREAAKKAGVEVEIEVEGD |
| PI67 | LHESQQEELLELVLRAAELAGVRVRIRFKGDEVEIEAEGDEEKVRRLKAVIERILAALEGKKVETVETETVYSGNGEEALDLFLRAQVEAGEGVRLRVRLRAPHIKQQRQLYRDVREAAKKAGVEVEIEVEGD |
| PI68 | LHESQQEELLELVLRAAELAGVRVRIRFKGDGVFISIGRVSEEKNERLLEEIRKAFEELGLDIEVFEGGTIVTKSFLKEMGKYYAKVLEEKKGPVRIRLNGFHIKQQRQLYRDVREAAKKAGVEVEIEVEGD |
| PI69 | LHESQQEELLELVLRAAELAGVRVRIRFKGDGVHISVERVSEAKNAALLAAIEEAFAALGLDVVVYPDGTVVTKNFIDEESAFYAKVLEEKKGPVKIRLSGYHIKQQRQLYRDVREAAKKAGVEVEIEVEGD |
| PI70 | LHESQQEELLELVLRAAELAGVRVRIRFKGDGVMIQLERVDEEKNAALLERIREAFERLGLKVEVHPGGTIVTENFIKEMSKYYEEVLKEKKGPVRFRLSGFHIKQQRQLYRDVREAAKKAGVEVEIEVEGD |
| PI71 | LHESQQEELLELVLRAAELAGVRVRIRFKGDEVELDVSPAELERMIRALEPYEEEARTLRALLELRERLARGEEIELDSRELSPETFKLVFETAASLWKGKIRVRLGHHIKQQRQLYRDVREAAKKAGVEVEIEVEGD |
| PI72 | LHESQQEELLELVLRAAELAGVRVRIRFKGDEVEIEVEELTLEQFDRLAAAMTEFDMGLSAEVLVMIVEKNIKLEPKPVELVLEGVPLAEAKARILRALVIAKQAGEKVRIRVTSFHIKQQRQLYRDVREAAKKAGVEVEIEVEGD |
| PI73 | LHESQQEELLELVLRAAELAGVRVRIRFKGDEVEIEVEPLTLEQLERLARAMVEFDMGLAAEVLLMIVERTITLEPEPQELELRGVSLEEAKAEILLALVRARQAGSKVRIRISGFHIKQQRQLYRDVREAAKKAGVEVEIEVEGD |
| PI74 | LHESQQEELLELVLRAAELAGVRVRIRFKGDEVEIEVEPLTLAQFERMARAMTQFDFGLAALVLKMIVEKTLTLEPRPVKLVLEGESVEEAKARILLALVRAAQAGEKVEIHITSFHIKQQRQLYRDVREAAKKAGVEVEIEVEGD |
| PI75 | LHESQQEELLELVLRAAELAGVRVRIRFKGDTVTIEVEAEGRRRELTLSEELPLEEAIEEISRFFREFWRECGYSEEEVELELVSTSEDVEEVLAEIFERMRAAMREGALKFRLRLRGLHIKQQRQLYRDVREAAKKAGVEVEIEVEGD |
| PI76 | LHESQQEELLELVLRAAELAGVRVRIRFKGDTVTIEVEDRGKRAELTLSEDMSLEEAVEKLSEWFRRFWLECGYTEETVELSLHSTEEDVEKVLEAIFERFRKAMEEGALRFRLRISGLHIKQQRQLYRDVREAAKKAGVEVEIEVEGD |
| MC01 | RAAALDAQKTRLAELSEYLPPDSPEMHDFRHGFDILVGQIHDALHLANEGAAPEEIARLEAEARRQEEELDALFAHVKTGEETPLSRRRLTLIEFRVFVERRAAEARTLSAEELEAARAAALDAQKTLVPAVMERTGETDPDSPEMHDFRHGFDILVGQIHDALHLAN |
| MC02 | RAAALDAQKELEEILRAMGYDPDSPEMHDFRHGFDILVGQIHDALHLANESGDEALLAEARAAAEAALGELVLRVFEDVRARLPEIQARVEALKERAAELSVAELEARRAAALDAQKRLLELMERTPVGARRLAELGPDSPEMHDFRHGFDILVGQIHDALHLAN |
| MC03 | RAAALDAQKLVMEALSKTRPPDSPEMHDFRHGFDILVGQIHDALHLANERGDYEAAARQARALLAELFARHPEAAAVVAAAEARRRAADADIDELRRLLDEALALPAAAAVLEALRAAALDAQKSKAAELAKELPPDSPEMHDFRHGFDILVGQIHDALHLAN |

| | |
|---|---|
| MC04 | RAAALDAQKLAEEVLLAKNAKLGLGPDSPEMHDFRHGFDILVGQIHDALHLANRAGTAAEFEAEQAAKYQLVVDKLQLELLQLRLEELLAAAAADPSVEVLRAARAAALDAQKRMAELLVKYTGADPDSPEMHDFRHGFDILVGQIHDALHLAN |
| MC05 | RAAALDAQKVAMEEYLRIGKKDGLTPDSPEMHDFRHGFDILVGQIHDALHLANTKGEDVEAALERLIAQDEVRGEELRLLLELVRELQRLRENEEERRKAVEEGWRRLRELRAARRARLEEERAAALDAQKRGAAQLAAEGVAPDSPEMHDFRHGFDILVGQIHDALHLAN |
| MC06 | RAAALDAQKTLLEELLRHGAAEGLTPDSPEMHDFRHGFDILVGQIHDALHLANTEGADVEEVLERLIAEDAVRREELLLLREAVETLERLREDEEARRRAAAEFWERWRALRAQRRADLEAQRAAALDAQKSGAAALAAQGVAPDSPEMHDFRHGFDILVGQIHDALHLAN |
| MC07 | RAAALDAQKALLEEALRIGEKEGLTPDSPEMHDFRHGFDILVGQIHDALHLANTLGVDVDAALAELIAADEVRREELELLLEAVRALEELRDDEEARRRYAAEFWEEYRRLRRERRERLEAERAAALDAQKAGAAALAAAGVAPDSPEMHDFRHGFDILVGQIHDALHLAN |
| MC08 | RAAALDAQKECMDLLSRVLDPDSPEMHDFRHGFDILVGQIHDALHLANEAGDEELKQEIDRQAEEVARQRLEQARARLREGEAAAPLREQYRARVAEGATAAQFEAERAAALDAQKTEAARLAREDPDSPEMHDFRHGFDILVGQIHDALHLAN |
| MC09 | RAAALDAQKECLALLSRILDPDSPEMHDFRHGFDILVGQIHDALHLANAAGDEAMKAAIEAEAERYAAERLAQARAALAVGEAAAPLREEFRARVAAGATAAEFEAERAAALDAQKSEAARLAVEDPDSPEMHDFRHGFDILVGQIHDALHLAN |
| MC10 | KDALTKMAAAAADAWSATDEIFDDLYKGGVVSKQDGHDFRHGFWILIGQIHDALHKGEDRTEEFRAQMRAHWDRIIELVEENIEKCGVTALKWYVLHVVIRAMRDPSELEKFIERLKKLSKDALTKMAAAAADAWSAGDAELELLGGTDVDRHDFRHGFWILIGQIHDALH |
| MC11 | KDALTKMAAAAADAWSAADLLQEALGLDKELAHDFRHGFWILIGQIHDALHAGDESALPALEERLEALRERLGELAVERVATAIADPEKFARLLLACARLGAAQFARVVEAVAELLPQLDKDALTKMAAAAADAWSAVDTVASLLSAEEQHDFRHGFWILIGQIHDALH |
| MC12 | KDALTKMAAAAADAWSACDYIERALGVDKDKVHDFRHGFWILIGQIHDALHAGDTSLLPELEAQLRAIREELGDIAVERVEVAISDPQKFRDLLLACGRLGAEEFKRIVEKLVELLPKLDKDALTKMAAAAADAWSAVDTLAALLSEEEQHDFRHGFWILIGQIHDALH |
| MC13 | KDALTKMAAAAADAWSACTYLQESLGLDKDLVHDFRHGFWILIGQIHDALHAGDTSALPALQAQLDAIRAELGKLACERISVCISDPEKFRKLLLAGTRLGAAEFQRIVKQLCELVPLLDKDALTKMAAAAADAWSAVDIYADLLSEEEQHDFRHGFWILIGQIHDALH |
| MC14 | KDALTKMAAAAADAWSAEELLEVAREKYGWSKHDFRHGFWILIGQIHDALHLHEQGKEEEVVRLVAECLEFLEELGVDELCPEYRNALAYARDLVDGYLEIRRFAEAARSASKDALTKMAAAAADAWSAEAGVDELCARYPDVDREVARHDFRHGFWILIGQIHDALH |
| MC15 | KDALTKMAAAAADAWSAEELLEVAKRKYGWSKHDFRHGFWILIGQIHDALHLSLRGEEDKFIQLVSECLDFIFELGVDEHCPEYKEELEYAKYLVDRYLEIKRLAEQLKTASKDALTKMAAAAADAWSAEEPVEELCKKFPSVDYETARHDFRHGFWILIGQIHDALH |
| MC16 | KDALTKMAAAAADAWSADELYERARELYGWSKHDFRHGFWILIGQIHDALHLYEEGKLDEVVDLVQRCLDFLRELGVDELVPEYREELAYAQYLVDRYLEIRELAAKLQTASKDALTKMAAAAADAWSAEEPVDELCAKFPSMDRETVAHDFRHGFWILIGQIHDALH |
| CHD01 | LAGIAEGWLVMERLLRAGADVDEALEAFAEVLRRYGHEPLEAVRREFEAVLAVVRDRVSPFELASLAALSAARLIFEERNDPLAEIIQEYLDLLVAITQQGLNPEETVKFIEDYLLKEYHPKKKEILEKITEEDKEVLEKHRELMDEYSEKGDIERLASVAATAMLHVYFSLGYNDPELSSWIFEFRRESIRNVEEGKRFFEKLRENFEEIIAELYVKHLAKKEKPWRIRQLEKLREIVEERKKKGIELTPFEIALEAYYVAVEEYLKE |
| CHD02 | LAGIAEGWLVMERLLRAGADVDEALEAFAEVLRRYGEERVEEMLEEARRLLAEATAAVELSPFELASLAALSAARLIFEERNDPLAEIIRWTLELLLEMLRAGLSPEETLAAIEERAAELLAEARKEMEEFLSEEDKEVLEKHRELMDEYSEKGDIERLASVAATAMLHVYFSLGARKERSSWIFEFRRESIRNATTEEEIEEFRREVEIEMLEIITEQLAKVDPEKARELRLKVLFARSLLKLREIVEERKKKGIELTPFEIALEAYYVAVEEYLKE |
| CHD03 | LAGIAEGWLVMERLLRAGADVDEALEAFAEVLRRYGFERVEEAVEEARRLLAEAQAAAELSPFELASL |

| | |
|---|---|
| | AALSAARLIFEERNDPLAEIIRWTLEQLLEMLRKGLSPEETLKYLEEKAEEELKIAEEKRNEFLSEEDKEV<br>LEKHRELMDEYSEKGDIERLASVAATAMLHVYFSLGGDEEKSSWIFEFRRESIRNATTEEEIKRMRREVD<br>IKLLEILTEELEKEDPELARKLRLLVLRVRSMEKLREIVEERKKKGIELTPFEIALEAYYVAVEEYLKE |
| CHD04 | LAGIAEGWLVMERLLRAGADVDEALEAFAEVLRRYGHEHLIEPVRERVREIVAKLRDKLSPFELASLAA<br>LSAARLIFEERNDPLAEIIEFTLETLVEILEKGLNEEESVKFIEEKIVEKNAPLLEEAERLFERLLEKLLSKE<br>DKEVLEKHRELMDEYSEKGDIERLASVAATAMLHVYFSLGLNDPKISSWIFEFRRESIRNLEIIPTEELRK<br>LYIRVKMMYELVKILLKVASEEEKKELEKLLELLEKILKSLELILKLREIVEERKKKGIELTPFEIALEAYY<br>VAVEEYLKE |
| CHD05 | LAGIAEGWLVMERLLRAGADVDEALEAFAEVLRREGHEHLIERIREEFLALLERLREKLSPFELASLAAL<br>SAARLIFEERNDPLAEIIKFTLETLERILEAGLNEEESIIEIENRVIEKNKPLLDEAREKLEEIKEKLLSEEDK<br>EVLEKHRELMDEYSEKGDIERLASVAATAMLHVYFSLGLNDPKESSWIFEFRRESIRNLSIFPTPEIRKLYI<br>EVELEYFLVKMLLKVASEEDKKELEELLELLEKVKKALDQILKLREIVEERKKKGIELTPFEIALEAYYVA<br>VEEYLKE |
| CHD06 | LAGIAEGWLVMERLLRAGADVDEALEAFAEVLRREGHEHIIEPMREEFERIYAVLRDKLSPFELASLAAL<br>SAARLIFEERNDPLAEIIRFTLETLVEILTAGLNEEEAIIFIERKFVEKMKPVIEEAERLYEELLEELLSEEDK<br>EVLEKHRELMDEYSEKGDIERLASVAATAMLHVYFSLGLNDPEISSWIFEFRRESIRNLSIMPTRELRFLYI<br>RTHTRLALVQILLRVADEETKKKLEKLLSLLEEIHRTLEQILKLREIVEERKKKGIELTPFEIALEAYYVAVE<br>EYLKE |
| CHD07 | LAGIAEGWLVMERLLRAGADVDEALEAFAEVLRREGVSEEALERWRAEARALLARFAPAPELSPFELAS<br>LAALSAARLIFEERNDPLAEIIRLYLELVGILLELKELDELKLIEEALKRLGKRVLEELRELREETESSLSEE<br>DKEVLEKHRELMDEYSEKGDIERLASVAATAMLHVYFSLGLDEPKYSSWIFEFRRESIRNPEFLDFTLEF<br>LELLFQLDKESLKRATRLVYALLLLIKSLPKLREIVEERKKKGIELTPFEIALEAYYVAVEEYLKE |
| CHD08 | LAGIAEGWLVMERLLRAGADVDEALEAFAEVLRRYGFTGFVERAREVLARHRDRLSPFELASLAALSA<br>ARLIFEERNDPLAEIIDAFLEAVEEIEKLEDLEEVVAYVAVRLSVAIDPSFAAHADEILDILLNIEETVSEEDK<br>EVLEKHRELMDEYSEKGDIERLASVAATAMLHVYFSLGLPRERSSWIFEFRRESIRNYSIEEIKERTEREL<br>RGLMTLGALLRLFSEHRESAIRAARLYVKLREIVEERKKKGIELTPFEIALEAYYVAVEEYLKE |
| CHD09 | LAGIAEGWLVMERLLRAGADVDEALEAFAEVLRRYGFEGAVEAMRPLVEELRERLSPFELASLAALSA<br>ARLIFEERNDPLAEIIRNYLEFFRETLELSPEEVLKKLEELAEEKEKKLEELVMKVLSEEDKEVLEKHREL<br>MDEYSEKGDIERLASVAATAMLHVYFSLGYNDPKVSSWIFEFRRESIRNWADLSPEELKKEIRRKMTVT<br>MAEILVETVLKEKDPELTKEIEENIKFLRNYEKLREIVEERKKKGIELTPFEIALEAYYVAVEEYLKE |
| CHD10 | LAGIAEGWLVMERLLRAGADVDEALEAFAEVLRREGHEHIVERTRELVLPLVGGLSPFELASLAALSAA<br>RLIFEERNDPLAEIIAAFIETVGRLPEEPEEVIEVFNKELAKRITEYLEKVAKEMEELATVELPPEDKEVLE<br>KHRELMDEYSEKGDIERLASVAATAMLHVYFSLGRYDPELSSWIFEFRRESIRNLPSLEDLKKGLRYYKE<br>IVWYTIRNLYGSEELREKTLKKLRDVLTLLKAISEIEKKLREIVEERKKKGIELTPFEIALEAYYVAVEEYL<br>KE |
| CHD11 | LAGIAEGWLVMERLLRAGADVDEALEAFAEVLRREGHEHLVEAIRERVLPLVGGLSPFELASLAALSAA<br>RLIFEERNDPLAEIIAAFLETVGRLKEEPEKVIEVFIEILAERITDYLTRVKEEMEKLATVELPKEDKEVLE<br>KHRELMDEYSEKGDIERLASVAATAMLHVYFSLGRFEPELSSWIFEFRRESIRNLAKLKNLKEGLKNYRE<br>IVWYTVLSLYGSDELREKTLNKLKSVLTLLKAMSEIEKKLREIVEERKKKGIELTPFEIALEAYYVAVEEY<br>LKE |
| CHD12 | LAGIAEGWLVMERLLRAGADVDEALEAFAEVLRREGLLDVERALARIREVLEEVRAVAGEELSPFELAS<br>LAALSAARLIFEERNDPLAEIIEISLEFLKRIKVESLEKVLEEIEEEYIPLLEEKLKEVEKAVLEKLSKEDKE<br>VLEKHRELMDEYSEKGDIERLASVAATAMLHVYFSLGKFSPEESSWIFEFRRESIRNLGVDPKIAKLRIQI<br>ELLKKINEILGGDEKLMKEAVMYYSIIMMKYKLREIVEERKKKGIELTPFEIALEAYYVAVEEYLKE |
| CHD13 | LAGIAEGWLVMERLLRAGADVDEALEAFAEVLRREGVEVDLAPIRELVERFRRLGLSPFELASLAALSA<br>ARLIFEERNDPLAEIIRAFEELAERMLELGGMKGLEAWLEERGEKADELFESLLSEEDKEVLEKHRELM<br>DEYSEKGDIERLASVAATAMLHVYFSLGVNDPELSSWIFEFRRESIRNAEEDVLKDIRLRESAFRAELLVK<br>LKEEELPEELVNELRVYVLLVKNLPKLREIVEERKKKGIELTPFEIALEAYYVAVEEYLKE |
| CHD14 | LAGIAEGWLVMERLLRAGADVDEALEAFAEVLRREGVEVPLEPIRELVERFRERGYSPFELASLAALSA<br>ARLIFEERNDPLAEIIAAFIELAERMLELGGMEGLYEWLREELAEARELFLSLLSEEDKEVLEKHRELMD |

| | |
|---|---|
| | EYSEKGDIERLASVAATAMLHVYFSLGLNDPELSSWIFEFRRESIRNAEEDPLEEIELREGAIMAQMLVLKYSEELSKELVKRLRVTVLLVKNLPKLREIVEERKKKGIELTPFEIALEAYYVAVEEYLKE |
| CHD15 | LAGIAEGWLVMERLLRAGADVDEALEAFAEVLRRHEPDLPEEALRRMLEEARALLEEFRKRGVSPFELASLAALSAARLIFEERNDPLAEIIDEFLALSREEDVLKALEERAEKAKEEAKKLWEELAKYLPAEDKEVLEKHRELMDEYSEKGDIERLASVAATAMLHVYFSLDPENKELSSWIFEFRRESIRNWHEIPMEELVELMWLELQLLVAKLYYKYKVPPEEKEKFKELYEKFKKAASQLLGYYKLREIVEERKKKGIELTPFEIALEAYYVAVEEYLKE |
| CHD16 | LAGIAEGWLVMERLLRAGADVDEALEAFAEVLRREGHEAVEAAREVVERHRDRLSPFELASLAALSAARLIFEERNDPLAEIIDIYLEVVDELDLSNEEIIELFIRKLKEFLEKRREKALEKKGAMSEEDKEVLEKHRELMDEYSEKGDIERLASVAATAMLHVYFSLGKFDPEKSSWIFEFRRESIRDEEIFEENKKVFEFFYEMLLEVSFEDAEELTKLYRNILEMEKKLREIVEERKKKGIELTPFEIALEAYYVAVEEYLKE |
| LBM01 | KYMLVGRMEEELNVTIFRWVDEDESEEDIEKKLLCEMQLKATIKNPELTRNKFPVLKFERKGNAIYATASLISKEEFEKIYSKETVEEVNIAKEVATLRQAGDDFSRRYRRDFAEMSEFMKDIEPLPEEVVEETLREIARIRLEHGDDLEGIKEVALEVFTHTEKMKTVVEELFRDGVNWGRIVAFFEFG |
| LBM02 | KYMLVMRLEGANVTIFRWAEEEKLEEVLEQSRKQLAFKALVEVAPQPEVLARFYRDISEELIALLRSFVALYGGEVPAELEALSPDPVARLLAVVQAALPTLRQAGDDFSRRYRRDFAEMSPLVKKVAENEDLDEQLEYLSTVVEELFRDGVNWGRIVAFFEFGRQYYEALGITNELGSFLILLGVIVTGDISTLEEMIEELEELLEYEEERKGNAIYVTARFK |
| LBM03 | KYMLVIEIEHPELENVTIFRWFEKDTWAEEILDFVEEVRAALDGKEYKKVKLSEEEAERKTLTLLGALFVALYGGEDLDELLKMVEDEEKWEEVMEIFLETLRQAGDDFSRRYRRDFAEMSKLLKEWKEKHKSEEELLDELLTVVEELFRDGVNWGRIVAFFEFGRLLGYSLEESLELFLRLLRRGSSMLVGGLAGMAVADLLRLRYGLEVEQEVKGNALHIRAVG |
| LBM04 | KYMLVIEDPANPNVTIFRWVEEKDLEREIYRTLKELYPEELEKEVYTRLSELLAEFVRLYGGRELPVVSTREELFELFREAEPTLRQAGDDFSRRYRRDFAEMSKFLAEYKAKYAGDEEAIEEMVETVVEELFRDGVNWGRIVAFFEFGRLLGLSDEEILELLLEYLGKELVREYFKKHGELKGNAVRLR |
| LBM05 | KYMLVARVGPNVTIFRWLDTIEEAIEYLKETFKEEIDEFKRRLLKVLMIEFVSLYGPEKREQVEKEVESLSLEELLAIPEIFRETLRQAGDDFSRRYRRDFAEMSRTFRKQLEELSRVLGEEEARRLLVDTVVEELFRDGVNWGRIVAFFEFGRIAGLSEEEVMELLADYLLLKDTELNKKAGYIFEIKGNAIIFSLAE |
| LBM06 | KYMLVIRLEGLNVTIFRWAEKEEVEKLIEETLKEVEEELERDKKKLKEMLGEELVDKIINWVKENLDEFVLLYGEEGEKLLSELNELEKLSLFLETLRQAGDDFSRRYRRDFAEMSKVLTKLKKEKTPEEVAEALRTVVEELFRDEVNWGRIVAFFEFGRLTGLGPLLENALILEFYKPLKEKLVTEVEVKGNAVHVTVKVK |
| LBM07 | KYMLVIKIKTKGRNVTIFRWATEEELKKFLSLAKELLERRIEEDKRRLREVFEETGFSIEEFVTLYGKPEHYELAEASGEEDKLMTLALLTLRQAGDDFSRRYRRDFAEMSKMIKSLVEKLKDKSIEEVKEELKDFMDTVVEELFRDEVNWGRIVAFFEFGRLVQTELTKLELEKEKGLSEEEVDFLLTLSEMIELGAKVEKKGNAIYITYEE |
| LBM08 | KYMLVIRSHGLTNVTIFRWVEEEEVEEVIEKLLRVLKFVEELYYEILKLLEEKVSGRVLELLKEFVALYGGEDPAALEGSLEEALAHAAPTLRQAGDDFSRRYRRDFAEMSAMIDDYLTENTNATLEDILEFVKTVVEELFRDGVNWGRIVAFFEFGRLVVEAVLRTLYRNIDEVEIEETNMIAKVKIERKGNAVHVDVELTPG |
| LBM09 | KYMLVVTDRRANVTIFRWAEEGHVLLGLYEALKEAIGRSLGVTDEETEQEVNKLQIEVLGEFVRLYGPELAPAVEELRKNITDAEKALEEVRALVGPTLRQAGDDFSRRYRRDFAEMSRMISEEIKKHADNIEELLEFFGTVVEELFRDGVNWGRIVAFFEFGRILLEEAEKEPRISQESLEKLQEFIEEVLTGAMLLLYFQKQGWRIERKGNAIYVTA |
| LBM10 | KYMLVYRSQKDKNVTIFRWIEKEDLFEEVYRILREEYGEELERQVNLRVSQLLQDFVELYGGEKLPLVASREGVRELFARARPTLRQAGDDFSRRYRRDFAEMSRLIAEEKAKYAEDAEEQERFFRTVVEELFRDGVNWGRIVAFFEFGRLWGLSDEEIFEALLEFIGRQLVKEHLKKNGEIKGNALYLE |
| LBM11 | KYMLVIKIKTRDENVTIFRWLSEEEFEKFLKTTEELYELILKWEVETLRALFAKAGVSAADFVALYGRPEDEALAERLPLEDREEALLLVTLRQAGDDFSRRYRRDFAEMSKIIKELVEKAKDLTIEEQLEYFKPYFDTVVEELFRDGVNWGRIVAFFEFGRLAFREILKQYLSREKGFSLELAKLAVTLAELRHHIVSIERKGNAVYITV |

| | |
|---|---|
| | EE |
| LBM12 | KYMLVFRDEERNVTIFRWVEGESIILAIAKALMEAEQRALGISEEEAERYRKVQQAAAMVDFVSLYGPELAPKAKELLEKVLEDEEAYEELKELVAPTLRQAGDDFSRRYRRDFAEMSEMIREKVLENSGNVEKLIEFYSTVVEELFRDGVNWGRIVAFFEFGRILLRTVEELGVLSEEELEKLKKYIEEEFDAALYVYYQLKQGLKIERKGNAIYITR |
| LBM13 | KYMLVVTDPRSNVTIFRWAEEGYVVYELGEAIVRARARSMGWADEEVPLVKNILVIKALLSFVKLYGPELTPELEKVIEEEKDVEVAKKKVLELAEPTLRQAGDDFSRRYRRDFAEMSRMIRELLRKYTGDVEKIQEFFGTVVEELFRDGVNWGRIVAFFEFGRLLLKEAEENEQMSEEDIEELKKFIEDMFDNVLVLLQAQEEGWRIERKGNAIYVTR |
| LBM14 | KYMLVVRDEKANVTIFRWVEEGKEVIGAAEALIELKGKALGLTPEEKERLLDLYVGEALVDFVALYGPEKAPEAAALLEEAKKDPAYFAELLKLAEPTLRQAGDDFSRRYRRDFAEMSRQLKALIEEHADDLEKLIEYFNTVVEELFRDGVNWGRIVAFFEFGRIMLELVEENKKVSEENKEKFKKYIEETYKLILALYYFQKKGLKIEKKGNAIYLTF |
| LBM15 | KYMLVVTDPEANVTIFRWAEEGDFLLALGDALVEMEAAKAGADAEGGEALRRERITEALISFVRLYGPELAPALEAARAETADAAEALARIRALAEPTLRQAGDDFSRRYRRDFAEMSRMIKDLLRKHANDVEAIKERIGTVVEELFRDGVNWGRIVAFFEFGRLVLKAAEEEPAMSEEVREELREFLERLFESVLLLYRAQEEGYRFERKGNAIYVTK |
| LBM16 | KYMLVIYDEKNPNVTIFRWVEKEKAEEELINTVYELYKEEIEKETILRLSRLLKDFVRLYGGEEIELVESREELFELLKRVEPTLRQAGDDFSRRYRRDFAEMSRFIAAFKEEHAGDEEAIKDFIDTVVEELFRDGVNWGRIVAFFEFGRLMGLSTDERYELLFNTMKKILIKEYIKKHGTKKGNAYHLK |
| LBM01_C4 | KYMLVVTDTNRNVTIFRWLDELPDEETLKLEAIKLLFYAMGPREALAFVLALMLWDKLDDFVSLYGPDVVPQWEATKSLSERLHLAIATLRQAGDDFSRRYRRDFAEMSKVIEREVKKSDLESLLDFIETVVEELFRDEVNWGRIVAFFEFGVIVLEHASEHSLELIDKMVEFISRLFPGSEEKMKQLILELVDIEIERKGNAVRITITFP |
| LBM02_C4 | KYMLVVEDEKANVTIFRWLEELPGEEELKIEAILLLLYEQGVRAALSFLLTTALWERVDDFVSLYGADRVAEWEATTSEAERLRLAMETLRQAGDDFSRRYRRDFAEMSKIIEEEVKKSDLEELLEFIGTVVEELFRDEVNWGRIVAFFEFGVIVLEFAKEHALELIKKFLEFVSSLFPGSLEKMKKRILELVSIEIERKGNAVYIRLTFP |
| LBM03_C4 | KYMLVVRDEKRNVTIFRWLEELPDERTLRMEAIKLLFYAEGVRTLLSFLTILMLWEELDSFVTLYGPERAAEWEATRSEAERVWLAMETLRQAGDDFSRRYRRDFAEMSKIIEEEVKKSNLKELLEFIRTVVEELFRDEVNWGRIVAFFEFGVIVLEHATEYALELIDEMLEILSSLFPGSRERMFERAERLFDIEIERKGNAVDITITRP |
| LBM04_C4 | KYMLVIEARNPLGNVTIFRWVDDIEKYAKSMKDVTLETSSKEKQEEVQDEFVKLYGGEKYEEWKKITDPEERQKLFNETLRQAGDDFSRRYRRDFAEMSRALREEFEKLEGDAKAVIEMAVTVVEELFRDGVNWGRIVAFFEFGGIIFGSMAAALYEDNDAIIEGLEAVLKTLEKKGLKLHIVIKLTTEVKGNAMRITIEFRVE |
| LBM05_C4 | KYMLVIEARNPLGNVTIFRWIDDIEKIKKSAKNVVLETSKKEETEEVYKEFVKLYGGEKYEEWLKIKDKEKKLELFQATLRQAGDDFSRRYRRDFAEMSRSLRKHFEELKGDKEAVIEAMVTVVEELFRDGVNWGRIVAFFEFGGIIMGYAAYALYEDVDAIVEALEAVLKTLEEKGLKLHIRIRLEVEVKGNAARIRIEYEVL |
| P401 | MEFVEEFKKLSRTERMMLLAWAMKNGTIGDVVVALVTGENAVVIVEIYAVAQSSHVPPKITLEKILNVHVGQAETLADVPEVLRELVETVLKPAASTNYGVNLGAFEGDPAHAQRLADLVLEALGLPPQPVQFLTYSNNVISGDFLWINTYTGEVVRYPMTITVPDTPEAQGIGNLLSKLAQLAKEDPGLVVLLVARGDYVNWYFTVFVIAARLG |
| P402 | SRTEAMMLLAWLVATGRRLGLSEAEIEEARSRLTLALVRKGLKEGAVVVVRLSARGSSHVPPSVSYTHLLDARIVEAEDKLEEVKKIIKEILEEHLKLTNYGVNLSVFTFNEEAKELGEKMKEILEEVLKEFGVEAKLEVGLVEGNTVTTNMVWINTYTGEAMELGPTTFTFVSDAVIEFAELLKEAGSVGIAISRGDYVNWYFDTDLVILPKK |
| P403 | SRTEAMMLLAWLERTGREQGTSEEEMLEVRSRLTLALLKKALEEGAVAVITMEARAQSSHVPPDVSFTHLLDTRIVEAEDVFEETKRLLLEVLNKLLSDTNYGVNLSVYIYTEEAKELGEKMREILEEVLRELGVE |

| | |
|---|---|
| | AELEVGLVEGNTVTVEMVWINNYTGELMKLGPMTFTFESELVKEAAELLKEAGSVGIMIARGDYVN<br>WYFRVEVIVVPKK |
| P404 | SRTELMMLLAWYTREINKDASKASEIAKELIELAKREGKLILVGAEVEAQSSHVPPTYYITKITSIWLLS<br>EEEALKALTEFLKSVETNYGVNIALWGPAEGAAELAEIVSKAVKEAFGVEIEMEIEDTEISGNKVVATH<br>AWINTYTGEVLHLPPAEIVFTDNGAGKFLYELAKELPGAMVGRGDYVNWYFTSIWLFR |
| P405 | LEKQLELAEELRKLSRTERMMLLAWYLQKGDKEMADLIRSVLFQEELEKNPEAWVLIKFKVKAQSSH<br>VPPQYSLSEPAEFEIIKTDEEVEELLKKWFESIPTNYGVNLIALDKDDVEKLFEENIDLPANKIVQALLD<br>YINSEPHLQYWINTYTGELVRLEIRNGQLVSQEVSELPFGSTQKERAEMIYSLVKKIEQETGKAVIISRG<br>DYVNWYFESYLFVK |
| P406 | MEEELALAEELEKLSRTELMMLLAWYLQAGDTAMANLIYSVLLREEIKKHPDSYLVIEFRAKAQSSHV<br>PPSYSLSESATASLIEKEEEVEEYLKEWTKSIKTNYGVNLILVDKEEVEKLFDKNLNLPLDSLVEELLKY<br>INEKEHTQYWINTYTGEVVTQKIKNGELVERKEEQLPFLSTQKERAEAIYELIKKLEKETGKAVIVMRG<br>DYVNWYFHGLLIVK |
| P407 | MLERIREVEEFARRAATLSRTELMMLLAWAIREGDKVRADILYLRAAGIGDEIAFIFRGKGQSSHVPPS<br>VTFTDELAVADSFEDLVTLAETTERVDELAEDEEKILEVLKETAKKLTNYGVNLIRIRYEVSEETLKKIE<br>AGDLESLPTLTMEADATWINSYTGEIERFKMTVTFSGIIFGLMLRRALARGERKTILLVARGDYVNWYF<br>TVVILTP |
| PO1 | DALAERLRELAAAAAARGDLRATAMALLALSRVRDGDRVEAFRTLGRLAVELARLGWDEETAALLE<br>EIERLLAELEATADPETVAELRLEFAFTRLAIAAARAGGEVFRVTSEEELDEAIAKIAEVGKHYMVLVSV<br>GKGTLILVTGVREVAELARELFPDAQLEFLTENLYFQARRASLALLEAA |
| PO2 | DALAELLRELAAAAAARGDLRATAMALLALSRVRDGDRVEAFRTLGRLAVELARLGWDAEVERLRA<br>RLAELLAELLAEVDADTAEELRLAWAFTELAIAAARAGGEVFAVTSEEELDEAIAKIAEVGKHYMVLV<br>SVGKGTLILATGDRRVADLARRLFPDAQLEFLTENLYFQARRASLALLEAA |
| PO3 | SLEELLRELAAAAAARGDLRATAMALLALSRVRDGDEKAAMADLLRLIELLRELAAERPEVREELEPR<br>IAELERLVEPYKVEYREELRRALEEALDEALEAVEAGEKVVIEKELVTTRIFIANVTSEEELDEAIAKIAE<br>VGKHYMVLVSVGKGTLILIASDDEEILREVVEERFPDAQLEFLTENLYFQARRASLALLEAA |
| PO4 | LEEVLRELAAAAAARGDLRATAMALLALSRVRDGGDLKEALLLLARLVEYTIREGGPADVRLLLEAL<br>RRMTELGLPPEVASFLRQLAALLEFVLEERDNIELIFMVTSEEELDEAIAKIAEVGKHYMVLVSVGKGT<br>LILIATKDLEATMEKLRKLLPDAQLEFLTENLYFQARRASLALLEAA |
| PO5 | LAEVLRELAAAAAARGDLRATAMALLALSRVRDGGPLEDALLLLARLVEHVAREGDAEALRLVREAL<br>ERAVELGLPEEAAAFLRQLAALLAFILEQRDNIELLLLVTSEEELDEAIAKIAEVGKHYMVLVSVGKGT<br>LILIATKDPEATMAALRERLPDAQLEFLTENLYFQARRASLALLEAA |
| PO6 | RLRERLIALLRELAAAAAARGDLRATAMALLALSRVRDGKENLTEATMLLIRTLAELAALDPEALAAL<br>AAELRRLIEELREGDEDYLEMLENMLLAAELLLLLPDEVVEVVTSEEELDEAIAKIAEVGKHYMVLVS<br>VGKGTLILAAGPLDRIERARERVRELRPDAQLEFLTENLYFQARRASLALLEAA |
| PO7 | ELRERLIELLRELAAAAAARGDLRATAMALLALSRVRDGKNDLTEATMLAMRTLARLAALDPEYLRR<br>LREELARLIEELRDGDREYLEMLENLLIVAELLLLPEEVVEVVTSEEELDEAIAKIAEVGKHYMVLVS<br>VGKGTLILAAGPLERIERAREAIQALRPDAQLEFLTENLYFQARRASLALLEAA |
| PO8 | MREELLALLRELAAAAAARGDLRATAMALLALSRVRDPATPIEEVAAAVERLLAALADADAELLARL<br>RRTADLLTLRERTRRTLRELAPAAPEEKIEELVEKIVRLFEEGKAVVFFVTSEEELDEAIAKIAEVGKHY<br>MVLVSVGKGTLILIAGDEETVRELAEELLELFPDAQLEFLTENLYFQARRASLALLEAA |
| PO9 | SMRAALEAELRELAAAAAARGDLRATAMALLALSRVRDGEEGVAPLLLRMYEEALRRGDEELAARL<br>RALLLRLTADDEAARLRLRLAEAEARARRAGIEVIASRDPAVAPIVIELLERFRERLPNSVIALVTSEEEL<br>DEAIAKIAEVGKHYMVLVSVGKGTLILIAGEREDLEEAREIILELYPDAQLEFLTENLYFQARRASLALL<br>EAA |
| PO10 | AVEVVEARLRELAAAAAARGDLRATAMALLALSRVRDGGEVPGGLELIRRLERELGLYSEEGRLERM<br>KEMAEEMIEAGPSDELRRYLEREKARMEIKLELLELAKEWEKKGLVASVVTSEEELDEAIAKIAEVGK |

| | |
|---|---|
| | HYMVLVSVGKGTLILVAGEPEQVREAAEEILSLYPDAQLEFLTENLYFQARRASLALLEAA |
| PO11 | AVDFVAAELRELAAAAAARGDLRATAMALLALSRVRDGETVVEGLDFIERLARELGLYSREGLLERLL<br>ELARRMIEAGPSPELRAALENRAGQYEIELELLEKAKEWEKKGVVASVVTSEEELDEAIAKIAEVGKH<br>YMVLVSVGKGTLILVAGEKDQVEEVAEEIKKKYPDAQLEFLTENLYFQARRASLALLEAA |
| PO12 | SPLLELADLLRELAAAAAARGDLRATAMALLALSRVRDLSLDEEKLREALEWARRVVARLDPALEPRV<br>ARLEEEALEHAPKVREVAEELGWTDEGIGTLAERLIEEALRAARTDQARFDEIAEKLIRALGVGYIARV<br>TSEEELDEAIAKIAEVGKHYMVLVSVGKGTLILIGAPLPVARLFARVAERLDAQLEFLTENLYFQARRAS<br>LALLEAA |
| PO13 | APLLELGALLRELAAAAAARGDLRATAMALLALSRVRDTPLDPAELARTLAWVGAVVARLDPALRPR<br>VEALRAEALARAPTVRRVAEEMGWTDMGIGTLAEKLISELLEAARTDRERFEEIAEVLKRAMGVGYI<br>AAVTSEEELDEAIAKIAEVGKHYMVLVSVGKGTLILIGAPEAVARLFAEQARKLDAQLEFLTENLYFQA<br>RRASLALLEAA |
| PO14 | DLVVARIRELAAAAAARGDLRATAMALLALSRVRDGASLEEALELLVRLLAEREEERFERLLALALEL<br>FERSRELAERFLRIVERVLGPEVAELLRFALEAEELFREETDPERLIEEIERLARRVGFALELTVTSEEELD<br>EAIAKIAEVGKHYMVLVSVGKGTLILYTFEHEVVGRAMYRALSLRGDAQLEFLTENLYFQARRASLAL<br>LEAA |
| PO15 | SLLVERIRELAAAAAARGDLRATAMALLALSRVRDGASEEEVLELLVRLLAELEDLRFERLEELALELY<br>EESRELAERFLRIVREILGPEVARLLEFVLRVKEIFEKEKDKEKIIEKVFELAREVGHALTISVTSEEELDE<br>AIAKIAEVGKHYMVLVSVGKGTLILYAFEHEAVGRAMYDALALRGDAQLEFLTENLYFQARRASLAL<br>LEAA |
| PO16 | SEVEERLRELAAAAAARGDLRATAMALLALSRVRDGNIEEANRLLLRLAQYVAARDPALLERVLALA<br>RLLAEQYPEAEGTLLMIEAIRSAARAGNIAFLVPVSKEVAEKILKLVELLNKEIPGEAVIAVVTSEEELDE<br>AIAKIAEVGKHYMVLVSVGKGTLILIATDHLDVAKRIAEEVLPDAQLEFLTENLYFQARRASLALLEAA |
| PO17 | MREEVEALLRELAAAAAARGDLRATAMALLALSRVRDGGNLASQVVALARLARVLSEADHPRVRRV<br>LELLAAEEHPEAEELVKGALVAIEKGTEIYNGTSPATEIALKIVLATKSVSLVYIVTSEEELDEAIAKIAEVG<br>KHYMVLVSVGKGTLILIAVEDESVVPAIRELLPDAQLEFLTENLYFQARRASLALLEAA |
| PO18 | LAEYLRELAAAAAARGDLRATAMALLALSRVRDGELLALAETLVALYLLYADRDADVAARLREHALA<br>AIERVLAEEKSEVARARALVLRLNLERAAEGTVVELVTSEEELDEAIAKIAEVGKHYMVLVSVGKGTLI<br>LVAGDRETVEAVAAELRAAFPDAQLEFLTENLYFQARRASLALLEAA |
| PO19 | MADFLRELAAAAAARGDLRATAMALLALSRVRDGLLLEEAAETLVALAELYADKDAAVAARLEALAR<br>ELLREVLEREKSEVTRARALALELRLRTAAAGTVVEVVTSEEELDEAIAKIAEVGKHYMVLVSVGKGT<br>LILVAGDEETVRRAAAELREAFPDAQLEFLTENLYFQARRASLALLEAA |
| PO20 | MAEFLRELAAAAAARGDLRATAMALLALSRVRDGLKLEEAARTLLALAELYADRDPAVAARLREHLLE<br>LLEEVLKEDKSVVTQAEALALRLRLELAERGTVVEVVTSEEELDEAIAKIAEVGKHYMVLVSVGKGT<br>LILVAGDLADVERVAEELRAAFPDAQLEFLTENLYFQARRASLALLEAA |
| PO21 | IADLLAAFLRELAAAAAARGDLRATAMALLALSRVRDGDLEEALRAVAEAILRLPPELVGIGAAAARR<br>VGGEYRDEVPEVLEAIRALRDELPDRGVVAVLNLVMASINRGVIPEEKIEEVIEIAKKIGAKVIEVVTSE<br>EELDEAIAKIAEVGKHYMVLVSVGKGTLILIAMDLDGAIEFALELYKYDAQLEFLTENLYFQARRASLA<br>LLEAA |
| PO22 | SALLELGALLRELAAAAAARGDLRATAMALLALSRVRDIPLDPAAAEEVLEWVRRVVARLDPALEPRV<br>RRLAEEALAALPRVRAVAERLGWSDMGVGTLARRLIRELLAAARTDDARFEEIARVLREALGVGYIAR<br>VTSEEELDEAIAKIAEVGKHYMVLVSVGKGTLILIGAPAAVARLFEEVARQEDAQLEFLTENLYFQARR<br>ASLALLEAA |
| PO23 | SEVEARLRELAAAAAARGDLRATAMALLALSRVRDGELEEAARLILRLAQYVASRDPALLERVLELAR<br>LLAEQHPEAEGLYLMIEAILLASRAGNTASLVLVSTEVAKKVLELVKKLNEKIPGKAFIAVVTSEEELDE<br>AIAKIAEVGKHYMVLVSVGKGTLILIATDDIETAAKIAQEVLPDAQLEFLTENLYFQARRASLALLEAA |
| PO24 | MLEEVEAALRELAAAAAARGDLRATAMALLALSRVRDGGDLAAQVAAIGELARVSGEEERPRLRAIL |

| | |
|---|---|
| | ERLAEAHPEARELVEGVLVAIENGTEIFNTTSPATELALRIVRATRSVRLVAIVTSEEELDEAIAKIAEVGKHYMVLVSVGKGTLILIAVDDAAVEPAIRALLPDAQLEFLTENLYFQARRASLALLEAA |
| PO25 | MEEFLRELAAAAAARGDLRATAMALLALSRVRDGLLLELAETLVALARLYAERDPEVRRRLEELAREAIRKVLAEEKSEVTRARALALELELELAERGTVVEVVTSEEELDEAIAKIAEVGKHYMVLVSVGKGTLILVAGSPEDVAAVAERLAAAFPDAQLEFLTENLYFQARRASLALLEAA |
| PO26 | MAETLRELAAAAAARGDLRATAMALLALSRVRDGDLLALAETLLAMYLHYRDVDAEVAALLRRWLEEALERVLREEESEVARARALALRLQLRLADKGVLVEVVTSEEELDEAIAKIAEVGKHYMVLVSVGKGTLILVAGDEADVAAVAEELRREFPDAQLEFLTENLYFQARRASLALLEAA |
| PO27 | LAEFLRELAAAAAARGDLRATAMALLALSRVRDGLLLELAEALLALYRLYADVDAEVAARLRAHALRAIDEVLARDRSEVTRARALVLRLNLEGAEEGTLVELVTSEEELDEAIAKIAEVGKHYMVLVSVGKGTLILVAGSPEAVRRAAEELEREFPDAQLEFLTENLYFQARRASLALLEAA |

**Table S5. Mutant variants of PO-responsive oligomers analysed in this study.**

| Design name | Amino acid sequence |
|---|---|
| PO18 L18A (PO18s) | LAEYLRELAAAAAARGDARATAMALLALSRVRDGELLALAETLVALYLLYADRDADVAARL REHALAAIERVLAEEKSEVARARALVLRLNLERAAEGTVVELVTSEEELDEAIAKIAEVGKHY MVLVSVGKGTLILVAGDRETVEAVAAELRAAFPDAQLEFLTENLYFQARRASLALLEAA |
| PO5 L18A (PO5s) | LAEVLRELAAAAAARGDARATAMALLALSRVRDGGPLEDALLLLARLVEHVAREGDAEALR LVREALERAVELGLPEEAAAFLRQLAALLAFILEQRDNIELLLLVTSEEELDEAIAKIAEVGKH YMVLVSVGKGTLILIATKDPEATMAALRERLPDAQLEFLTENLYFQARRASLALLEAA |
| PO15 L20A | LLVERIRELAAAAAARGDARATAMALLALSRVRDGASEEEVLELLVRLLAELEDLRFERLEE LALELYEESRELAERFLRIVREILGPEVARLLEFVLRVKEIFEKEKDKEKIIEKVFELAREVGHA LTISVTSEEELDEAIAKIAEVGKHYMVLVSVGKGTLILYAFEHEAVGRAMYDALALRGDAQL EFLTENLYFQARRASLALLEAA |
| PO5 S176A | LAEVLRELAAAAAARGDLRATAMALLALSRVRDGGPLEDALLLLARLVEHVAREGDAEALR LVREALERAVELGLPEEAAAFLRQLAALLAFILEQRDNIELLLLVTSEEELDEAIAKIAEVGKH YMVLVSVGKGTLILIATKDPEATMAALRERLPDAQLEFLTENLYFQARRAALALLEAA |
| PO5 S176D | LAEVLRELAAAAAARGDLRATAMALLALSRVRDGGPLEDALLLLARLVEHVAREGDAEALR LVREALERAVELGLPEEAAAFLRQLAALLAFILEQRDNIELLLLVTSEEELDEAIAKIAEVGKH YMVLVSVGKGTLILIATKDPEATMAALRERLPDAQLEFLTENLYFQARRADLALLEAA |
| PO18s S176A | LAEYLRELAAAAAARGDARATAMALLALSRVRDGELLALAETLVALYLLYADRDADVAARL REHALAAIERVLAEEKSEVARARALVLRLNLERAAEGTVVELVTSEEELDEAIAKIAEVGKHY MVLVSVGKGTLILVAGDRETVEAVAAELRAAFPDAQLEFLTENLYFQARRAALALLEAA |
| PO18s S176D | LAEYLRELAAAAAARGDARATAMALLALSRVRDGELLALAETLVALYLLYADRDADVAARL REHALAAIERVLAEEKSEVARARALVLRLNLERAAEGTVVELVTSEEELDEAIAKIAEVGKHY MVLVSVGKGTLILVAGDRETVEAVAAELRAAFPDAQLEFLTENLYFQARRADLALLEAA |
| PO5s S176A | LAEVLRELAAAAAARGDARATAMALLALSRVRDGGPLEDALLLLARLVEHVAREGDAEALR LVREALERAVELGLPEEAAAFLRQLAALLAFILEQRDNIELLLLVTSEEELDEAIAKIAEVGKH YMVLVSVGKGTLILIATKDPEATMAALRERLPDAQLEFLTENLYFQARRAALALLEAA |
| PO5s S176D | LAEVLRELAAAAAARGDARATAMALLALSRVRDGGPLEDALLLLARLVEHVAREGDAEALR LVREALERAVELGLPEEAAAFLRQLAALLAFILEQRDNIELLLLVTSEEELDEAIAKIAEVGKH YMVLVSVGKGTLILIATKDPEATMAALRERLPDAQLEFLTENLYFQARRADLALLEAA |
| PO5 S131C A183C | LAEVLRELAAAAAARGDLRATAMALLALSRVRDGGPLEDALLLLARLVEHVAREGDAEALR LVREALERAVELGLPEEAAAFLRQLAALLAFILEQRDNIELLLLVTSEEELDEAIAKIAEVGKH YMVLVCVGKGTLILIATKDPEATMAALRERLPDAQLEFLTENLYFQARRASLALLEAC |
| PO5 K134C A183C | LAEVLRELAAAAAARGDLRATAMALLALSRVRDGGPLEDALLLLARLVEHVAREGDAEALR LVREALERAVELGLPEEAAAFLRQLAALLAFILEQRDNIELLLLVTSEEELDEAIAKIAEVGKH YMVLVSVGCGTLILIATKDPEATMAALRERLPDAQLEFLTENLYFQARRASLALLEAC |
| PO5 G135C A183C | LAEVLRELAAAAAARGDLRATAMALLALSRVRDGGPLEDALLLLARLVEHVAREGDAEALR LVREALERAVELGLPEEAAAFLRQLAALLAFILEQRDNIELLLLVTSEEELDEAIAKIAEVGKH YMVLVSVGKCTLILIATKDPEATMAALRERLPDAQLEFLTENLYFQARRASLALLEAC |

**Table S6. Sequences of membrane-targeting variants of obligate and responsive-oligomers for *in vitro* membrane binding.** The introduced MTS sequence is underlined.

| Designs | Sequence |
|---|---|
| PI31_MTS | LHESQQEELLELVLRAAELAGVRVRIRFKGDEIEVELYVEPEIEPEEFTERMKKFFEELKKLAEERGLVEEVKKITFTIVLERNVRPEDCEAIEEVGKEAKKAGLKVKFVVRITGYHIKQQRQLYRDVREAAKKAGVEVEIEVEGSGIEEEKKGFLKRLFGG |
| LP2_MTS | MKLKEVIEEFMEENKDELEDYEILIDEKYYDLDVLEYKIEELEINKEETEEEINEKLNELEEKYEKEGYFLVYIAYLKNNKIELLFVKLDEENQEKLRNDGSGIEEEKKGFLKRLFGG |
| MC11_MTS | KDALTKMAAAAADAWSAADLLQEALGLDKELAHDFRHGFWILIGQIHDALHAGDESALPALEERLEALRERLGELAVERVATAIADPEKFARLLLACARLGAAQFARVVEAVAELLPQLDKDALTKMAAAAADAWSAVDTVASLLSAEEQHDFRHGFWILIGQIHDALHGSGIEEEKKGFLKRLFGGGSGIEEEKKGFLKRLFGG |
| CHD04_MTS | LAGIAEGWLVMERLLRAGADVDEALEAFAEVLRRYGHEHLIEPVRERVREIVAKLRDKLSPFELASLAALSAARLIFEERNDPLAEIIEFTLETLVEILEKGLNEEESVKFIEEKIVEKNAPLLEEAERLFERLLEKLLSKEDKEVLEKHRELMDEYSEKGDIERLASVAATAMLHVYFSLGLNDPKISSWIFEFRRESIRNLEIIPTEELRKLYIRVKMMYELVKILLKVASEEEKKELEKLLELLEKILKSLELILKLREIVEERKKKGIELTPFEIALEAYYVAVEEYLKEGSGIEEEKKGFLKRLFGG |
| LBM10_MTS | KYMLVYRSQKDKNVTIFRWIEKEDLFEEVYRILREEYGEELERQVNLRVSQLLQDFVELYGGEKLPLVASREGVRELFARARPTLRQAGDDFSRRYRRDFAEMSRLIAEEKAKYAEDAEEQERFFRTVVEELFRDGVNWGRIVAFFEFGRLWGLSDEEIFEALLEFIGRQLVKEHLKKNGEIKGNALYLEGSGIEEEKKGFLKRLFGG |
| PO5s_MTS | LAEVLRELAAAAAARGDARATAMALLALSRVRDGGPLEDALLLLARLVEHVAREGDAEALRLVREALERAVELGLPEEAAAFLRQLAALLAFILEQRDNIELLLLVTSEEELDEAIAKIAEVGKHYMVLVSVGKGTLILIATKDPEATMAALRERLPDAQLEFLTENLYFQARRASLALLEAAGSGIEEEKKGFLKRLFGG |

63. A. Hill, A. Hill, A. Paganini-Hill, The possible effects of the aggregation of the molecules of haemoglobin on its dissociation curves. *J. Physiol.*

64. U. Mueller, T. Barthel, L. Benz, V. Bon, T. Crosskey, C. Genter Dieguez, R. Förster, C. Gless, T. Hauß, U. Heinemann, M. Hellmig, D. James, F. Lennartz, M. Oelker, R. Ovsyannikov, P. Singh, M. C. Wahl, G. Weber, M. S. Weiss, The macromolecular

crystallography beamlines of the Helmholtz-Zentrum Berlin at the BESSY II storage ring: history, current status and future directions. *J. Synchrotron Radiat.* **32**, 766–778 (2025).

76. D. Liebschner, P. V. Afonine, M. L. Baker, G. Bunkóczi, V. B. Chen, T. I. Croll, B. Hintze, L.-W. Hung, S. Jain, A. J. McCoy, N. W. Moriarty, R. D. Oeffner, B. K. Poon, M. G.

Prisant, R. J. Read, J. S. Richardson, D. C. Richardson, M. D. Sammito, O. V. Sobolev, D. H. Stockwell, T. C. Terwilliger, A. G. Urzhumtsev, L. L. Videau, C. J. Williams, P. D. Adams, Macromolecular structure determination using X-rays, neutrons and electrons: recent developments in Phenix. *Acta Crystallogr. Sect. Struct. Biol.* **75**, 861–877 (2019).

86. B. Ramm, P. Glock, P. Schwille, In Vitro Reconstitution of Self-Organizing Protein Patterns on Supported Lipid Bilayers. *J. Vis. Exp. JoVE*, e58139 (2018).

87. T. D. Goddard, C. C. Huang, E. C. Meng, E. F. Pettersen, G. S. Couch, J. H. Morris, T. E. Ferrin, UCSF ChimeraX: Meeting modern challenges in visualization and analysis. doi: 10.1002/pro.3235.
